# Prediction of elimination of intrahepatic cccDNA in hepatitis B virus-infected patients by a combination of noninvasive viral markers

**DOI:** 10.1101/2022.11.04.515164

**Authors:** Kwang Su Kim, Masashi Iwamoto, Kosaku Kitagawa, Sanae Hayashi, Senko Tsukuda, Takeshi Matsui, Masanori Atsukawa, Natthaya Chuaypen, Pisit Tangkijvanich, Lena Allweiss, Takara Nishiyama, Naotoshi Nakamura, Yasuhisa Fujita, Eiryo Kawakami, Shinji Nakaoka, Masamichi Muramatsu, Kazuyuki Aihara, Takaji Wakita, Alan S. Perelson, Maura Dandri, Koichi Watashi, Shingo Iwami, Yasuhito Tanaka

## Abstract

Evaluation of intrahepatic covalently closed circular DNA (cccDNA) is a key for searching an elimination of hepatitis B virus (HBV) infection. HBV RNA and HBV core-related antigen have been proposed as surrogate markers for evaluating cccDNA activity, although they do not necessarily estimate the amount of cccDNA. Here, we developed a novel multiscale mathematical model describing intra- and inter-cellular viral propagation, based on the experimental quantification data in both HBV-infected cell culture and humanized mouse models. We applied it to HBV-infected patients under treatment and developed a model which can predict intracellular HBV dynamics only by use of noninvasive extracellular surrogate biomarkers. Importantly, the model prediction of the amount of cccDNA in patients over time was confirmed to be well-correlated with the liver biopsy data. Thus, our noninvasive method enables to predict the amount of cccDNA in patients and contributes to determining the treatment endpoint required for elimination of intrahepatic cccDNA.

## Introduction

Chronic infection with hepatitis B virus (HBV) elevates the risk of developing hepatocellular carcinoma. The WHO has estimated that 297 million people worldwide are living with HBV and that 820,000 people died from this infection in 2019 (https://www.who.int/en/news-room/fact-sheets/detail/hepatitis-b)^1^. Persistence of HBV infection is attributable to the formation of covalently closed circular DNA (cccDNA) in the nucleus of an infected hepatocyte. The cccDNA acts as a reservoir that transcribes HBV RNA and produces HBV DNA through reverse transcription. The cccDNA also drives transcription to produce viral proteins such as HBV surface antigen (HBsAg) and HBV core-related antigen (HBcrAg), comprising HBV core antigen (HBcAg), HBV e antigen (HBeAg) and a 22-kDa truncated core-related protein (p22cr). HBV DNA integrated in a cellular chromosome is an additional source to produce a part of HBV antigens especially HBsAg.

Pegylated interferon alpha (PEG IFN-α) and nucleos(t)ide analogues (NAs) are used for treatment of chronic hepatitis B (CHB). PEG IFN-α activates host immune responses and suppresses viral replication. NAs inhibit the reverse transcription to reduce HBV DNA, which results in the improvement of liver pathology. In most patients, these therapies reduce serum HBV DNA to undetectable level but their effects on HBV antigens such as HBsAg are limited to still show a positivity, which is defined as a *partial cure*. A *functional cure,* that is, undetectable HBV DNA and HBsAg in the serum as well as cccDNA silencing with or without seroconversion, is limited by these therapies^1^, and is a current clinical goal of anti-HBV therapy. A *complete cure*, that is, undetectable HBV DNA and HBsAg in the serum and cccDNA clearance in the liver is the eventual goal for HBV elimination. Because of the difficulty in transcriptional silencing and elimination of cccDNA, patients often require life-long treatment and few maintain sustained viral or clinical remission off therapy^2^.

Quantification of cccDNA amount in a patient’s liver requires a liver biopsy, which is not generally done in clinical practice. Therefore, noninvasive viral markers that reflect the cccDNA in the liver are used for evaluating functional cure. While the level of HBsAg in the serum has been shown to have only a weak or no correlation with cccDNA especially in HBeAg-negative patients as well as HBsAg is produced not from persistent cccDNA transcription but from integrated HBV DNA genomes, there are accumulating reports suggesting that the amounts of HBV RNA and HBcrAg in the serum better reflect the transcriptionally active cccDNA in the liver, since they are not produced by integrated viral DNA. However, since expression of HBV RNA and HBcrAg depends on not only the amount of cccDNA but also the transcriptional activity of cccDNA, which can vary among the patient cohort and other factors such as the disease phase and whether patients are being given antiviral therapy (i.e., huge interindividual variation may be present), they are not necessarily useful for predicting the amount of cccDNA. Thus, lack of a noninvasive method for monitoring the amount of intrahepatic cccDNA is a gap toward evaluation for the status of complete cure.

In this study, we propose a predictive method for quantifying the amount of intrahepatic cccDNA. We developed a multiscale mathematical model that described the HBV propagation process based on the experimental data in cell culture and humanized mice models. Our method uses three viral markers—HBsAg, HBcrAg and HBV DNA—to estimate the amount of intrahepatic cccDNA. We demonstrated that it can be applied to clinical data under treatment in both HBeAg-positive and -negative patient cohorts and confirmed the prediction well-captured the cccDNA level in paired liver biopsy. This noninvasive method predicting the dynamics of intrahepatic cccDNA amount in patients was also shown to propose the endpoint of anti-HBV treatments until elimination of cccDNA.

## Results

### Mathematical model for calculating HBV dynamics in a cell culture model

To develop a mathematical model reflecting the dynamics of HBV propagation including cccDNA, we performed cell culture experiments using primary human hepatocytes (PHH) because cccDNA can be “directly” quantified in this system. PHH were infected with HBV and the amount of extracellular and intracellular HBV DNA and intracellular cccDNA were monitored longitudinally (every three to four days up to 24-31 days post-inoculation) under with or without drug treatment (**Fig. 1**, **Fig. S1**, **Fig. S2** and **ONLINE METHODS**). Note that PHH were maintained at 100% confluent conditions with 2% concentration of dimethyl sulfoxide (DMSO) including medium during the entire infection assay, to supports low cell growth and prevent cell division^3–5^. We developed the mathematical model (**Fig. 1A**) given by Eqs.(S1-S4) in **Supplementary Note 1** and fitted the model to the time-course quantification datasets obtained with and without treatment with entecavir (ETV) (**Supplementary Note 5**). Inhibiting HBV DNA production by ETV perturbates intracellular HBV replication, which enabled us to estimate parameters in the mathematical model^6^. The typical behaviour of the model using these best-fit parameter estimates is shown together with the data in **Fig. 1B**, and the estimated parameters and initial values are listed in **Table S1**. It was estimated that 214 copies of HBV DNA is produced from cccDNA in a cell per day in average; only 0.00126% of the produced HBV DNA is used for recycling back to produce cccDNA (**Table S2**); and the mean half-life of cccDNA is 51 days in PHH (**Table 1**), which is consistent with previous results showing the cccDNA half-life and the limited recycling activity in PHH^4,5,7^.

**Figure 1.**
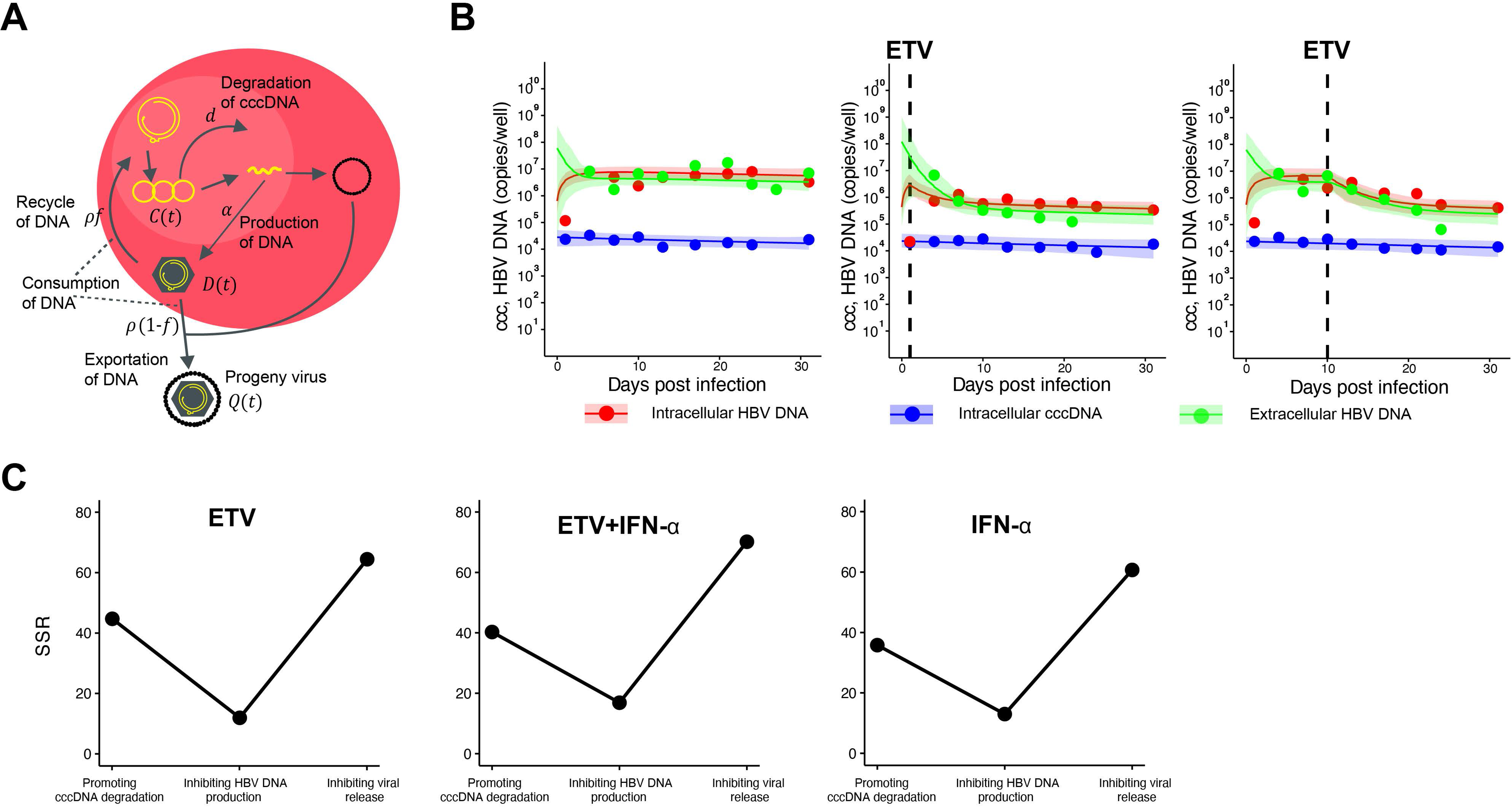
Dynamics of HBV infection in PHH cells: **(A)** Modeling of the intracellular viral life cycle in HBV-infected primary human hepatocytes is shown. Intracellular HBV DNA is produced from cccDNA at rate *α* and is consumed at rate *ρ* That is, a fraction 1 – *f* of HBV DNA assembled with viral proteins as virus particles is exported from infected cells, and the other fraction *f* is reused for further cccDNA formation having a degradation rate of *d*. **(B)** Fits of the mathematical model (solid lines) to the experimental data (filled circles) on intracellular HBV DNA and cccDNA, and extracellular HBV DNA in PHH without treatment, or treated with ETV at different times post-infection (red: intracellular HBV DNA, blue: intracellular cccDNA, green: extracellular HBV DNA). The shadowed regions correspond to 95% posterior intervals and the solid curves give the best-fit solution (mean) for Eqs. (S1-3) to the time-course dataset. All data were fitted simultaneously. **(C)** Sum of squared residuals from best-fits of the mathematical models assuming hypothetical mechanisms of action of ETV and IFN-α.

**Table 1.** Estimated half-life of cccDNA.

| **PHH and Humanized mouse** |  |  |
| --- | --- | --- |
| **Object in data analysis** | **Mean (day)** | **95% CI (day)** |
| PHH | 51 | 14 – 191 |
| Humanized mouse with ETV | 86 | 51 – 170 |
| Humanized mouse with IFN-α | 43 | 33 – 57 |
| **HBV-infected patient** |  |  |
| **Object in data analysis** | **Median (day)** | **Range (day)** |
| NAs (ETV or LAM)-treated patient | 572 | 63 – 2846 |
| Patient without or before PEG IFN-α treatment | 829 | 52 – 6488 |
| - VR of HBeAg positive | 707 | 276 – 3049 |
| - non-VR of HBeAg positive | 985 | 410 – 5429 |
| - PVR of HBeAg negative | 710 | 65 – 4391 |
| - non-VR of HBeAg negative | 804 | 52 – 6488 |
| PEG IFN-α-treated patient for VR of HBeAg positive | 59 | 18 – 332 |
| PEG IFN-α-treated patient for non-VR of HBeAg positive | 198 | 61 – 538 |
| PEG IFN-α-treated patient for PVR of HBeAg negative | 68 | 19 – 425 |
| - monotherapy | 64 | 19 – 425 |
| - combinations with NAs | 100 | 32 – 279 |
| PEG IFN-α-treated patient for non-PVR of HBeAg negative | 221 | 45 – 541 |
| - monotherapy | 251 | 45 – 541 |
| - combinations with NAs | 197 | 55 – 420 |

To address the effect of cytokines on HBV dynamics and predict their possible mechanisms of action, we analyzed the time-course datasets with the mathematical model assuming hypothetical mechanisms of action (Eqs.(S5-S7) in **Supplementary Note 1**). We found that our simple statistical test, that is, calculating the sum of squared residuals (SSR) and selecting a mathematical model with the smallest SSR, could successfully predict the mechanism of action of ETV that inhibit HBV DNA production, rather than facilitate cccDNA degradation or inhibit viral release (**Fig. 1C** and **Fig. S2**). By applying this model, IFN-α was predicted to dominantly target the process for HBV DNA production (**Fig. 1C** and **Table S2**). This is consistent with that IFN-α inhibits the transcription and encapsidation, and promotes viral RNA degradation (that correspond to the “HBV DNA production” in this model)^8–11^. On the other hand, it was difficult to detect subdominant effects on other points of action due to the dominance of HBV DNA inhibition. Thus, by setting the prerequisite that HBV DNA replication can be sufficiently inhibited by IFN-α, we attempted to detect the “subdominant” mechanism of action (e.g., promoting cccDNA degradation as reported^12^) in the following experiments.

### Extended mathematical model captures cccDNA half-life and its decay as induced by anti-HBV drugs in an *in vivo* model

While we can “directly” monitor cccDNA dynamics in hepatocyte cell culture experiments (**Fig. 1**, **Fig. S1** and **Fig. S2**), it is difficult to obtain time-course measurements of cccDNA *in vivo*. We thus extended the above combined experimental-theoretical approach to describe HBV dynamics *in vivo* and to estimate the cccDNA half-life using surrogate biomarkers present in peripheral blood. To check the performance of our extended approach, we first conducted HBV infection experiment with humanized liver uPA/SCID mice: after inoculation with HBV and reaching a sustained HBV DNA load (approximately 5.6 × 10^8^ copies/ml) at 53 days post-inoculation, mice were treated with or without ETV or PEG IFN-α continuously to longitudinally monitor four different biomarkers in the peripheral blood every three to seven days up to 70 days post-treatment: extracellular HBV DNA, HBcrAg, HBeAg and HBsAg (**Fig. 2**, **Fig. S1** and **ONLINE METHODS**).

**Figure 2.**
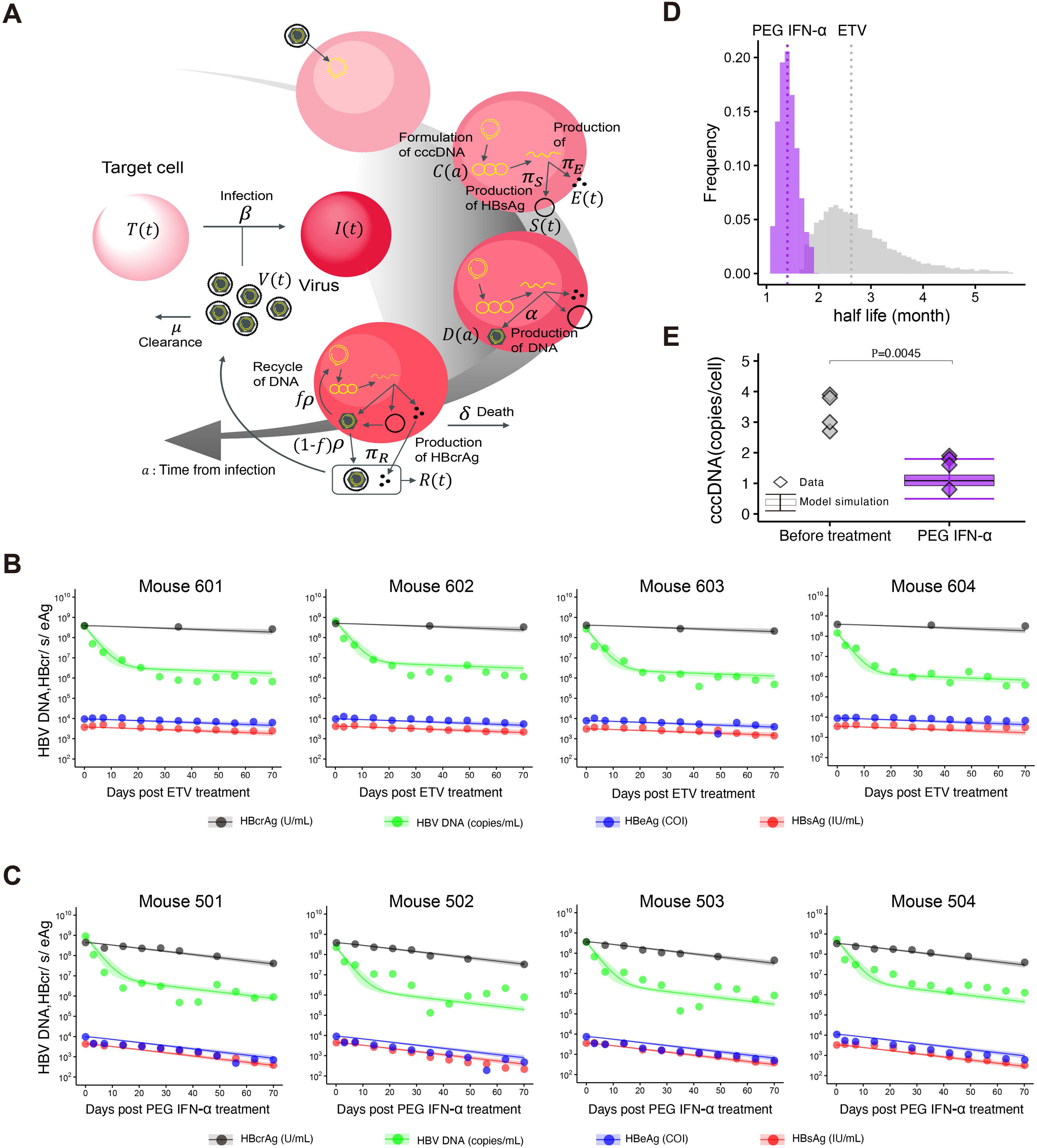
Dynamics of HBV infection in humanized mice: **(A)** Multiscale modeling of intracellular replication and intercellular infection is described. The entry virion forms cccDNA in the nucleus and produces intracellular HBV DNA at rate *α*. HBsAg, HBeAg and HBcrAg antigens are also produced from cccDNA at rates *π_S_, π_E_* and *π_R_* and cleared at *σ_s_, σ_E_* and *σ_R_* in peripheral blood, respectively. The intracellular HBV DNA is consumed at rate *ρ*, of which a fraction 1 – *f* of HBV DNA assembled with viral proteins as virus particles is exported from infected cells and the other fraction *f* is reused for further cccDNA formation having a degradation rate of *d*. The infected cells are dead at rate *γ* and the exported viral particles, which are cleared at rate *μ*, infect their target cells at rate *β*. **(B)** and **(C)** show fits of the mathematical model to the surrogate biomarkers in peripheral blood of humanized mice treated with ETV or PEG IFN-α (black: HBcrAg, green: HBV DNA, blue: HBeAg, red: HBsAg). The shadowed regions correspond to 95% posterior intervals and the solid curves give the best-fit solution (mean) for Eqs. (S34-37) or (S45-48) to the time-course dataset. All data were fitted simultaneously. **(D)** The distribution of the half-life of cccDNA, log2 /*d*, under treatment with PEG IFN-α inferred by MCMC computations. **(E)** Comparisons of predicted cccDNA copies/cell by Eq. (S50) with estimated parameters and the observed cccDNA levels at baseline and 70 days after PEG IFN-α treatment in humanized mice. Black line indicates the median, box and whiskers show the interquartile range (IQR) and 1.5×IQR, respectively.

Here, to precisely quantify both intracellular and extracellular virus dynamics from these biomarkers, we used a multiscale mathematical model of HBV infection combining the intracellular mathematical model (Eqs.(S1-S3)) with the standard virus dynamics model^13,14^, in which an infected cell produces progeny HBVs extracellularly that then are degraded or infect other cells (**Fig. 2A** and Eqs.(S8-S15) in **Supplementary Note 2**). We derived simple linearized equations (Eqs.(S34-S37) in **Supplementary Note 3** and Eqs.(S45-S48) in **Supplementary Note 4**) for fitting to the time-course datasets quantified with mice upon or without ETV or PEG IFN-α treatment (**Table S3**, **Table S4** and **Supplementary Note 5**), and showed that the model well-captured the experimental quantification data over time with best fit parameters (**Fig. 2BC**). Note that the decay rates of infected cells were estimated separately from human albumin in peripheral blood of humanized mice (**Fig. S3**) and the clearance rates of extracellular HBV DNA and antigens were fixed as previously estimated values, that is, *μ* = 16.1 d^-1 15^ and *σ* = 1.00 d^-1 16^.

When we applied the mathematical model to the evaluation of the drug effects on viral replication and amount of cccDNA, it is assumed that ETV almost completely blocks intracellular HBV replications and *de novo* infections but has no direct effect on the cccDNA degradation, as reported previously (**Supplementary Note 3**)^17–20^. Interestingly, we found the mean half-life of cccDNA was 86 days in the humanized mice under ETV treatment (**Fig. 2D** and **Table 1**). In addition to the potent antiviral effect of PEG IFN-α as observed in HBV infection of PHH (**Fig. 1C**) and other reports^21^, our analysis demonstrated that PEG IFN-α treatment significantly reduces the half-life of cccDNA to around 43 days (**Fig. 2D** and **Table 1**). This calculation is supported by our previous mouse experiments that PEG IFN-α treatment for 42 days reduced cccDNA levels to 23-33%, which was semi-quantified with the bands detected by southern blot^22^ (**Table S5**). Note that this cccDNA half-life upon PEG IFN-α treatment is estimated under the assumption that no *de novo* infections occurs due to the robust antiviral effects of PEG IFN-α; the cccDNA half-life value can be even shorter when a low level *de novo* infections occurs upon PEG IFN-α treatment (**Supplementary Note 4**).

Importantly, the intrahepatic cccDNA levels experimentally measured in the liver that was collected from the humanized mice (cccDNA was measured by collecting the liver from sacrificed mice, and digested with plasmid-safe adenosine triphosphate dependent deoxyribonuclease DNase (PSAD), followed by absolute quantification by droplet digital PCR (ddPCR))^22,23^ were confirmed to be within the range of values calculated by our mathematical model (**Fig. 2E**). Taken together, our extended approach with surrogate biomarkers in peripheral blood predicted intrahepatic cccDNA dynamics and captured the reduction of the half-life of cccDNA in *vivo* by treatment with PEG IFN-α.

### Combination of a mathematical model and noninvasive viral markers can predict the amount of intrahepatic cccDNA in chronically HBV-infected patients

We extended our mathematical model-based analysis to clinical datasets to address the amount of cccDNA. We analyzed CHB cohorts comprising a total of 226 patients in three Japanese and one Thailand hospitals among who 199 patients were treated with PEG IFN-α monotherapy or PEG IFN-α combination with NAs (ETV or lamivudine (LAM)) for 48 weeks and 27 patients received NAs. Serum from these patients were collected from the start of treatment (day 0) to end of treatment (48 weeks) to detect HBcrAg, HBV DNA, and HBsAg (**Fig. S1C**). We separated the patients into four groups according to their HBeAg status and their eventual virological response to treatment. *Virological response* (VR) was defined as HBeAg clearance and HBV DNA level <2,000 IU/ml at 48 weeks after treatment in HBeAg-positive CHB. *Persistent VR* (PVR) was defined as HBeAg clearance and HBV DNA level <2,000 IU/ml at 96 weeks after treatment in HBeAg-negative CHB. Non-VR and non-PVR were those who did not reach the criteria for VR and PVR, respectively. We analyzed the following longitudinally monitored biomarkers in peripheral blood^24,25^: extracellular HBV DNA, HBcrAg and HBsAg for up to 48 weeks after starting treatment (**Fig. 3**, **Fig. S1**, **Fig. S4** and **ONLINE METHODS**). We also used the derived linearized model equations under the assumption of negligible *de novo* infections under treatment, as did in the earlier mouse infection analysis (Eqs. (S45-S46)(S48) in **Supplementary Note 4**)^18,19,26–28^. All biomarkers of all patients were simultaneously fitted using a nonlinear mixed-effect modeling approach (**Supplementary Note 5**), which uses samples to estimate population parameters while accounting for inter-individual variation (**Fig. S4**, **Table S6** and **Table S7**).

**Figure 3.**
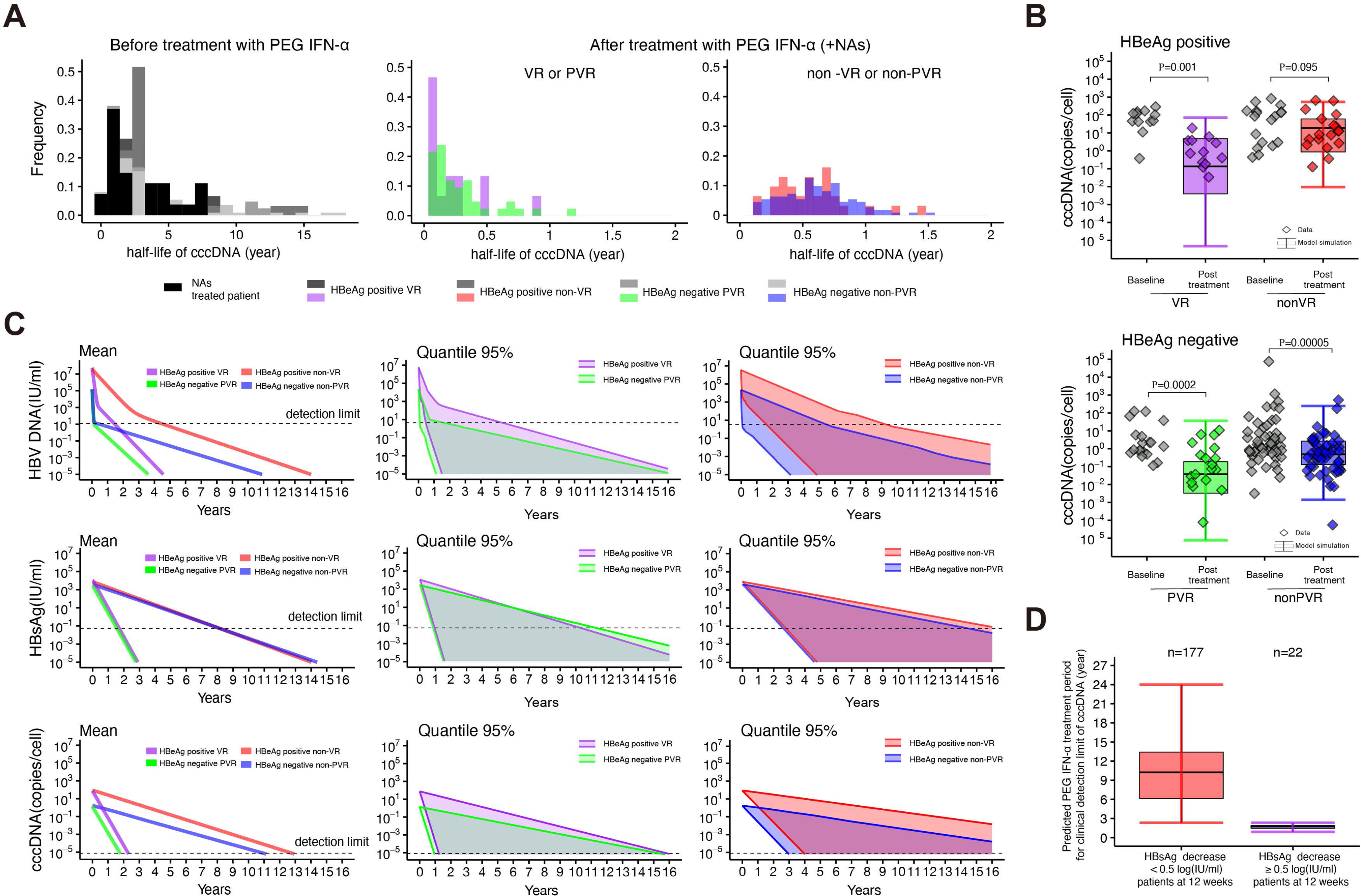
Dynamics of HBV infection in patients treated with PEG IFN-α: **(A)** The distributions of the half-life of cccDNA before and after treatment with PEG IFN-α for HBeAg-positive/negative and (P)VR/non-(P)VR patients are shown. **(B)** Comparisons of predicted cccDNA per cell from Eq. (S50) with estimated parameters and the observed cccDNA at baseline and at 48 weeks after treatment in hepatocytes of HBeAg-positive/negative and (P)VR/non-(P)VR patients treated with PEG IFN-α. **(C)** Predicted dynamics of HBV DNA, HBsAg and cccDNA under a hypothetical long PEG IFN-α treatment are calculated. The solid lines in the left panels give the mean of Eqs. (S45-S46)(S50) with estimated parameters, and the shadowed regions in the middle and right panels correspond to 95% predictive intervals for HBeAg-positive/negative and (P)VR/non-(P)VR patients. The horizontal dashed lines in HBV DNA, HBsAg and cccDNA show the detection limits. **(D)** Predicted PEG IFN-α treatment period needed to drive the cccDNA level below the detection limit for patients stratified on the basis of HBsAg reduction at 12 weeks after treatment (red: less than 0.5 log_10_ (IU/ml), purple: greater than 0.5 log_10_ (IU/ml)). Black line indicates the median; box and whiskers show the interquartile range (IQR) and 1.5×IQR, respectively.

The model predicted that the decay rate of cccDNA varies among patients (**Table S7**) showing a median half-life of cccDNA of around 2.3 years in the patients without (or before) PEG IFN-α treatment, and no significant difference in half-life among the four groups of patients, before treatment: HBeAg-positive/negative at baseline and PVR/non-PVR patients (707, 985, 710, and 804 days) (**Fig. 3A** and **Table 1**). Interestingly, PEG IFN-α significantly decreased the cccDNA half-life in all patients regardless of combination with NAs (**Fig. 3A** and **Table 1**): the median values in patients achieving VR and PVR were 59 days (range, 18-332 days) and 68 days (range, 19-425 days) in HBeAg-positive and HBeAg-negative patients, respectively, and for non-VR and -PVR patient groups it was 198 days (range, 61-538 days) and 221 days (range, 45-541 days) for HBeAg-positive and HBeAg-negative patients, respectively. There were significant differences in the half-life between patients achieving (P)VR and non-(P)VR patients (*p* < 0.01 by Mann-Whitney U tests). The estimated half-lives of cccDNA in different sub-groups of patients were summarized in **Table 1**.

The amount of cccDNA in patients before treatment is quantified as median 3.0 (CI 95% 0.1-683.6) copies/cell, which is close to previous reports (**Fig. 3B**)^29–32^. Significant differences in the amount of cccDNA at the beginning of treatment were also observed between HBeAg-positive and HBeAg-negative patients (*p* < 0.01 by Mann-Whitney U tests) (**Fig. 3B**), but the cccDNA half-life was not significantly different (**Table 1**). Note that no significant differences were observed in the half-life of cccDNA after PEG IFN-α treatment according to *CC* or *CT* genotype on the IL28B SNP (**Fig. S6**). We next examined the validity of the estimates of the half-life of cccDNA decay under PEG IFN-α treatment calculated by Eq. (S50) in **Supplementary Note 4** by using paired-liver biopsy samples (pre-treatment, and at 48 weeks end of PEG IFN-α treatment). Experimental measurement of cccDNA (used the PSAD-treated liver samples) shows that the amounts of cccDNA were significantly reduced in the VR (HBeAg-positive) and PVR (HBeAg-negative) patients for PEG IFN-α while those in non-VR and non-PVR showed a minimal decrease (**Fig. 3B**). In fact, the decay rates of cccDNA for all the four cohorts (VR, non-VR, PVR, non-PVR) measured were within the range of values calculated by our mathematical model (**Fig. 3B**), indicating that our mathematical model captured the decay of cccDNA in both the HBeAg-positive and HBeAg-negative cohorts. These results demonstrate that our extended approach constructed on the basis of experimental data can be applied to the prediction of intrahepatic cccDNA.

### Calculation of effectiveness for cccDNA elimination

Estimation of the turnover of intrahepatic cccDNA would be important for the evaluation and design of treatment for cccDNA clearance. The liver biopsy data indicate that PEG IFN-α reduced the amount of cccDNA but is difficult to eliminate cccDNA within 48 weeks of treatment (**Fig. 3B**), consistent with previous reports that PEG IFN-α can potentially target and reduce cccDNA, but the clinical effects of cccDNA clearance is seen in only a minority of CHB patients^33,34^. Given the clear reduction in cccDNA amount especially in VR- and PVR-patients observed in the liver biopsy and the accelerating cccDNA decay shown by our model (2.3 years to 59-221 days as half-life), 48 weeks of PEG IFN-α treatment may not be sufficient but prolonged treatment may be beneficial to eliminate cccDNA. Aiming to design a better treatment for cccDNA clearance, we thus calculated the duration of PEG IFN-α treatment needed to achieve negativity for cccDNA as well as HBV DNA and HBsAg by using the mathematical model with our best-fit estimated parameters.

First, we defined the eradication boundary for each biomarker; under 12 (IU/ml) for HBV DNA^35,36^, 0.05 (IU/ml) for HBsAg^37–39^, and 0.8 × 10^-5^ (copies/cell)^23,40^ for cccDNA as described previously, and defined values below these thresholds as achieving negativity (**Table 2** and **Supplementary Note 6**). We then simulated HBV DNA, HBsAg and cccDNA dynamics using Eqs. (S45-S46)(S50) in **Supplementary Note 5** with the estimated individual parameters for each group of patients and the initial conditions for all patient (**Table S7**). The predicted dynamics of HBV DNA, HBsAg and cccDNA with 95% predicted intervals for HBeAg-positive/negative and (P)VR/non-(P)VR patients under a hypothetical long PEG IFN-α treatment are calculated in **Fig. 3C**. Our *in silico* simulations estimated that the periods required for HBsAg clearance by PEG IFN-α are longer than those for HBV DNA clearance, and those for cccDNA clearance are further longer in patients of all the four groups, which is consisted with the clinical observations (**Table 2**)^41^. To achieve HBV DNA clearance, HBeAg-positive patients also require a longer period of PEG IFN-α treatment than do HBeAg-negative patients regardless of VR status (**Table 2**). On average, treatment with PEG IFN-α for more than 10 years is required to eradicate cccDNA in patients who are non-(P)VR regardless of HBeAg status. The mean treatment periods of HBeAg-positive patients for cccDNA clearance are 2.3 years (95% CI, 1.2-15.9 years) and 12.7 years (95% CI, 4.0-29.8 years) for VR and non-VR patients, respectively (**Table 2**).

**Table 2.** Predicted PEG IFN-α treatment periods needed to reach the detection limit for HBV DNA, HBsAg and cccDNA/cell.

| Type of biomarker | HBeAg-positive VR | HBeAg-positive non-VR | HBeAg-negative PVR | HBeAg-negative non-PVR |
| --- | --- | --- | --- | --- |
| HBV DNA (IU/ml) | 1.0<sup>†</sup> (0.5 – 5.4)<sup>‡</sup>years | 4.9 (1.6 – 9.3)years | 0.4 (0.1 – 1.7) years | 0.5 (0.1 – 4.4) years |
| HBsAg (IU/ml) | 1.7 (0.9 – 10.3)years | 8.2 (2.8 – 17)years | 1.6 (0.9 – 11.3)years | 8.5 (2.6 – 15)years |
| cccDNA (copies/cell) | 2.3 (1.2 – 15.9)years | 12.7 (4.0 – 29.8)years | 1.8 (0.9 – 15.4) years | 10.8 (2.9 – 21.2) years |

Assumed detection limits are 12(IU/ml)<sup>35,36</sup>, 0.05(IU/ml)<sup>37-39</sup>, and $0.8 \times 10^{-5}$ (copies/cell)<sup>40</sup> for HBV DNA, HBsAg, and cccDNA, respectively.
<sup>†</sup> Mean value
<sup>‡</sup> 95% confidence interval

By simulating HBsAg and cccDNA dynamics using estimated individual parameters in 199 patients who received PEG IFN-α, we calculated the required period of PEG IFN-α treatment to achieve cccDNA negative for patients stratified on the basis of HBsAg reduction at 12 weeks after treatment^42^ (**Fig. 3D**). If the reduction in HBsAg was less than 0.5 log_10_ (IU/ml)^43^, our simulation predicted that a median of 10.3 years of PEG IFN-α treatments (IQR, 6 to 13.9 years) are needed to eliminate cccDNA. On the other hand, if the HBsAg reduction exceeded 0.5 log10 (IU/ml), the period of treatment for cccDNA clearance is predicted to be 1.7 years (IQR, 1.5 to 1.9 years). This simulation could be applied to determine an appropriate treatment period on demand. Since cccDNA clearance from the liver is the final goal of antiviral treatment in CHB^42^, our approach is potentially useful for the optimal design of response-guided treatment with PEG IFN-α.

## Discussion

HBcrAg and HBV RNA have been proposed as surrogate markers for the transcriptionally active cccDNA^44–49^ and have been used to evaluate the antiviral effect of drugs to functional cure. Recent clinical studies for new anti-HBV candidates such as HBV capsid inhibitors or small interfering RNAs (siRNAs) measured HBV RNA as well as HBV DNA and viral antigens as biomarkers^50,51,52^ and suggests their effect on the cccDNA activity to discuss the drug potential for achieving a functional cure. However, these markers do not necessarily correlate with the amount of cccDNA since their values are also reflected by the transcriptional activity of cccDNA. A previous study estimates the turnover of cccDNA by monitoring the signature mutation (M204I/V) induced by lamivudine treatment in HBV RNA in the serum^7^. While this method is an innovative proposal, it is unclear whether the method will be useful in estimating the cccDNA amount and turnover in patients under PEG IFN-α therapy without the signature mutation. It is a significant challenge to develop a noninvasive method that estimates the amount and turnover of cccDNA for searching and arguing a complete cure.

Here, we developed a multiscale mathematical model for quantifying HBV viral dynamics based on *in vitro* and *in vivo* experimental data and applied this model to the analysis of CHB patients. The amount of intrahepatic cccDNA and its dynamics are predicted by quantification of three serum viral biomarkers—HBV DNA, HBsAg and HBcrAg—in this multiscale model. The estimated half-life and reduction of intrahepatic cccDNA in PEG IFN-α treated patients were supported by clinical datasets including paired liver biopsy data for HBeAg-negative and HBeAg-positive cohorts. Our modeling approach is a noninvasive method that allows the time-course estimation of the amount of cccDNA in CHB patients undergoing treatment and predicting the appropriate duration of therapy for cccDNA clearance (**Fig. 3C-D**).

It is clear from previous studies that 48 weeks of PEG IFN-α treatment is effective for eliminating HBV DNA, and HBsAg in some cases, but not sufficient to eliminate cccDNA^53–55^, which are also supported by our calculations (**Table 2** and **Fig. 3C**). We also propose in this study that prolonged PEG-IFN treatment is effective for improving cccDNA elimination: In our simulations, extending the treatment by 40 weeks (to a total of 88 weeks, or 1.7 years) showed a higher rate of cccDNA elimination in both HBeAg-positive and HBeAg-negative patients whose HBsAg decreased more than 0.5 log10 (IU/ml) at 12 weeks (the right panel in **Fig. 3D**). Previous trials of extended-duration PEG IFN-α treatment in HBeAg-negative patients with poor IFN response^56^ achieved significantly better VR and HBsAg loss^57–60^, although extended PEG IFN-α treatment did not necessarily improve viral elimination in all the patients. According to our calculation, actually, in CHB patients whose HBsAg did not decrease by more than 0.5 log_10_ (IU/ml) at week 12 after PEG IFN-α treatment, the benefit for improving the cccDNA elimination with extending the treatment period will be low. If the treatment period were extended for 6 years, we calculated that the probability of cccDNA elimination would be 23% (the left panel in **Fig. 3D**). However, such a long treatment period may not be realistic from the viewpoint of adverse effects. The validity of our estimation needs to be verified in the future, it is because little paper had quantified the cccDNA under anti-HBV drugs.

Clinical guidelines on the management of HBV infection in EU, USA and Japan specify a duration of PEG IFN-α treatment of 48 weeks. However, if evidence accumulates that extending the treatment duration increases the rate of achieving elimination of intrahepatic cccDNA, the benefit of extending treatment may outweigh the adverse effects. Our approach could also be helpful in predicting response to PEG IFN-α in terms of the adverse effects and cost-effectiveness of treatment. For example, treatment could be extended only in patients who display better sensitivity to PEG IFN-α and/or in patients who could discontinue drugs without risk of viral reactivation. Thus, our multiscale mathematical model may be more helpful in determining the duration of treatment in the future.

The limitation of our study is the experimental quantification method of cccDNA: We quantified cccDNA by PCR-based methods, because of the requirement of large number of quantifications for the mathematical model. Standardization of the detection method for cccDNA by real time PCR has been discussed over the years^22,23^. We have to be careful about the possible overestimation of cccDNA amount even if minimizing the contamination of rcDNA by PSAD digestion as used in this study. However, the cccDNA half-life value estimated by our method is roughly unaffected by a slight shift of cccDNA levels. We minimized this limitation by comparing the PCR-based cccDNA quantification data with the values detected by southern blot in HBV-infected chimeric mice (**Fig. 2D, Table 1, and Table S5**). There are also a few assumptions in our mathematical model underlying the intra- and inter-cellular HBV propagation. We assumed the negligible *de novo* infections under ETV and PEG-IFN treatment, that is, NAs and PEG-IFN inhibit HBV replication by around 100% (i.e., *ε* ≈ 1) (**Supplementary Note 3**). These assumptions may overestimate the mean half-life of cccDNA. After additional datasets on the time-course of the biomarkers with different intensities of NAs and PEG-IFN treatments become available, more precisely the inhibition rate, *ε*, will be determined and our estimation will be improved. Another assumption is that the cccDNA degradation rate under PEG-IFN treatment, *d_IFN_*, includes different immune responsiveness that may develop during the treatment and also affect kinetics of clearance or alter cccDNA activity without clearance. However, the clearance mechanisms accompanying PEG-IFN treatment in our mathematical model may be too simplified for the “all-in-one” cccDNA degradation, since there have been cases in which seroconversion of viral markers has been observed after completion of PEG-IFN treatment ^38,43^. This is presumed to be induced as a result of cccDNA degradation based on PEG-IFN, but it is thought to be achieved by a more complex pathway involving immune cells rather than direct cccDNA degradation by PEG-IFN, which is still an unclear mechanism. Quantitative (and time-dependent) mechanism of PEG-IFN that alters intracellular HBV replication is necessary to improve our mathematical modeling in which variations due to the different immune responsiveness are taken into account. Although current simple but quantitative mathematical model successfully predicts the amount of cccDNA in patients from our noninvasive extracellular surrogate biomarkers, more precise mathematical modeling that improves these limitations will be beneficial for further designing current and future available CHB treatments.

In summary, our multiscale mathematical model combined with an individual patient’s extracellular surrogate viral biomarkers, HBsAg, HBcrAg and HBV DNA, predicts the amount of intrahepatic cccDNA and opens new avenues to design a therapeutic strategy achieving a complete HBV cure.

## METHODS

### Study design

The objective of this study was to establish a multiscale mathematical model for quantifying intrahepatic cccDNA with a noninvasive method, which is based on the results of cell culture and mouse experiment, and it will apply the quantification of amount of intrahepatic cccDNA in CHB patient and estimate the effect of anti-HBV drugs on cccDNA half-life. HBV infection assaies using cell culture and mouse models were performed as a single-center and open-labeled study at National institute of infectious diseases and Phoenix Bio Co., Ltd. (Hiroshima, Japan), respectively. All viral markers obtained from these experiments were quantified, and each quantification method is described in detail in the following sections. As cell culture infection assay, PHH (n=3) isolated from humanized mouse were used to evaluate the effect of treatment with ETV, interferon alpha (IFNα), and ETV + IFNα compared to no-treatment (control group) samples. For mouse experiment, severe combined immunodeficiency mice (n=4) transgenic for the urokinase-type plasminogen activator gene (cDNA-uPA^wild/+^/SCID^+/+^ mice), with their livers replaced by human hepatocytes, were infected with HBV. When HBV levels in the serum reached a plateau after day 53 of infection, mice were treated with ETV or PEG IFN-α and viral markers in the serum and liver were quantified. All efforts were made during the study to minimize animal suffering and to reduce the number of animals used in the experiments. In these experiments, sample size was selected based on previous literature and previous experience.

The novel multiscale mathematical model describing intracellular viral propagation, which is based on the above experimental quantification data, was applied to HBV-infected patients to predict the intracellular HBV dynamics. The CHB patient samples in this study were enrolled totally 226 patients at the Nagoya City University Hospital, Teine-Keijinkai Hospital and Nippon Medical School Chibahokusoh Hospital in Japan and the King Chulalongkorn Memorial Hospital, Bangkok, in Thailand. They were classified into two clinical groups: (i) 199 CHB patients receiving PEG IFN-α monotherapy or PEG IFN-α combination with NAs, which include 46 HBeAg-positive patients and 94 HBeAg-negative patients treated with PEG IFN-α alone and 59 HBeAg-negative patients treated with PEG-IFN-α and ETV combination and (ii) 27 patients receiving NAs (control group). Patients coinfected with HCV and/or human immunodeficiency virus (HIV) were excluded. They were not performed blinded. The study size was determined by the number of samples that were obtained from the cohort study and not based on any power calculations. Written informed consent was obtained from each patient and the study protocol conformed to the ethical guidelines of the Declaration of Helsinki and was approved by the appropriate institutional ethics review committees of each institute.

### HBV infection in primary human hepatocytes

PHH used for the HBV infection assay were maintained according to the manufacturer’s protocol (Phoenix Bio Co., Ltd, Hiroshima, Japan). HBV (genotypeD) used as the inoculum was recovered from the culture supernatant of Hep38.7-Tet cells cultured under tetracycline depletion and concentrated up to 200-fold by polyethylene glycol concentration^61^. PHH were seeded into 96-well plate at 7×10^4^ cells/well and were inoculated with HBV at 8,000 genome equivalents (GEq)/cell in the presence of 4% polyethylene glycol 8,000 (PEG8000) for 16 h. After washing out free HBV, PHH were continuously treated with ETV at 1 μM, interferon alpha (IFNα) at 1,000 IU/ml, ETV at 1 μM + IFNα at 1000 IU/ml or without treatment (control). Cell division is known to reduce the cccDNA per cell in HBV-infected cells^4^; therefore, to avoid this, we maintained PHH at 100% confluent conditions during the entire infection assay. Moreover, a high concentration of dimethyl sulfoxide (DMSO) was included in the culture medium as described previously^62^, which does not allow cell growth and prevents cccDNA loss by cell division^3–5^. Since the experiments using PHH were conducted under the above conditions, cell growth dynamics were ignored in our analysis. The culture supernatant from HBV-infected cells and the cells were recovered to quantify HBV DNA in the culture supernatant, total HBV DNA in the cells and cccDNA by real-time PCR. For real-time PCR, the primer-probe sets used in this study were 5’-AAGGTAGGAGCTGAGCATTCG-3’, 5’-AGGCGGATTTGCTGGCAAAG-3’ and 5’-FAM-AGCCCTCAGGCTCAGGGCATAC-TAMRA-3’ for detecting HBV DNA and 5’-CGTCTGTGCCTTCTCATCTGC-3’, 5’-GCACAGCTTGGAGGCTTGAA-3’ and 5’-CTGTAGGCATAAATTGGT(MGB)-3’ for cccDNA^61^.

### HBV infection of humanized mouse

Humanized mouse were purchased from Phoenix Bio Co., Ltd. (Hiroshima, Japan). The animal protocol was approved by the Ethics Committees of Phoenix Bio Co., Ltd (Permit Number:2200). These mice were infected with HBV at 1.0 × 10^6^ copies/mouse that was obtained from human hepatocyte chimeric mice previously infected with genotype C2/Ce, as described previously^63^. Day 53 after inoculation, HBV-infected mice, which showed a plateau HBV levels in serum, were treated with ETV (at a dose of 0.02 mg/kg, once a day) or PEG IFN-α (at a dose of 0.03 mg/kg, twice a week) continuously for over 70 days (**Fig. 2BC** and **Fig. S1B**). The human albumin level in the serum was measured as described previously^64^. The HBV DNA titer was measured by real-time PCR as previously described^65^. HBsAg, HBcrAg and HBeAg were measured by chemiluminescent enzyme immunoassay using a commercial assay kit (Fujirebio Inc., Tokyo, Japan). The detection limit of the HBsAg assay and HBcrAg assay were 0.005 IU/ml and 1.0 kU/ml, respectively. The cut-off index (COI) of the HBeAg was <1.00 (**Fig. 2BC** and **Fig. S3**). Intrahepatic HBV cccDNA was extracted from a dissected liver treated with PSAD to digest genomic DNA and rcDNA as described previously^66^ (**Fig. 2E**). Genomic DNA was isolated from the livers of chimeric mice using the phenol/chloroform method as previously described^67^. The cccDNA-specific primer-probe set for cccDNA amplification was used for ddPCR assay^66^. After the generation of reaction droplets, intrahepatic cccDNA was amplified using a C1000 touch™ Thermal Cycler (Bio-Rad, Hercules, California, USA). In all cases, intrahepatic cccDNA values were normalized by the cell number measured by the hRPP30 copy number variation assay (Bio-Rad, Pleasanton, California, USA)^68^. Of note, hRPP30 levels were separately determined using DNA that was not treated with PSAD. Group means of the difference in cccDNA/hepatocyte were compared by unpaired t-test.

### PEG IFN-α and NAs-treated HBV patients

The data obtained from a total of 226 patients with CHB classified into two clinical groups: (i) treatment with PEG IFN-α monotherapy or PEG IFN-α combination with NAs and (ii) patients receiving only NAs which defined as control group in this study was used (**Fig. 3A**, **Fig. S1C** and **Fig. S4A-E**).

These 199 patients (i) were treated with PEG IFN-α (180 μg/week) alone or ETV (0.5 mg/day) for 48 weeks and followed up for a minimum of 24 weeks after therapy. Of these 199 patients, the 46 patients with HBeAg-positive CHB were seropositive for HBsAg and HBeAg for at least 6 months before therapy and the other 153 patients with HBeAg-negative CHB were seropositive for HBsAg for at least 6 months, negative for HBeAg and positive for anti-HBe antibody. These 27 patients (ii) were treated with ETV (0.5 or 1mg/day) or LAM (100 mg/day) continuously. Of these 27 patients, 15 patients with HBeAg-positive CHB were seropositive for HBsAg and HBeAg at study entry and the other 12 patients with HBeAg-negative CHB were seropositive for HBsAg at study entry, negative for HBeAg and positive for anti-HBe antibody. *VR* was defined as HBeAg clearance and HBV DNA level <2,000 IU/ml at 48 weeks after treatment in HBeAg-positive CHB. *PVR* was defined as HBeAg clearance and HBV DNA level <2,000 IU/ml at 96 weeks after treatment in HBeAg-negative CHB.

Qualitative HBsAg, HBeAg and anti-HBe in sera were measured by commercially available enzyme-linked immunosorbent assay kits (Abbott Laboratories, Chicago, IL, USA). HBsAg titers were quantified by use of Elecsys HBsAg II Quant reagent kits (Roche Diagnostics, Indianapolis, IN, USA). HBV DNA levels were quantified by use of the Abbott RealTime HBV assay (Abbott Laboratories, Chicago, IL, USA). The lower limit of detection of serum HBV DNA is 10 IU/ml. HBcrAg was measured by chemiluminescent enzyme immunoassay using a commercial assay kit (Fujirebio Inc., Tokyo, Japan). Paired liver biopsies were performed before and at the end of PEG IFN-α treatment for intrahepatic cccDNA analysis (week 0 and 48). After treatment with PSAD to digest linear genomic DNA and relaxed circular HBV DNA, intrahepatic cccDNA was determined by real-time PCR as described previously^24^. The beta-globin gene was used as an internal control and normalized for human genomic DNA in terms of copies/cell. Quantification of beta-globin was performed by a commercially available human genomic DNA kit (The LightCycler Control Kit DNA, Roche Diagnostics, Basel, Switzerland)^69^.

### Statistical analysis

Mathematical modeling, transformation to reduced model and its linearization, data fitting and parameter estimations are described in **Supplementary Note 1-6** in detailed. All analyses of samples were conducted using custom script in R and visualized using RStudio. For comparisons between groups, Mann-Whitney U tests were used. All tests were declared significant for *p* < 0.01.

Additional methods are described in Supplementary Information.

## AUTHOR CONTRIBUTIONS

KW, SI and YT designed the research. MI, SH, ST, LA, MD, KW and YT conducted the experiments. KSK, KK, SN, ASP and SI carried out the computational analysis. KW, SI, and YT supervised the project. All authors contributed to writing the manuscript.

## ACKNOWLEDGMENTS

This study was supported in part by a Grant-in-Aid for JSPS Research Fellows 20J00868 (to M.I.), 21K15453 (to M.I.); Scientific Research (KAKENHI) B 18H01139 (to S.I.), 16H04845 (to S.I.), 20H03499 (to K.W.), 21H02449 (to K.W.); Scientific Research in Innovative Areas 20H05042 (to S.I.); the Ministry of Education, Culture, Sports, Science, and Technology, 20K16996 (to S.H.); AMED Strategic International Brain Science Research Promotion Program 22wm0425011s0302 (to K.A.); AMED JP22dm0307009 (to K.A.); AMED CREST 19gm1310002 (to S.I.); AMED Development of Vaccines for the Novel Coronavirus Disease, 21nf0101638s0201 (to S.I.); AMED Japan Program for Infectious Diseases Research and Infrastructure, 22wm0325007s8002 (to S.H.), 22wm0325007j0103 (to K.W.), 22wm0325007h0001 (to S.I.), 22wm0325004s0201 (to S.I.), 22wm0325012s0301 (to S.I.), 22wm0325015s0301 (to S.I.); AMED Research Program on Emerging and Re-emerging Infectious Diseases 21wm0325007s8002 (to S.H.), 22fk0108140s0802 (to S.I.); AMED Research Program on HIV/AIDS 22fk0410052s0401 (to S.I.); AMED Program for Basic and Clinical Research on Hepatitis 22fk0210094 (to S.I.); AMED Program on the Innovative Development and the Application of New Drugs for Hepatitis B 22fk0310504j0001 (to K.W.), 22fk0310504h0501 (to S.I.); AMED International Collaborative Research Program Strategic International Collaborative Research Program (SICORP) 22jm0210068j0004 (to K.W.); AMED Research Program on Hepatitis 19fk0210036h0502 (to S.I.), 19fk0210036j0002 (to K.W.), 19fk0310114h0103 (to S.I.), 19fk0310114j0003 (to K.W.), 19fk0310101j1003 (to K.W.), 19fk0310103j0203 (to K.W.), JP21fk0310101 (to Y.T.); JST MIRAI JPMJMI22G1 (to S.I. and K.W.); Moonshot R&D JPMJMS2021 (to K.A. and S.I.) and JPMJMS2025 (to S.I.); The National Research Foundation of Korea (NRF) grant funded by the Korea government (MSIT) (2022R1C1C2003637) (to K.S.K.); National Institutes of Health grants R01-OD011095, R01-AI078881, R01-AI116868 and R01-AI028433 (to A.S.P); Smoking Research Foundation (to K.W.); The Takeda Science Foundation (to K.W.); Taiju Life Social Welfare Foundation (to K.W.); Shin-Nihon of Advanced Medical Research (to S.I.); SECOM Science and Technology Foundation (to S.I.); The Japan Prize Foundation (to S.I.).

## Supplementary Information

**Figure S1.**
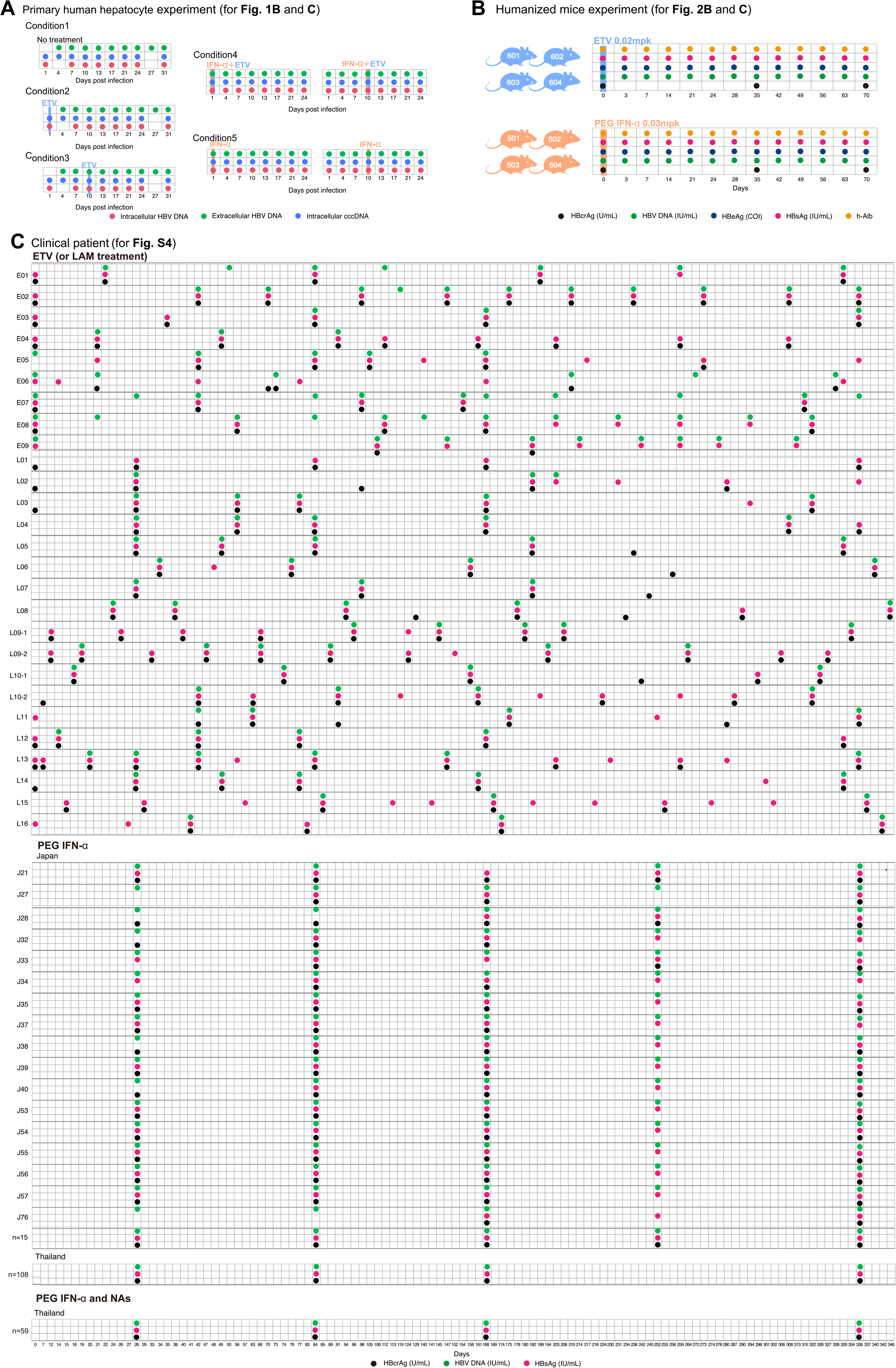
Summary of HBV infection datasets: Detailed data-sampling schedule for HBV-infected **(A)** primary human hepatocytes, **(B)** humanized mice and **(C)** clinical patients.

**Figure S2.**
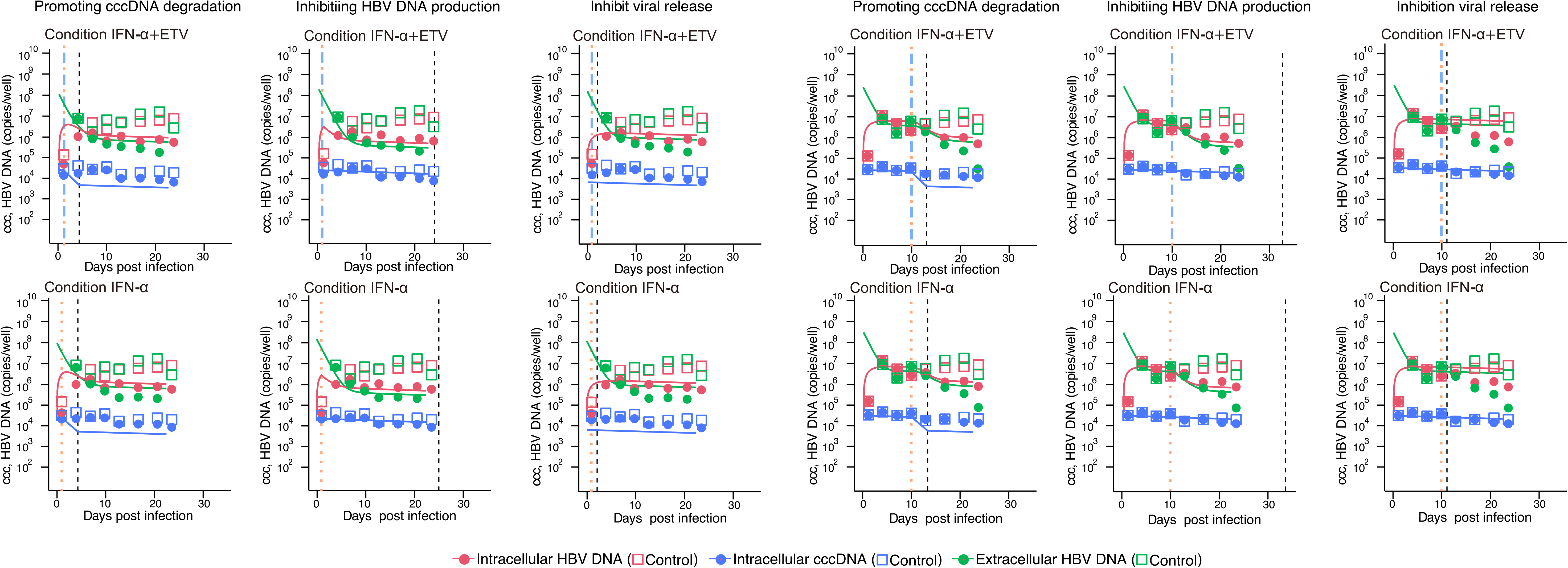
*In silico* experiments to evaluate the antiviral effect of cytokines: Decay characteristics of intracellular cccDNA, intracellular HBV DNA, and extracellular HBV DNA in primary human hepatocytes with antiviral agents are predicted by mathematical models assuming hypothetical mechanisms of action of cytokines. The closed dots (with cytokines), the empty squares (without cytokines), and the solid curves correspond to the observed and estimated intracellular HBV DNA (red), intracellular cccDNA (blue), and extracellular HBV DNA (blue). The colored and black vertical lines show the timing of initiation of the cytokines in the experiments and the estimated times the cytokine effects ended, respectively.

**Figure S3.**
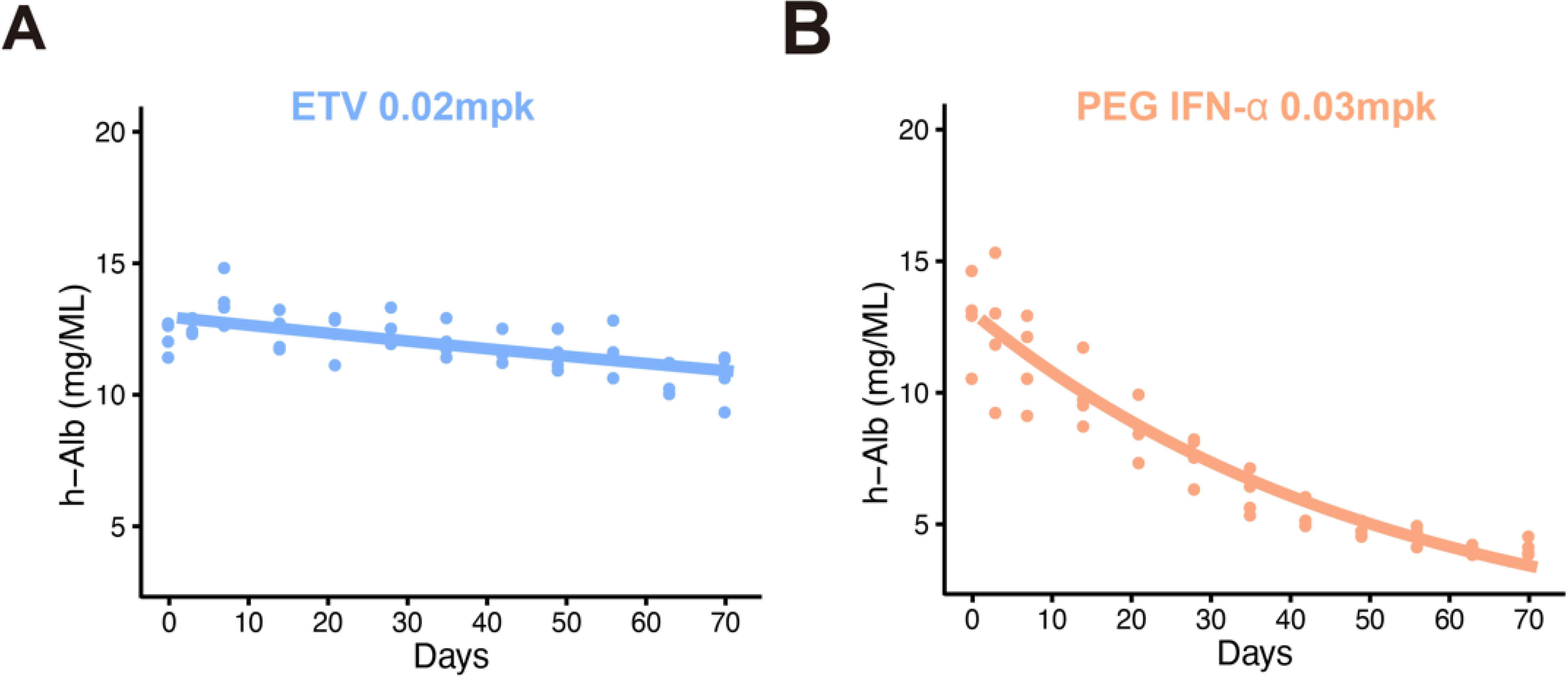
Experiments using HBV-infected humanized mice: Decay characteristics for h-Alb in peripheral blood of humanized mice treated with **(A)** ETV or **(B)** PEG IFN-α.

**Figure S4.**
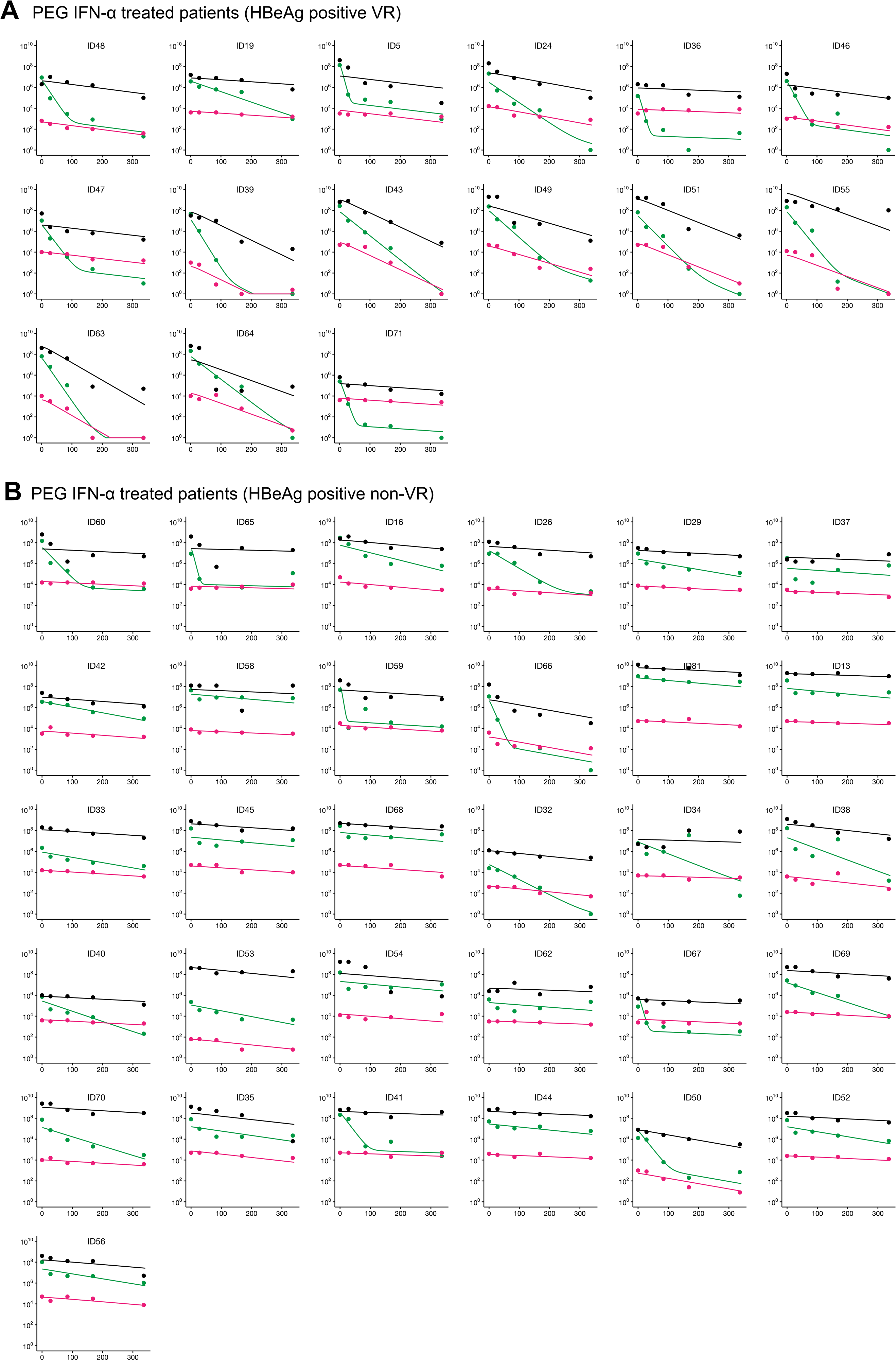
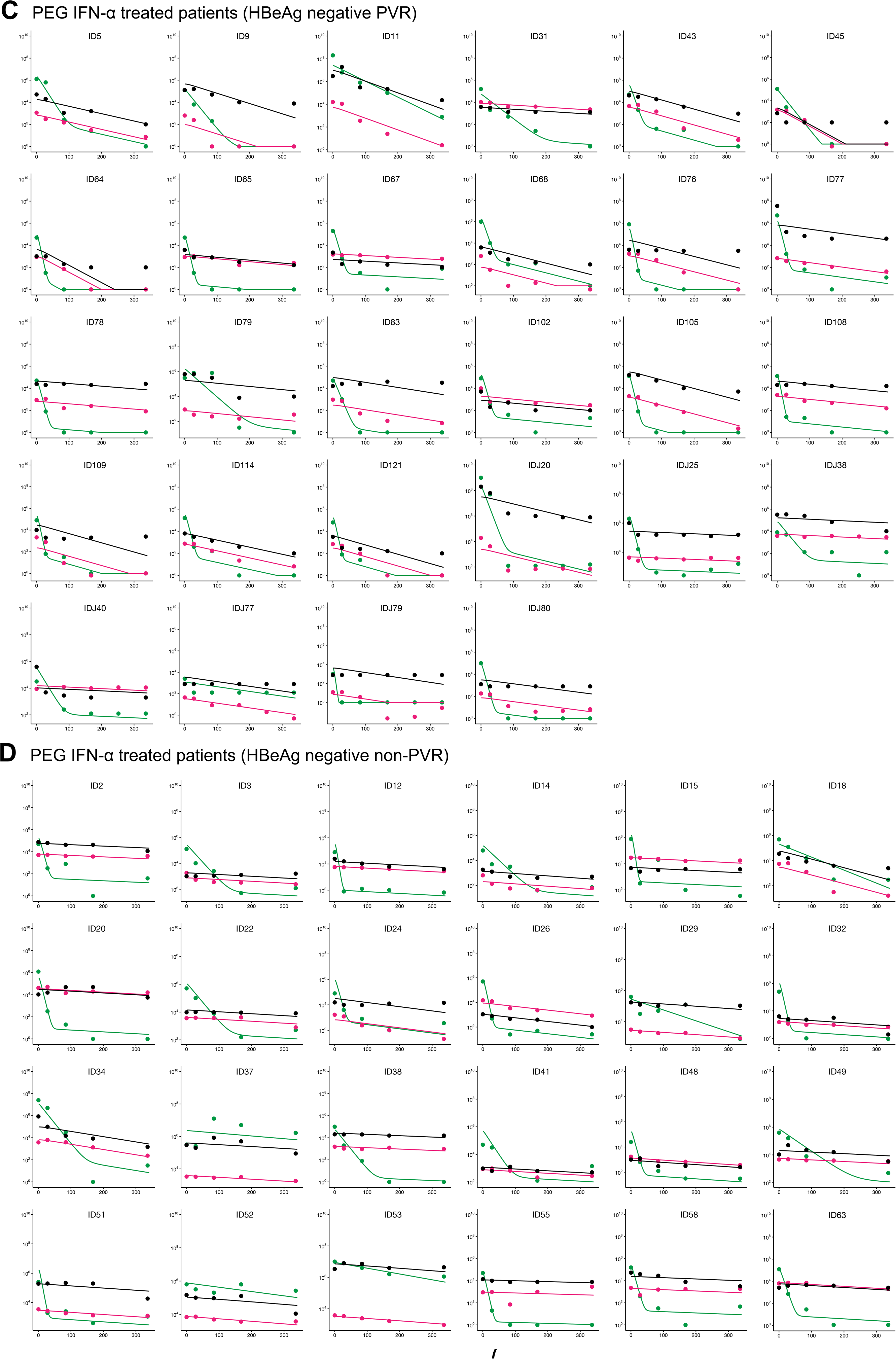
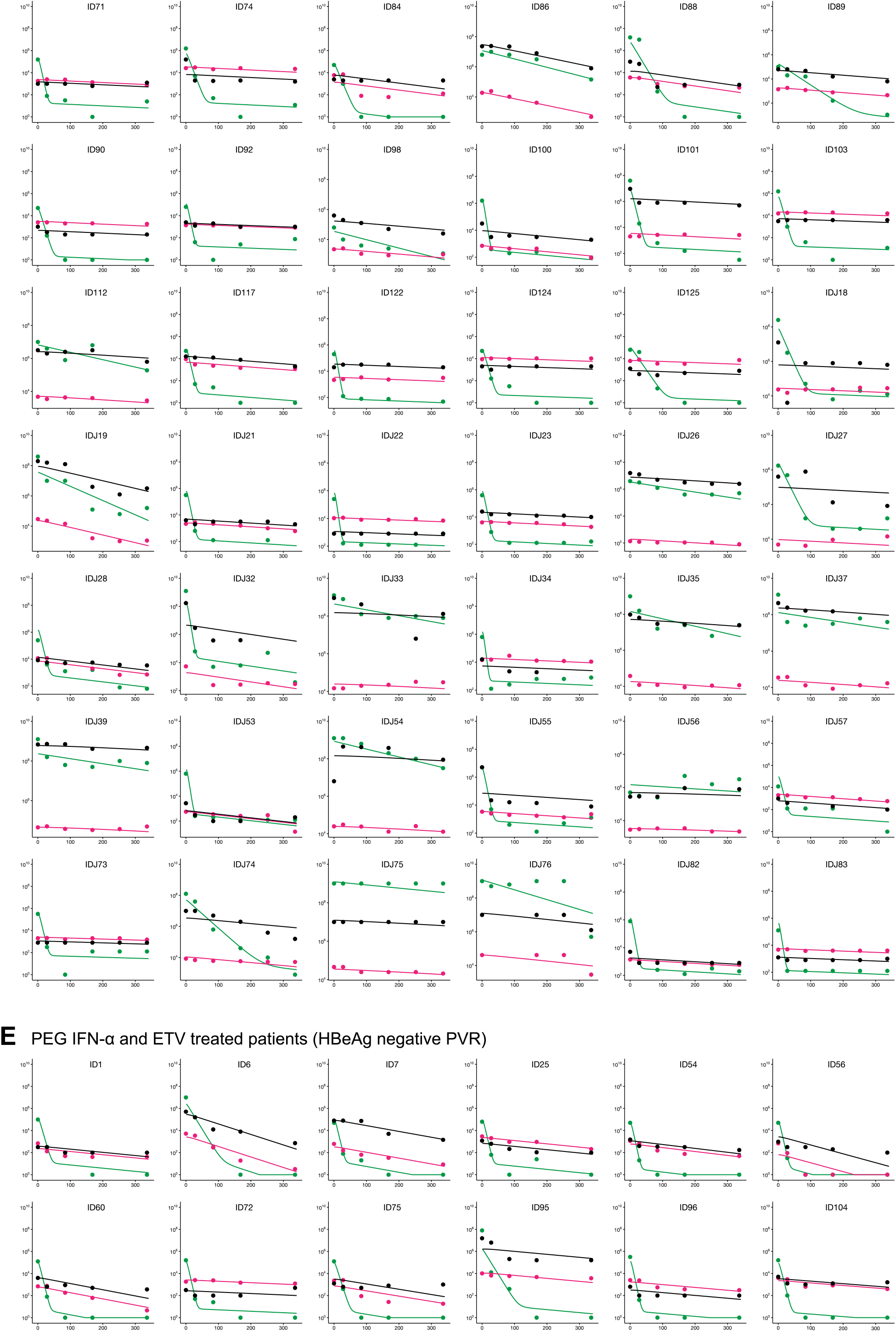
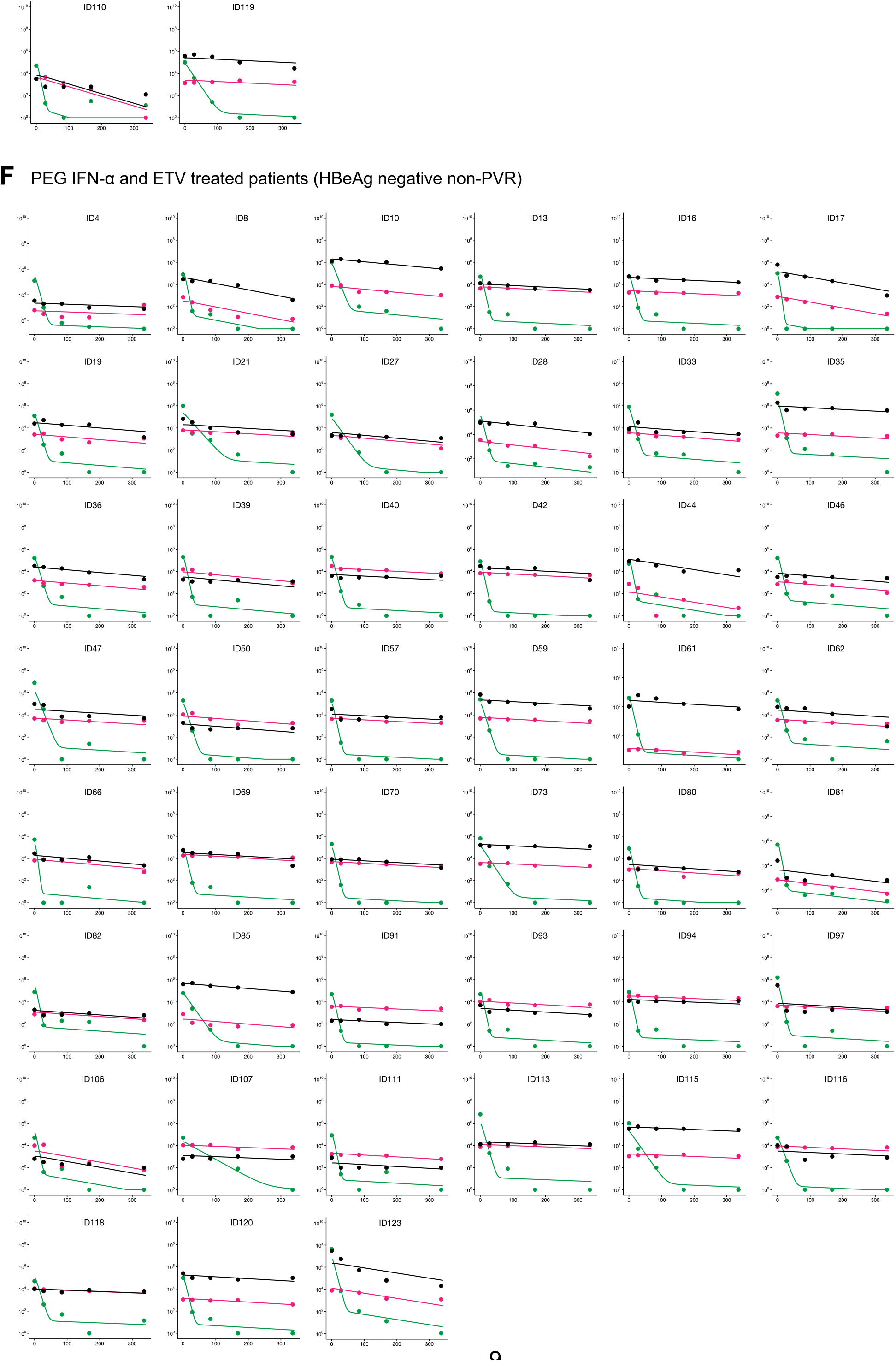
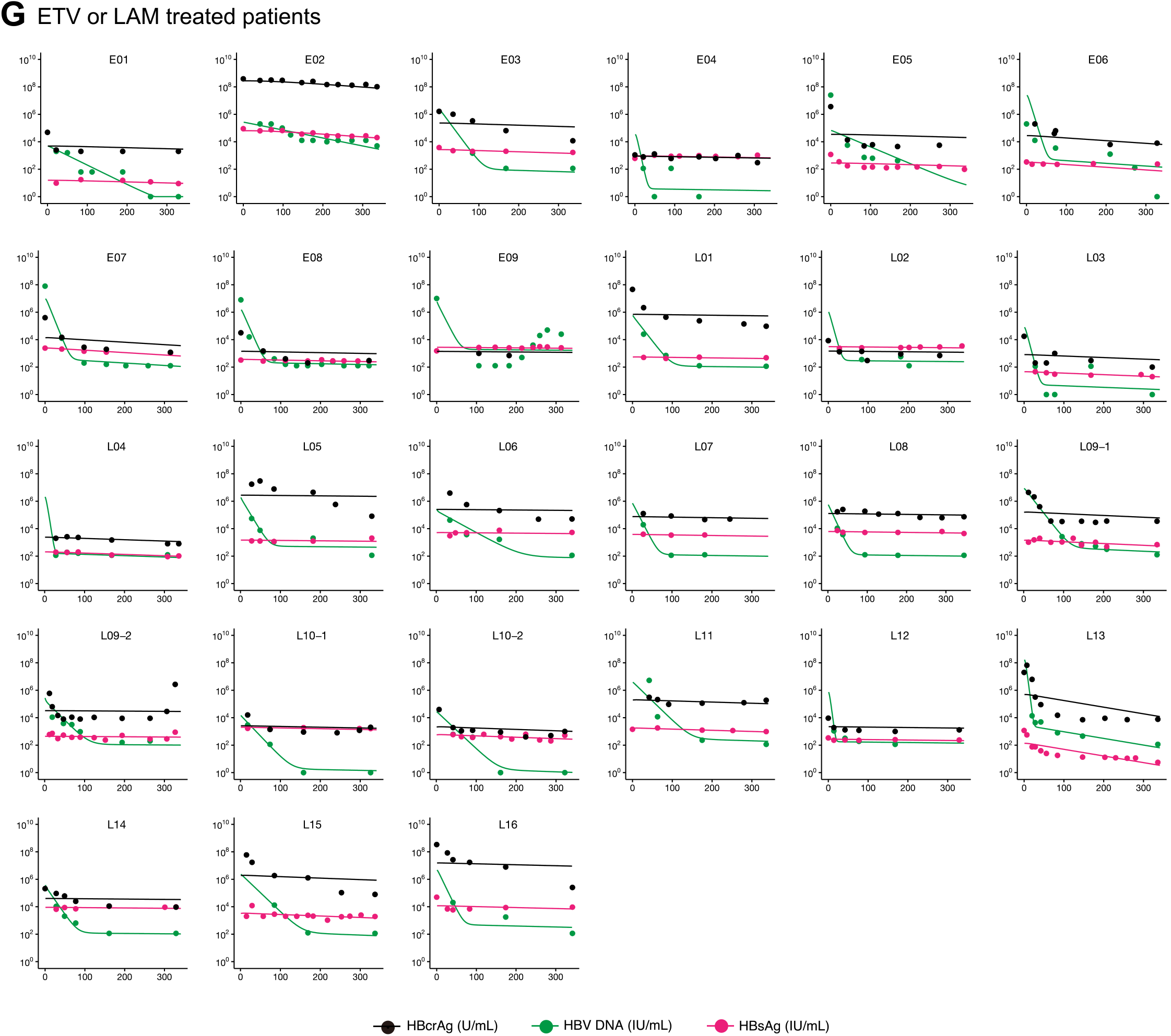
HBV-infected patients treated with PEG IFN-α or ETV/LAM: Decay characteristics are shown for extracellular HBV DNA, HBsAg and HBcrAg in peripheral blood of HBeAg-positive patients treated with PEG IFN-α **(A)** with VR or **(B)** without VR (non-VR), HBeAg-negative patients treated with PEG IFN-α **(C)** with PVR or **(D)** without PVR (non-PVR), **(E)** HBeAg-negative patients treated with PEG IFN-α and ETV with PVR **(F)** HBeAg-negative patients treated with PEG IFN-α and ETV without PVR (non-PVR) **(G)** patients treated with ETV or LAM.

**Figure S5.**
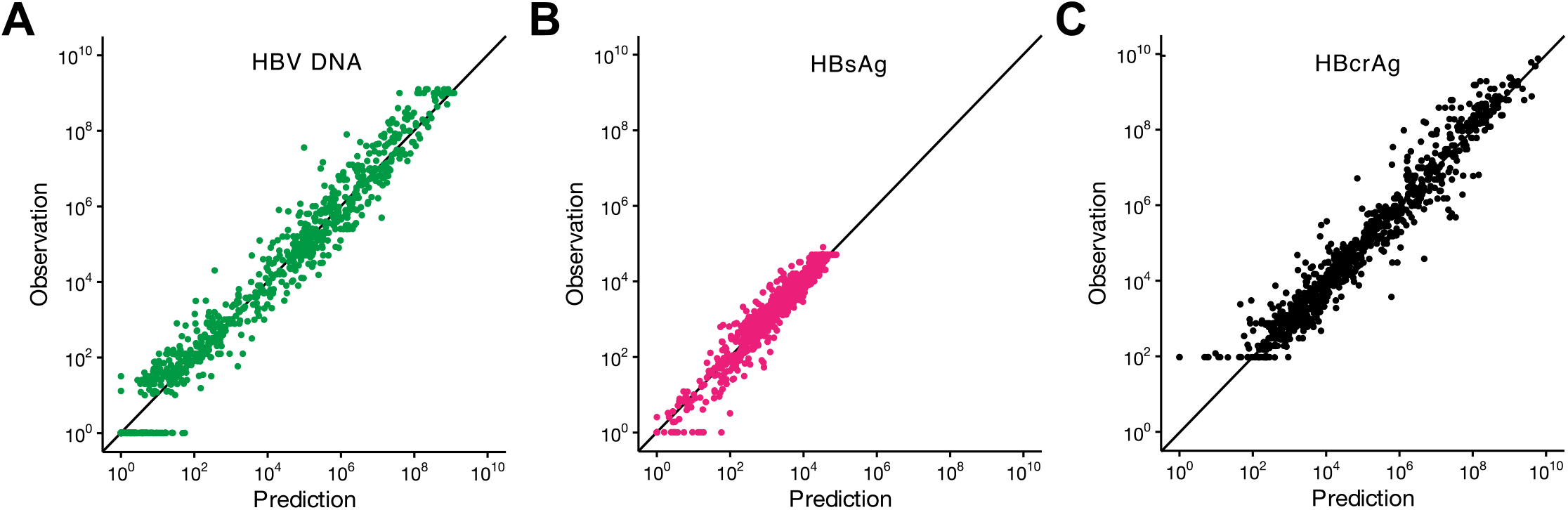
Quality of data fitting for HBV-infected patients: Correlations for observation and prediction by Eqs.(S45-46)(S48) for **(A)** HBV DNA, **(B)** HBsAg and **(C)** HBcrAg are shown.

**Figure S6.**
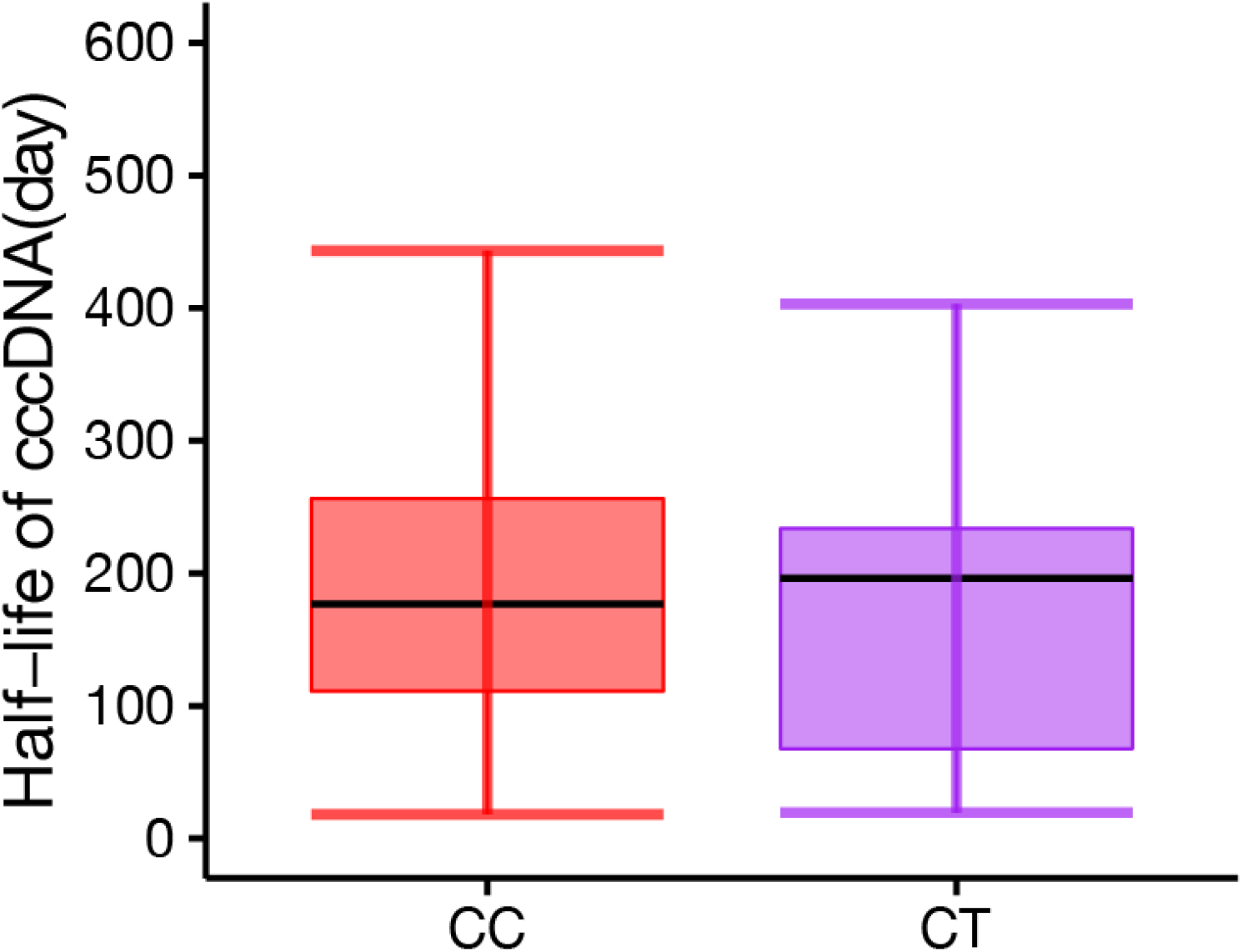
Comparison of half-life of cccDNA among different IL28B SNPs: Estimated half-life of cccDNA in hepatocyte from patients with IL28B *CC* (n=208) or *CT* (n=18) genotype treated with PEG IFN-α are shown. Black line indicates the median; box and whiskers show the interquartile range (IQR) and 1.5xIQR, respectively.

**Table S1.** Estimated parameters and initial values for HBV infection in PHH.

| Parameter or variable | Symbol | Unit | Mean | 95% CI |
| --- | --- | --- | --- | --- |
| Production rate of HBV DNA from cccDNA | $\alpha$ | day<sup>-1</sup> | $2.14 \times 10^2$ | $(0.62 - 6.32) \times 10^2$ |
| Fraction of HBV DNA recycling for cccDNA | $f$ | --- | $1.26 \times 10^{-5}$ | $2.71 \times 10^{-10} - 1.38 \times 10^{-4}$ |
| Degradation rate of cccDNA | $d$ | day<sup>-1</sup> | $1.90 \times 10^{-2}$ | $(0.34 - 4.58) \times 10^{-2}$ |
| Consumption rate of HBV DNA for virion | $\rho$ | day<sup>-1</sup> | $6.49 \times 10^{-1}$ | 0.21 - 1.77 |
| Degradation rate of extracellular HBV DNA | $d_E$ | day<sup>-1</sup> | 1.10 | 0.45 - 2.47 |
| Inhibition rate of ETV | $\varepsilon$ | --- | 0.89 | 0.75 - 0.97 |
| Initial value for cccDNA * | $C(0)$ | copies/well | $2.87 \times 10^4$ | $(1.38 - 5.23) \times 10^4$ |
| Initial value for cccDNA ** | $C(0)$ | copies/well | $2.31 \times 10^4$ | $(1.21 - 4.03) \times 10^4$ |
| Initial value for cccDNA *** | $C(0)$ | copies/well | $2.36 \times 10^4$ | $(1.25 - 403) \times 10^4$ |
| Initial value for intracellular HBV DNA* | $D(0)$ | copies/well | $1.89 \times 10^5$ | $(1.75 - 7.90) \times 10^5$ |
| Initial value for intracellular HBV DNA ** | $D(0)$ | copies/well | $3.52 \times 10^5$ | $(0.03 - 1.46) \times 10^5$ |
| Initial value for intracellular HBV DNA *** | $D(0)$ | copies/well | $1.76 \times 10^5$ | $(1.80 - 7.58) \times 10^5$ |
| Initial value for extracellular HBV DNA * | $Q(0)$ | copies/well | $1.53 \times 10^8$ | $1.61 \times 10^4 - 1.35 \times 10^9$ |
| Initial value for extracellular HBV DNA ** | $Q(0)$ | copies/well | $3.63 \times 10^8$ | $2.88 \times 10^4 - 3.82 \times 10^9$ |
| Initial value for extracellular HBV DNA *** | $Q(0)$ | copies/well | $1.80 \times 10^8$ | $1.66 \times 10^4 - 1.62 \times 10^9$ |

\* These values are estimated for condition 1.
\*\* These values are estimated for condition 2.
\*\*\* These values are estimated for condition 3.

**Table S2.** Estimated parameters and initial values for hypothetical mechanisms of action for antivirals against HBV infection in PHH.

| Parameters or variables | Symbol | Unit | Value |
| --- | --- | --- | --- |
| Production rate of HBV DNA from cccDNA | $\alpha$ | day<sup>-1</sup> | $2.14 \times 10^2$ |
| Fraction of HBV DNA recycling for cccDNA | $f$ | --- | $1.26 \times 10^{-5}$ |
| Degradation rate of cccDNA | $d$ | day<sup>-1</sup> | $1.90 \times 10^{-2}$ |
| Consumption rate of HBV DNA for virion | $\rho$ | day<sup>-1</sup> | $6.49 \times 10^{-1}$ |
| Degradation rate of extracellular HBV DNA | $d_E$ | day<sup>-1</sup> | 1.10 |
| **ETV** |  |  |  |
| Promotion rate of cccDNA degradation | $\varepsilon_d$ | --- | 11.8 |
| Inhibition rate of HBV DNA production | $\varepsilon_\alpha$ | --- | 0.90 |
| Inhibition rate of viral releasing | $\varepsilon_f$ | --- | $4.35 \times 10^{-4}$ |
| Time for cytokine non-responding on promoting cccDNA degradation | $\tau_d$ | day | 6.41 |
| Time for cytokine non-responding on inhibiting HBV DNA production | $\tau_\alpha$ | day | 24.2 |
| Time for cytokine non-responding on inhibiting viral releasing | $\tau_f$ | day | 1.01 |
| Initial value for cccDNA for promoting cccDNA degradation | $C_d(0)$ | copies/well | $1.70 \times 10^4$ |
| Initial value for cccDNA for inhibiting HBV DNA production | $C_\alpha(0)$ | copies/well | $2.02 \times 10^4$ |
| Initial value for cccDNA for inhibiting viral releasing | $C_f(0)$ | copies/well | $5.49 \times 10^3$ |
| Initial value for intracellular HBV DNA for promoting cccDNA degradation | $D_d(0)$ | copies/well | $2.32 \times 10^4$ |
| Initial value for intracellular HBV DNA for inhibiting HBV DNA production | $D_\alpha(0)$ | copies/well | $2.32 \times 10^4$ |
| Initial value for intracellular HBV DNA for inhibiting viral releasing | $D_f(0)$ | copies/well | $2.32 \times 10^4$ |
| Initial value for extracellular HBV DNA for promoting cccDNA degradation | $Q_d(0)$ | copies/well | $1.18 \times 10^8$ |
| Initial value for extracellular HBV DNA for inhibiting HBV DNA production | $Q_\alpha(0)$ | copies/well | $1.69 \times 10^8$ |
| Initial value for extracellular HBV DNA for inhibiting viral releasing | $Q_f(0)$ | copies/well | $1.56 \times 10^8$ |
| **ETV + IFN-α** |  |  |  |
| Promotion rate of cccDNA degradation | $\varepsilon_d$ | --- | 27.6 |
| Inhibition rate of HBV DNA production | $\varepsilon_\alpha$ | --- | 0.90 |
| Inhibition rate of viral releasing | $\varepsilon_f$ | --- | $3.90 \times 10^{-4}$ |
| Time for cytokine non-responding on promoting cccDNA degradation | $\tau_d$ | day | 3.24 |
| Time for cytokine non-responding on inhibiting HBV DNA production | $\tau_\alpha$ | day | 23.1 |
| Time for cytokine non-responding on inhibiting viral releasing | $\tau_f$ | day | 1.09 |
| Initial value for cccDNA for promoting cccDNA degradation | $C_d(0)$ | copies/well | $2.01 \times 10^4$ |
| Initial value for cccDNA for inhibiting HBV DNA production | $C_\alpha(0)$ | copies/well | $1.71 \times 10^4$ |
| Initial value for cccDNA for inhibiting viral releasing | $C_f(0)$ | copies/well | $4.65 \times 10^3$ |
| Initial value for intracellular HBV DNA for promoting cccDNA degradation | $D_d(0)$ | copies/well | $3.96 \times 10^4$ |
| Initial value for intracellular HBV DNA for inhibiting HBV DNA production | $D_\alpha(0)$ | copies/well | $3.96 \times 10^4$ |
| Initial value for intracellular HBV DNA for inhibiting viral releasing | $D_f(0)$ | copies/well | $3.95 \times 10^4$ |
| Initial value for extracellular HBV DNA for promoting cccDNA degradation | $Q_d(0)$ | copies/well | $1.39 \times 10^8$ |
| Initial value for extracellular HBV DNA for inhibiting HBV DNA production | $Q_\alpha(0)$ | copies/well | $2.06 \times 10^8$ |
| Initial value for extracellular HBV DNA for inhibiting viral releasing | $Q_f(0)$ | copies/well | $1.83 \times 10^8$ |
| **IFN-α** |  |  |  |
| Promotion rate of cccDNA degradation | $\varepsilon_d$ | --- | 24.2 |
| Inhibition rate of HBV DNA production | $\varepsilon_\alpha$ | --- | 0.89 |
| Inhibition rate of viral releasing | $\varepsilon_f$ | --- | $2.55 \times 10^{-4}$ |
| Time for cytokine non-responding on promoting cccDNA degradation | $\tau_d$ | day | 3.39 |
| Time for cytokine non-responding on inhibiting HBV DNA production | $\tau_\alpha$ | day | 24.5 |
| Time for cytokine non-responding on inhibiting viral releasing | $\tau_f$ | day | 1.19 |
| Initial value for cccDNA for promoting cccDNA degradation | $C_d(0)$ | copies/well | $1.83 \times 10^4$ |
| Initial value for cccDNA for inhibiting HBV DNA production | $C_\alpha(0)$ | copies/well | $1.65 \times 10^4$ |
| Initial value for cccDNA for inhibiting viral releasing | $C_f(0)$ | copies/well | $4.79 \times 10^3$ |
| Initial value for intracellular HBV DNA for promoting cccDNA degradation | $D_d(0)$ | copies/well | $3.22 \times 10^4$ |
| Initial value for intracellular HBV DNA for inhibiting HBV DNA production | $D_\alpha(0)$ | copies/well | $3.22 \times 10^4$ |
| Initial value for intracellular HBV DNA for inhibiting viral releasing | $D_f(0)$ | copies/well | $3.21 \times 10^4$ |
| Initial value for extracellular HBV DNA for promoting cccDNA degradation | $Q_d(0)$ | copies/well | $1.12 \times 10^8$ |
| Initial value for extracellular HBV DNA for inhibiting HBV DNA production | $Q_\alpha(0)$ | copies/well | $1.75 \times 10^8$ |
| Initial value for extracellular HBV DNA for inhibiting viral releasing | $Q_f(0)$ | copies/well | $1.52 \times 10^8$ |

**Table S3.** Estimated parameters for HBV infection in humanized mouse.

| Parameters or variables | Symbol | Unit | Mean | 95% CI |
| --- | --- | --- | --- | --- |
| Combined parameter<sup>†</sup> | $f\alpha$ | - | $1.9 \times 10^{-3}$ | $(0.7 - 2.8) \times 10^{-3}$ |
| Inhibition rate of HBV DNA production | $\varepsilon$ | - | $9.7 \times 10^{-1}$ | $(9.6 - 9.8) \times 10^{-1}$ |
| Decay rate of infected cells | $\delta$ | day<sup>-1</sup> | $2.4 \times 10^{-3}$ | --- |
| Decay rate of infected cells with IFN-α | $\delta_{\text{IFN}}$ | day<sup>-1</sup> | $1.9 \times 10^{-2}$ | --- |
| Degradation rate of cccDNA | $d$ | day<sup>-1</sup> | $8.8 \times 10^{-3}$ | $(7.2 - 10.5) \times 10^{-3}$ |
| Degradation rate of cccDNA with IFN-α | $d_{\text{IFN}}$ | day<sup>-1</sup> | $1.6 \times 10^{-1}$ | $(1.5 - 1.8) \times 10^{-1}$ |
| Release rate of intracellular HBV DNA | $\rho$ | day<sup>-1</sup> | $3.9 \times 10^{-1}$ | $(3.4 - 4.2) \times 10^{-1}$ |

<sup>†</sup> Production rate of HBV DNA from cccDNA × Fraction of HBV DNA recycling for cccDNA

**Table S4.**
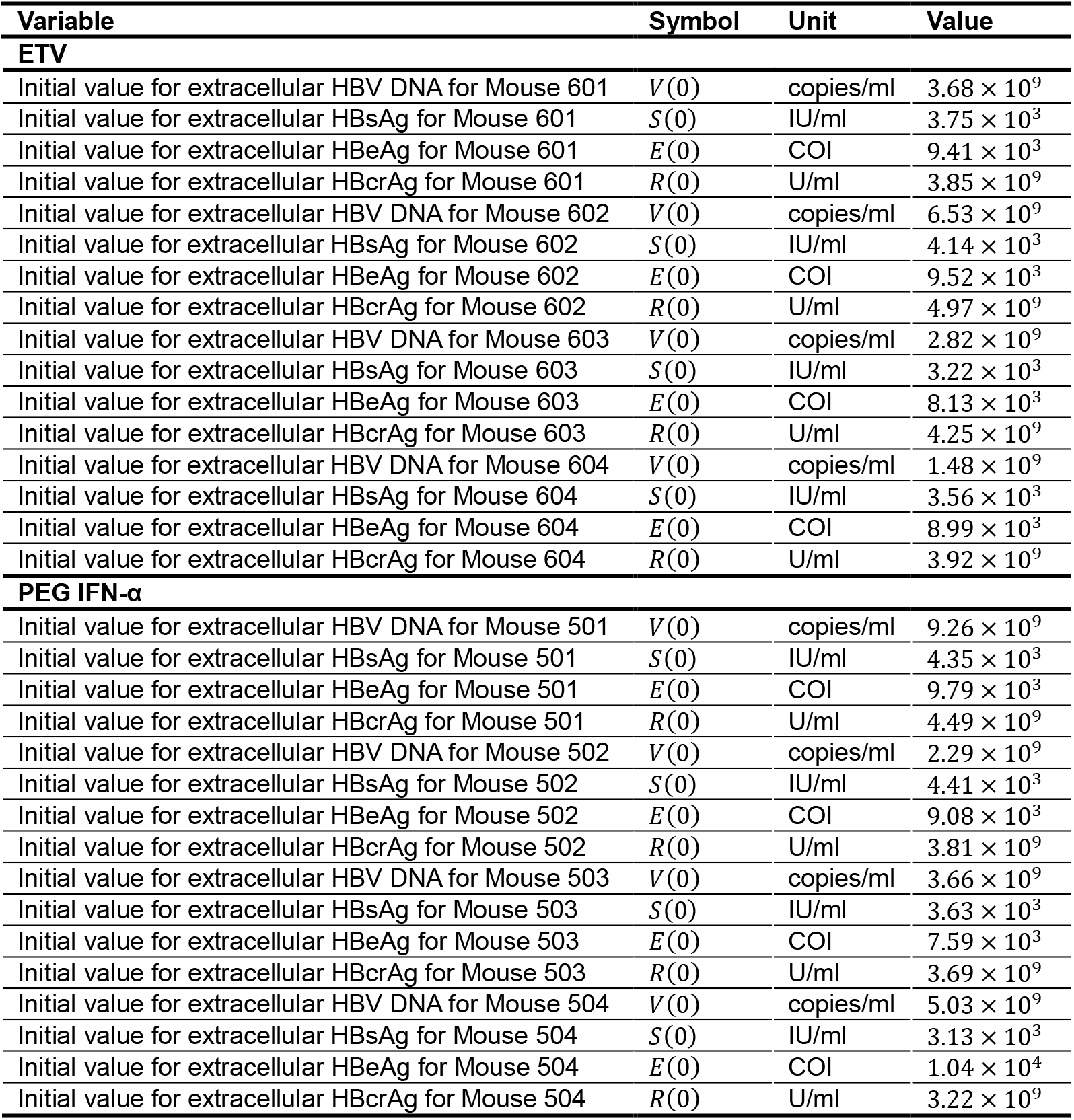
Fixed initial values for HBV infection in humanized mouse.

**Table S5.** Quantified results for cccDNA in HBV infected mouse.

| **Experimental group A** | **cccDNA<sup>†</sup>**<br>**(band volume)** | **Average**<br>**(band volume)** | **% of control** |
| --- | --- | --- | --- |
| untreated control mouse A1 | $5.11 \times 10^7$ | $4.83 \times 10^7$ | 100 |
| untreated control mouse A2 | $4.55 \times 10^7$ | — | — |
| PEG IFN-α treated mouse A1 | $1.74 \times 10^7$ | $1.60 \times 10^7$ | 33 |
| PEG IFN-α treated mouse A2 | $1.46 \times 10^7$ | — | — |
| **Experimental group B** | **cccDNA**<br>**(band volume)** | **Average**<br>**(band volume)** | **% of control** |
| untreated control mouse B1 | $1.31 \times 10^7$ | $1.13 \times 10^7$ | 100 |
| untreated control mouse B2 | $9.44 \times 10^6$ | — | — |
| PEG IFN-α treated mouse B1 | $3.14 \times 10^6$ | $2.62 \times 10^6$ | 23 |
| PEG IFN-α treated mouse B2 | $2.10 \times 10^6$ | — | — |

<sup>†</sup>cccDNA band volume was quantified from Southern blot data<sup>1</sup>. Briefly, mice infected with HBV at 12 weeks were treated with or without PEG IFN-α for 6 weeks, and then they were sacrificed. cccDNA levels were determined by Southern blot in Epicentre-based DNA extracts without proteinase K after PSD digestion. Experimental group A and B were performed as independent experiments.

**Table S6.** Estimated population parameters and initial values for HBV-infected patients treated with PEG IFN-α or ETV/LAM.

| Parameter or variable | Symbol | Unit | Value (S.E.) | I.V.* (S.E.) |
| --- | --- | --- | --- | --- |
| Combined parameter<sup>†</sup> | $f\alpha$ | - | $1.1 (0.96) \times 10^{-4}$ | - |
| Inhibition rate of HBV DNA production | $\varepsilon$ | - | 0.99 (0.00083) | - |
| Decay rate of infected cell | $\delta$ | day<sup>-1</sup> | $1.34 (0.46) \times 10^{-4}$ | 1.16 (0.50) |
| Decay rate of infected cell with PEG IFN-α | $\delta_{IFN}$ | day<sup>-1</sup> | $5.07 (0.91) \times 10^{-4}$ | 0.45 (0.24) |
| Consumption rate of HBV DNA for virion | $\rho$ | day<sup>-1</sup> | $1.18 (0.17) \times 10^{-1}$ | 1.88 (0.15) |
| Degradation rate of cccDNA | $d$ | day<sup>-1</sup> | $6.94 (2.44) \times 10^{-4}$ | 1.70 (0.25) |
| Degradation rate of cccDNA with PEG IFN-α | $d_{IFN}$ | day<sup>-1</sup> | $3.23 (0.4) \times 10^{-3}$ | 1.00 (0.09) |
| Initial value for extracellular HBV DNA for PEG IFN-α-treated patients | $V(0)$ | IU/ml | $7.97 (1.77) \times 10^5$ | 2.84 (0.16) |
| Initial value for extracellular HBsAg for PEG IFN-α-treated patients | $S(0)$ | IU/ml | $3.23 (0.39) \times 10^3$ | 1.63 (0.09) |
| Initial value for extracellular HBcrAg for PEG IFN-α-treated patients | $R(0)$ | IU/ml | $1.58 (0.49) \times 10^5$ | 4.37 (0.22) |
| Initial value for extracellular HBV DNA for ETV or LAM-treated patients | $V(0)$ | IU/ml | $4.40 (2.64) \times 10^5$ | 2.92 (0.42) |
| Initial value for extracellular HBsAg for ETV or LAM-treated patients | $S(0)$ | IU/ml | $1.15 (0.37) \times 10^2$ | 1.78 (0.30) |
| Initial value for extracellular HBcrAg for ETV or LAM-treated patients | $R(0)$ | IU/ml | $2.94 (2.00) \times 10^4$ | 3.55 (0.52) |

\* Interpatient variability.
<sup>†</sup> Production rate of HBV DNA from cccDNA × Fraction of HBV DNA recycling for cccDNA.

**Table S7.**
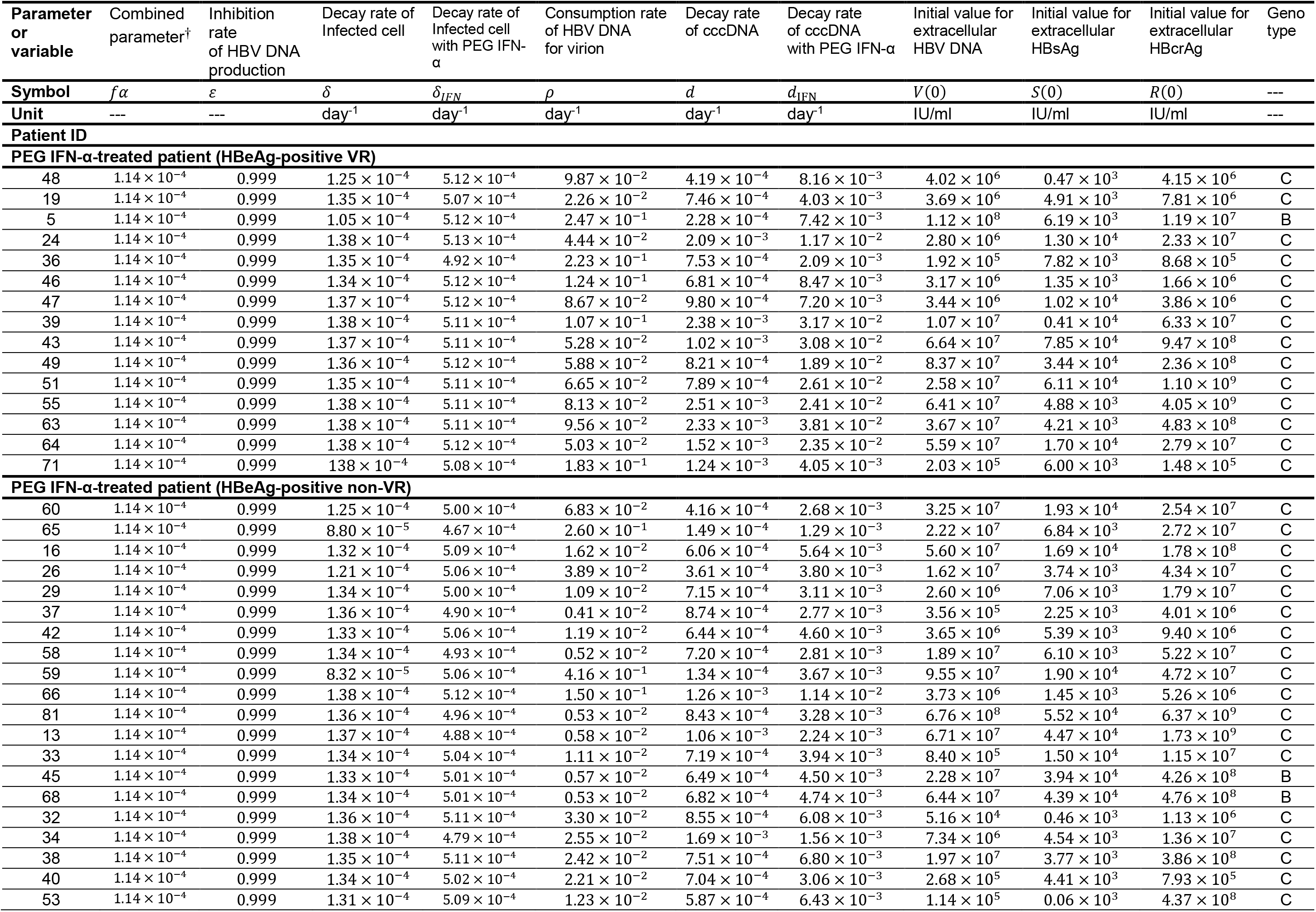
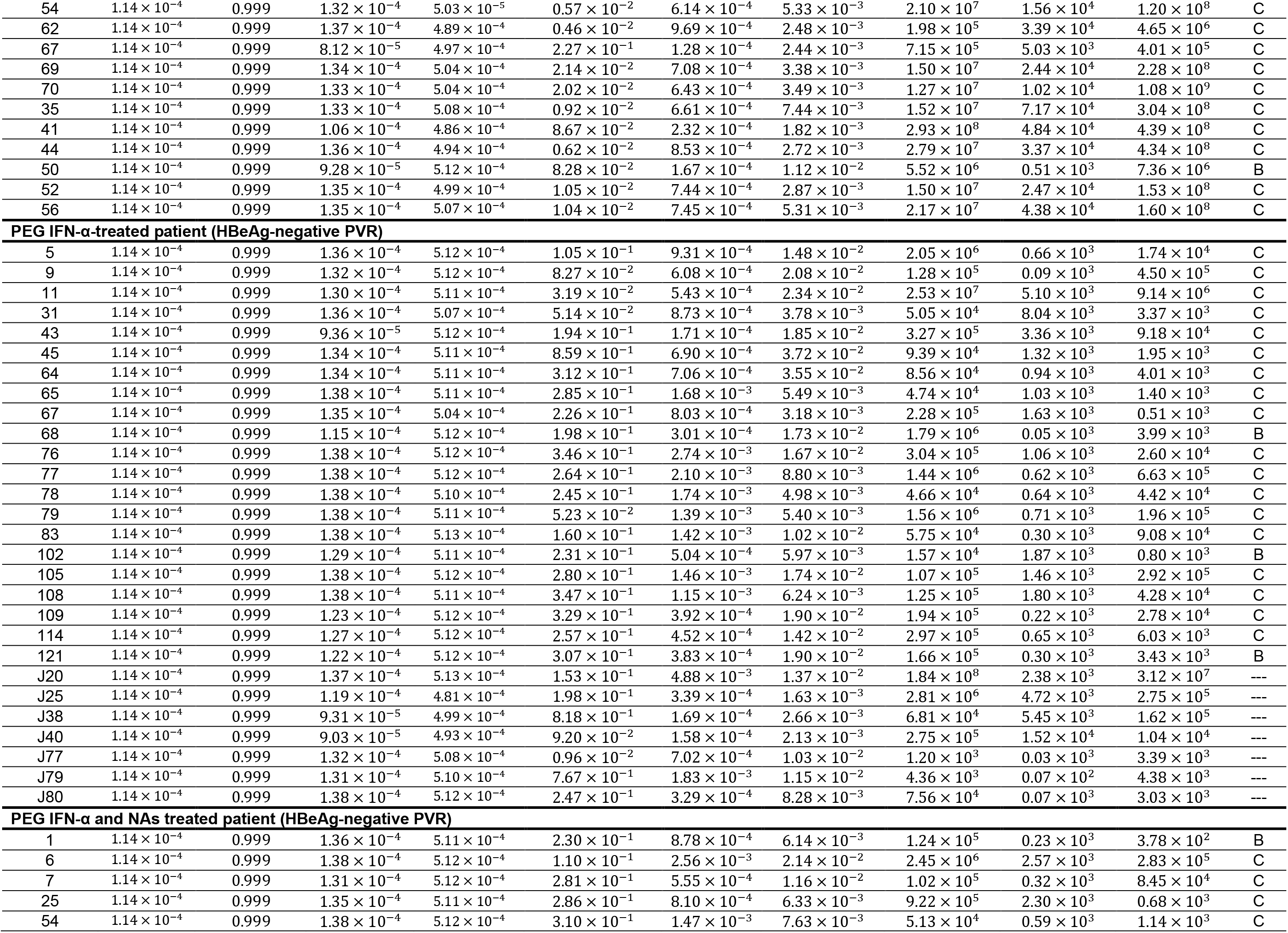
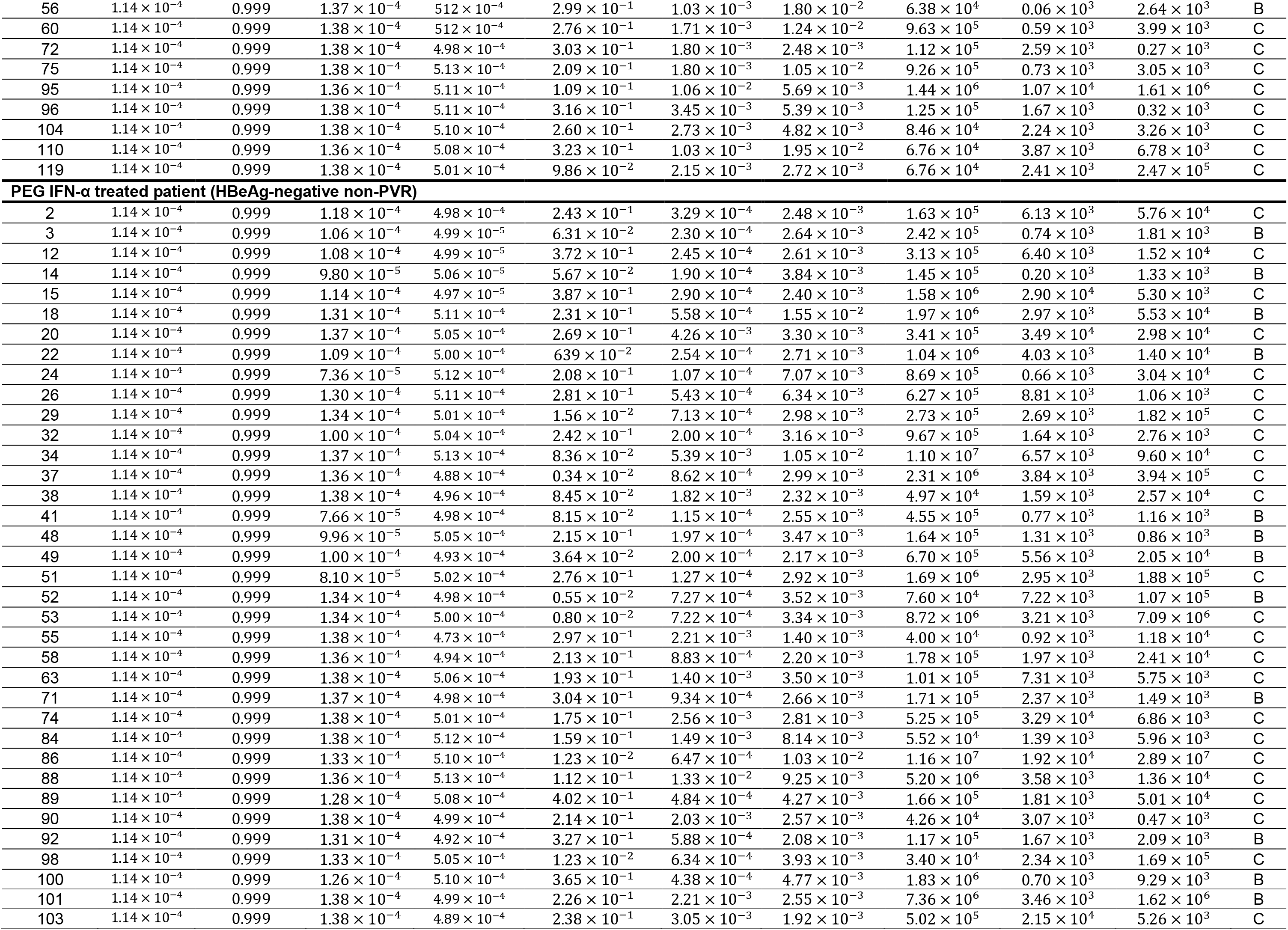
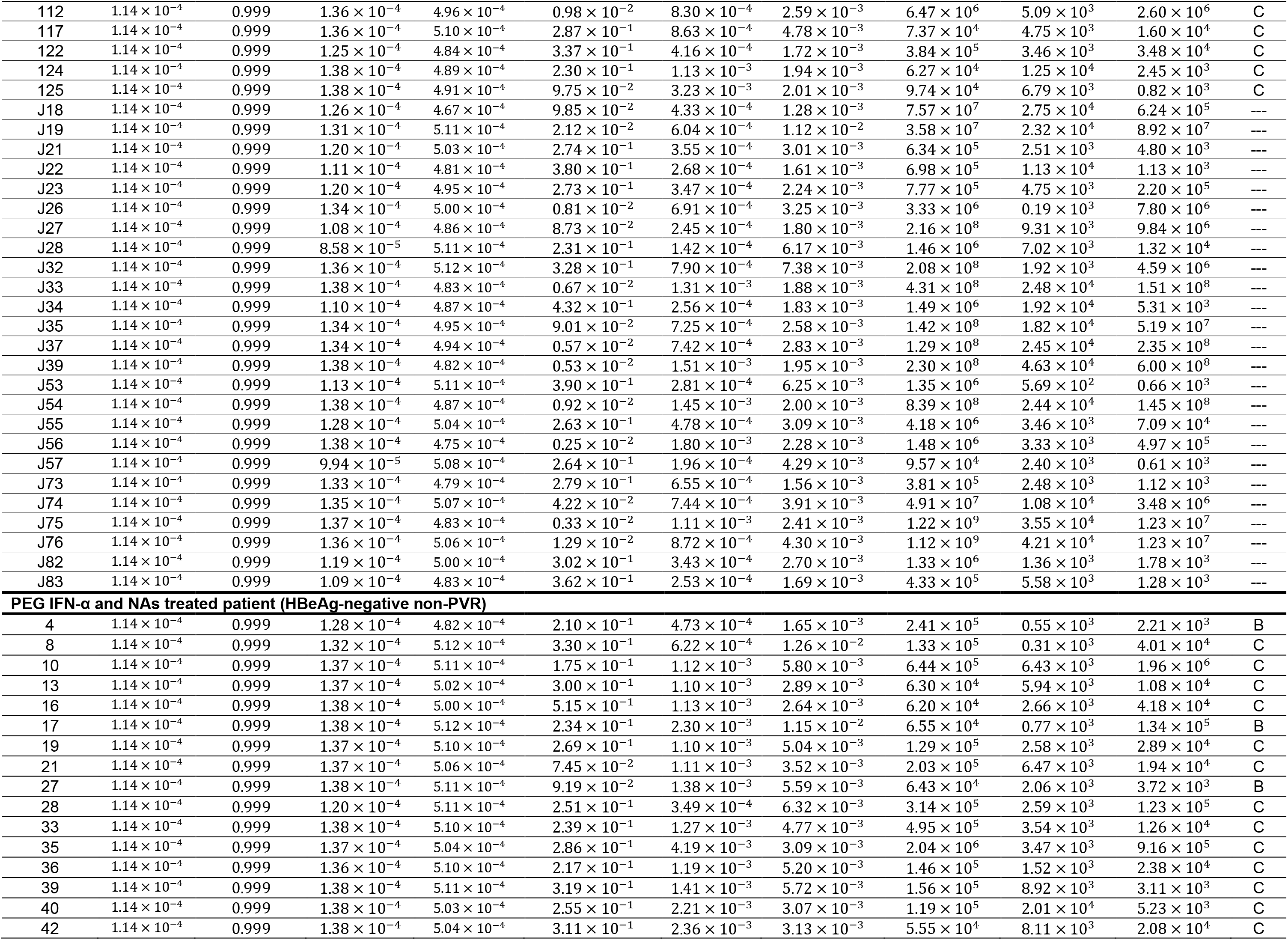
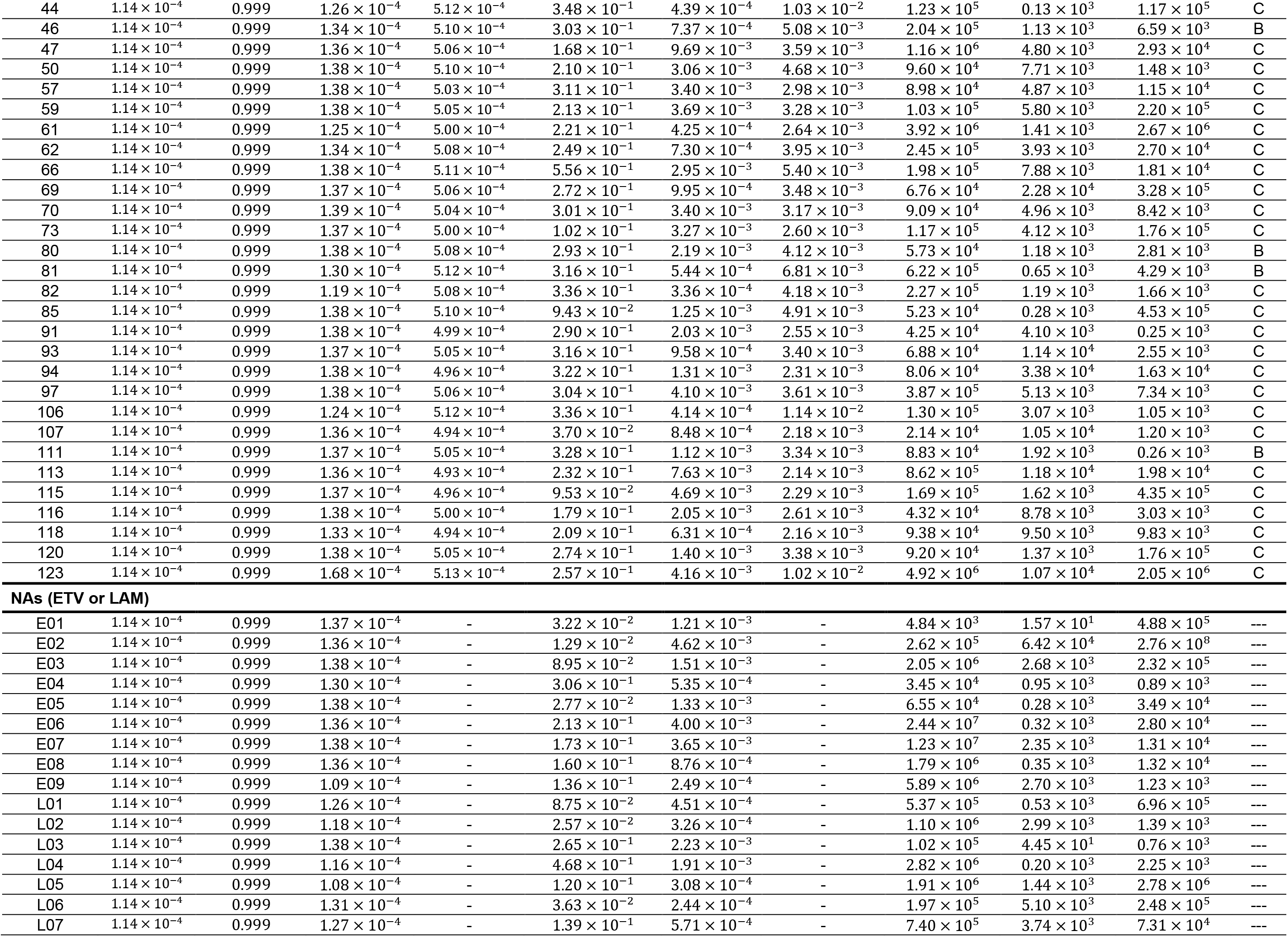
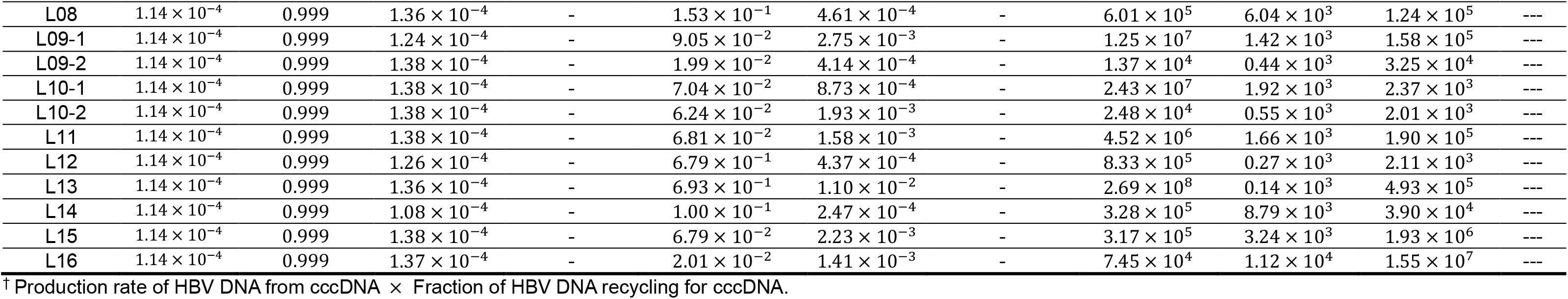
Estimated individual parameters and initial values for HBV-infected patients treated with PEG IFN-α or ETV/LAM.

| Parameter or variable | Combined parameter<sup>†</sup> | Inhibition rate of HBV DNA production | Decay rate of Infected cell | Decay rate of Infected cell with PEG IFN-α | Consumption rate of HBV DNA for virion | Decay rate of cccDNA | Decay rate of cccDNA with PEG IFN-α | Initial value for extracellular HBV DNA | Initial value for extracellular HBsAg | Initial value for extracellular HBcrAg | Geno type |
| --- | --- | --- | --- | --- | --- | --- | --- | --- | --- | --- | --- |
| Symbol | $f\alpha$ | $\varepsilon$ | $\delta$ | $\delta_{IFN}$ | $\rho$ | $d$ | $d_{IFN}$ | $V(0)$ | $S(0)$ | $R(0)$ | --- |
| Unit | --- | --- | day<sup>-1</sup> | day<sup>-1</sup> | day<sup>-1</sup> | day<sup>-1</sup> | day<sup>-1</sup> | IU/ml | IU/ml | IU/ml | --- |
| **Patient ID** |  |  |  |  |  |  |  |  |  |  |  |
| **PEG IFN-α-treated patient (HBeAg-positive VR)** |  |  |  |  |  |  |  |  |  |  |  |
| 48 | $1.14 \times 10^{-4}$ | 0.999 | $1.25 \times 10^{-4}$ | $5.12 \times 10^{-4}$ | $9.87 \times 10^{-2}$ | $4.19 \times 10^{-4}$ | $8.16 \times 10^{-3}$ | $4.02 \times 10^6$ | $0.47 \times 10^3$ | $4.15 \times 10^6$ | C |
| 19 | $1.14 \times 10^{-4}$ | 0.999 | $1.35 \times 10^{-4}$ | $5.07 \times 10^{-4}$ | $2.26 \times 10^{-2}$ | $7.46 \times 10^{-4}$ | $4.03 \times 10^{-3}$ | $3.69 \times 10^6$ | $4.91 \times 10^3$ | $7.81 \times 10^6$ | C |
| 5 | $1.14 \times 10^{-4}$ | 0.999 | $1.05 \times 10^{-4}$ | $5.12 \times 10^{-4}$ | $2.47 \times 10^{-1}$ | $2.28 \times 10^{-4}$ | $7.42 \times 10^{-3}$ | $1.12 \times 10^8$ | $6.19 \times 10^3$ | $1.19 \times 10^7$ | B |
| 24 | $1.14 \times 10^{-4}$ | 0.999 | $1.38 \times 10^{-4}$ | $5.13 \times 10^{-4}$ | $4.44 \times 10^{-2}$ | $2.09 \times 10^{-3}$ | $1.17 \times 10^{-2}$ | $2.80 \times 10^6$ | $1.30 \times 10^4$ | $2.33 \times 10^7$ | C |
| 36 | $1.14 \times 10^{-4}$ | 0.999 | $1.35 \times 10^{-4}$ | $4.92 \times 10^{-4}$ | $2.23 \times 10^{-1}$ | $7.53 \times 10^{-4}$ | $2.09 \times 10^{-3}$ | $1.92 \times 10^5$ | $7.82 \times 10^3$ | $8.68 \times 10^5$ | C |
| 46 | $1.14 \times 10^{-4}$ | 0.999 | $1.34 \times 10^{-4}$ | $5.12 \times 10^{-4}$ | $1.24 \times 10^{-1}$ | $6.81 \times 10^{-4}$ | $8.47 \times 10^{-3}$ | $3.17 \times 10^6$ | $1.35 \times 10^3$ | $1.66 \times 10^6$ | C |
| 47 | $1.14 \times 10^{-4}$ | 0.999 | $1.37 \times 10^{-4}$ | $5.12 \times 10^{-4}$ | $8.67 \times 10^{-2}$ | $9.80 \times 10^{-4}$ | $7.20 \times 10^{-3}$ | $3.44 \times 10^6$ | $1.02 \times 10^4$ | $3.86 \times 10^6$ | C |
| 39 | $1.14 \times 10^{-4}$ | 0.999 | $1.38 \times 10^{-4}$ | $5.11 \times 10^{-4}$ | $1.07 \times 10^{-1}$ | $2.38 \times 10^{-3}$ | $3.17 \times 10^{-2}$ | $1.07 \times 10^7$ | $0.41 \times 10^4$ | $6.33 \times 10^7$ | C |
| 43 | $1.14 \times 10^{-4}$ | 0.999 | $1.37 \times 10^{-4}$ | $5.11 \times 10^{-4}$ | $5.28 \times 10^{-2}$ | $1.02 \times 10^{-3}$ | $3.08 \times 10^{-2}$ | $6.64 \times 10^7$ | $7.85 \times 10^4$ | $9.47 \times 10^8$ | C |
| 49 | $1.14 \times 10^{-4}$ | 0.999 | $1.36 \times 10^{-4}$ | $5.12 \times 10^{-4}$ | $5.88 \times 10^{-2}$ | $8.21 \times 10^{-4}$ | $1.89 \times 10^{-2}$ | $8.37 \times 10^7$ | $3.44 \times 10^4$ | $2.36 \times 10^8$ | C |
| 51 | $1.14 \times 10^{-4}$ | 0.999 | $1.35 \times 10^{-4}$ | $5.11 \times 10^{-4}$ | $6.65 \times 10^{-2}$ | $7.89 \times 10^{-4}$ | $2.61 \times 10^{-2}$ | $2.58 \times 10^7$ | $6.11 \times 10^4$ | $1.10 \times 10^9$ | C |
| 55 | $1.14 \times 10^{-4}$ | 0.999 | $1.38 \times 10^{-4}$ | $5.11 \times 10^{-4}$ | $8.13 \times 10^{-2}$ | $2.51 \times 10^{-3}$ | $2.41 \times 10^{-2}$ | $6.41 \times 10^7$ | $4.88 \times 10^3$ | $4.05 \times 10^9$ | C |
| 63 | $1.14 \times 10^{-4}$ | 0.999 | $1.38 \times 10^{-4}$ | $5.11 \times 10^{-4}$ | $9.56 \times 10^{-2}$ | $2.33 \times 10^{-3}$ | $3.81 \times 10^{-2}$ | $3.67 \times 10^7$ | $4.21 \times 10^3$ | $4.83 \times 10^8$ | C |
| 64 | $1.14 \times 10^{-4}$ | 0.999 | $1.38 \times 10^{-4}$ | $5.12 \times 10^{-4}$ | $5.03 \times 10^{-2}$ | $1.52 \times 10^{-3}$ | $2.35 \times 10^{-2}$ | $5.59 \times 10^7$ | $1.70 \times 10^4$ | $2.79 \times 10^7$ | C |
| 71 | $1.14 \times 10^{-4}$ | 0.999 | $1.38 \times 10^{-4}$ | $5.08 \times 10^{-4}$ | $1.83 \times 10^{-1}$ | $1.24 \times 10^{-3}$ | $4.05 \times 10^{-3}$ | $2.03 \times 10^5$ | $6.00 \times 10^3$ | $1.48 \times 10^5$ | C |
| **PEG IFN-α-treated patient (HBeAg-positive non-VR)** |  |  |  |  |  |  |  |  |  |  |  |
| 60 | $1.14 \times 10^{-4}$ | 0.999 | $1.25 \times 10^{-4}$ | $5.00 \times 10^{-4}$ | $6.83 \times 10^{-2}$ | $4.16 \times 10^{-4}$ | $2.68 \times 10^{-3}$ | $3.25 \times 10^7$ | $1.93 \times 10^4$ | $2.54 \times 10^7$ | C |
| 65 | $1.14 \times 10^{-4}$ | 0.999 | $8.80 \times 10^{-5}$ | $4.67 \times 10^{-4}$ | $2.60 \times 10^{-1}$ | $1.49 \times 10^{-4}$ | $1.29 \times 10^{-3}$ | $2.22 \times 10^7$ | $6.84 \times 10^3$ | $2.72 \times 10^7$ | C |
| 16 | $1.14 \times 10^{-4}$ | 0.999 | $1.32 \times 10^{-4}$ | $5.09 \times 10^{-4}$ | $1.62 \times 10^{-2}$ | $6.06 \times 10^{-4}$ | $5.64 \times 10^{-3}$ | $5.60 \times 10^7$ | $1.69 \times 10^4$ | $1.78 \times 10^8$ | C |
| 26 | $1.14 \times 10^{-4}$ | 0.999 | $1.21 \times 10^{-4}$ | $5.06 \times 10^{-4}$ | $3.89 \times 10^{-2}$ | $3.61 \times 10^{-4}$ | $3.80 \times 10^{-3}$ | $1.62 \times 10^7$ | $3.74 \times 10^3$ | $4.34 \times 10^7$ | C |
| 29 | $1.14 \times 10^{-4}$ | 0.999 | $1.34 \times 10^{-4}$ | $5.00 \times 10^{-4}$ | $1.09 \times 10^{-2}$ | $7.15 \times 10^{-4}$ | $3.11 \times 10^{-3}$ | $2.60 \times 10^6$ | $7.06 \times 10^3$ | $1.79 \times 10^7$ | C |
| 37 | $1.14 \times 10^{-4}$ | 0.999 | $1.36 \times 10^{-4}$ | $4.90 \times 10^{-4}$ | $0.41 \times 10^{-2}$ | $8.74 \times 10^{-4}$ | $2.77 \times 10^{-3}$ | $3.56 \times 10^5$ | $2.25 \times 10^3$ | $4.01 \times 10^6$ | C |
| 42 | $1.14 \times 10^{-4}$ | 0.999 | $1.33 \times 10^{-4}$ | $5.06 \times 10^{-4}$ | $1.19 \times 10^{-2}$ | $6.44 \times 10^{-4}$ | $4.60 \times 10^{-3}$ | $3.65 \times 10^6$ | $5.39 \times 10^3$ | $9.40 \times 10^6$ | C |
| 58 | $1.14 \times 10^{-4}$ | 0.999 | $1.34 \times 10^{-4}$ | $4.93 \times 10^{-4}$ | $0.52 \times 10^{-2}$ | $7.20 \times 10^{-4}$ | $2.81 \times 10^{-3}$ | $1.89 \times 10^7$ | $6.10 \times 10^3$ | $5.22 \times 10^7$ | C |
| 59 | $1.14 \times 10^{-4}$ | 0.999 | $8.32 \times 10^{-5}$ | $5.06 \times 10^{-4}$ | $4.16 \times 10^{-1}$ | $1.34 \times 10^{-4}$ | $3.67 \times 10^{-3}$ | $9.55 \times 10^7$ | $1.90 \times 10^4$ | $4.72 \times 10^7$ | C |
| 66 | $1.14 \times 10^{-4}$ | 0.999 | $1.38 \times 10^{-4}$ | $5.12 \times 10^{-4}$ | $1.50 \times 10^{-1}$ | $1.26 \times 10^{-3}$ | $1.14 \times 10^{-2}$ | $3.73 \times 10^6$ | $1.45 \times 10^3$ | $5.26 \times 10^6$ | C |
| 81 | $1.14 \times 10^{-4}$ | 0.999 | $1.36 \times 10^{-4}$ | $4.96 \times 10^{-4}$ | $0.53 \times 10^{-2}$ | $8.43 \times 10^{-4}$ | $3.28 \times 10^{-3}$ | $6.76 \times 10^8$ | $5.52 \times 10^4$ | $6.37 \times 10^9$ | C |
| 13 | $1.14 \times 10^{-4}$ | 0.999 | $1.37 \times 10^{-4}$ | $4.88 \times 10^{-4}$ | $0.58 \times 10^{-2}$ | $1.06 \times 10^{-3}$ | $2.24 \times 10^{-3}$ | $6.71 \times 10^7$ | $4.47 \times 10^4$ | $1.73 \times 10^9$ | C |
| 33 | $1.14 \times 10^{-4}$ | 0.999 | $1.34 \times 10^{-4}$ | $5.04 \times 10^{-4}$ | $1.11 \times 10^{-2}$ | $7.19 \times 10^{-4}$ | $3.94 \times 10^{-3}$ | $8.40 \times 10^5$ | $1.50 \times 10^4$ | $1.15 \times 10^7$ | C |
| 45 | $1.14 \times 10^{-4}$ | 0.999 | $1.33 \times 10^{-4}$ | $5.01 \times 10^{-4}$ | $0.57 \times 10^{-2}$ | $6.49 \times 10^{-4}$ | $4.50 \times 10^{-3}$ | $2.28 \times 10^7$ | $3.94 \times 10^4$ | $4.26 \times 10^8$ | B |
| 68 | $1.14 \times 10^{-4}$ | 0.999 | $1.34 \times 10^{-4}$ | $5.01 \times 10^{-4}$ | $0.53 \times 10^{-2}$ | $6.82 \times 10^{-4}$ | $4.74 \times 10^{-3}$ | $6.44 \times 10^7$ | $4.39 \times 10^4$ | $4.76 \times 10^8$ | B |
| 32 | $1.14 \times 10^{-4}$ | 0.999 | $1.36 \times 10^{-4}$ | $5.11 \times 10^{-4}$ | $3.30 \times 10^{-2}$ | $8.55 \times 10^{-4}$ | $6.08 \times 10^{-3}$ | $5.16 \times 10^4$ | $0.46 \times 10^3$ | $1.13 \times 10^6$ | C |
| 34 | $1.14 \times 10^{-4}$ | 0.999 | $1.38 \times 10^{-4}$ | $4.79 \times 10^{-4}$ | $2.55 \times 10^{-2}$ | $1.69 \times 10^{-3}$ | $1.56 \times 10^{-3}$ | $7.34 \times 10^6$ | $4.54 \times 10^3$ | $1.36 \times 10^7$ | C |
| 38 | $1.14 \times 10^{-4}$ | 0.999 | $1.35 \times 10^{-4}$ | $5.11 \times 10^{-4}$ | $2.42 \times 10^{-2}$ | $7.51 \times 10^{-4}$ | $6.80 \times 10^{-3}$ | $1.97 \times 10^7$ | $3.77 \times 10^3$ | $3.86 \times 10^8$ | C |
| 40 | $1.14 \times 10^{-4}$ | 0.999 | $1.34 \times 10^{-4}$ | $5.02 \times 10^{-4}$ | $2.21 \times 10^{-2}$ | $7.04 \times 10^{-4}$ | $3.06 \times 10^{-3}$ | $2.68 \times 10^5$ | $4.41 \times 10^3$ | $7.93 \times 10^5$ | C |
| 53 | $1.14 \times 10^{-4}$ | 0.999 | $1.31 \times 10^{-4}$ | $5.09 \times 10^{-4}$ | $1.23 \times 10^{-2}$ | $5.87 \times 10^{-4}$ | $6.43 \times 10^{-3}$ | $1.14 \times 10^5$ | $0.06 \times 10^3$ | $4.37 \times 10^8$ | C |
| 54 | $1.14 \times 10^{-4}$ | 0.999 | $1.32 \times 10^{-4}$ | $5.03 \times 10^{-5}$ | $0.57 \times 10^{-2}$ | $6.14 \times 10^{-4}$ | $5.33 \times 10^{-3}$ | $2.10 \times 10^7$ | $1.56 \times 10^4$ | $1.20 \times 10^8$ | C |
| 62 | $1.14 \times 10^{-4}$ | 0.999 | $1.37 \times 10^{-4}$ | $4.89 \times 10^{-4}$ | $0.46 \times 10^{-2}$ | $9.69 \times 10^{-4}$ | $2.48 \times 10^{-3}$ | $1.98 \times 10^5$ | $3.39 \times 10^4$ | $4.65 \times 10^6$ | C |
| 67 | $1.14 \times 10^{-4}$ | 0.999 | $8.12 \times 10^{-5}$ | $4.97 \times 10^{-4}$ | $2.27 \times 10^{-1}$ | $1.28 \times 10^{-4}$ | $2.44 \times 10^{-3}$ | $7.15 \times 10^5$ | $5.03 \times 10^3$ | $4.01 \times 10^5$ | C |
| 69 | $1.14 \times 10^{-4}$ | 0.999 | $1.34 \times 10^{-4}$ | $5.04 \times 10^{-4}$ | $2.14 \times 10^{-2}$ | $7.08 \times 10^{-4}$ | $3.38 \times 10^{-3}$ | $1.50 \times 10^7$ | $2.44 \times 10^4$ | $2.28 \times 10^8$ | C |
| 70 | $1.14 \times 10^{-4}$ | 0.999 | $1.33 \times 10^{-4}$ | $5.04 \times 10^{-4}$ | $2.02 \times 10^{-2}$ | $6.43 \times 10^{-4}$ | $3.49 \times 10^{-3}$ | $1.27 \times 10^7$ | $1.02 \times 10^4$ | $1.08 \times 10^9$ | C |
| 35 | $1.14 \times 10^{-4}$ | 0.999 | $1.33 \times 10^{-4}$ | $5.08 \times 10^{-4}$ | $0.92 \times 10^{-2}$ | $6.61 \times 10^{-4}$ | $7.44 \times 10^{-3}$ | $1.52 \times 10^7$ | $7.17 \times 10^4$ | $3.04 \times 10^8$ | C |
| 41 | $1.14 \times 10^{-4}$ | 0.999 | $1.06 \times 10^{-4}$ | $4.86 \times 10^{-4}$ | $8.67 \times 10^{-2}$ | $2.32 \times 10^{-4}$ | $1.82 \times 10^{-3}$ | $2.93 \times 10^8$ | $4.84 \times 10^4$ | $4.39 \times 10^8$ | C |
| 44 | $1.14 \times 10^{-4}$ | 0.999 | $1.36 \times 10^{-4}$ | $4.94 \times 10^{-4}$ | $0.62 \times 10^{-2}$ | $8.53 \times 10^{-4}$ | $2.72 \times 10^{-3}$ | $2.79 \times 10^7$ | $3.37 \times 10^4$ | $4.34 \times 10^8$ | C |
| 50 | $1.14 \times 10^{-4}$ | 0.999 | $9.28 \times 10^{-5}$ | $5.12 \times 10^{-4}$ | $8.28 \times 10^{-2}$ | $1.67 \times 10^{-4}$ | $1.12 \times 10^{-2}$ | $5.52 \times 10^6$ | $0.51 \times 10^3$ | $7.36 \times 10^6$ | B |
| 52 | $1.14 \times 10^{-4}$ | 0.999 | $1.35 \times 10^{-4}$ | $4.99 \times 10^{-4}$ | $1.05 \times 10^{-2}$ | $7.44 \times 10^{-4}$ | $2.87 \times 10^{-3}$ | $1.50 \times 10^7$ | $2.47 \times 10^4$ | $1.53 \times 10^8$ | C |
| 56 | $1.14 \times 10^{-4}$ | 0.999 | $1.35 \times 10^{-4}$ | $5.07 \times 10^{-4}$ | $1.04 \times 10^{-2}$ | $7.45 \times 10^{-4}$ | $5.31 \times 10^{-3}$ | $2.17 \times 10^7$ | $4.38 \times 10^4$ | $1.60 \times 10^8$ | C |
| **PEG IFN- $\alpha$ -treated patient (HBeAg-negative PVR)** |  |  |  |  |  |  |  |  |  |  |  |
| 5 | $1.14 \times 10^{-4}$ | 0.999 | $1.36 \times 10^{-4}$ | $5.12 \times 10^{-4}$ | $1.05 \times 10^{-1}$ | $9.31 \times 10^{-4}$ | $1.48 \times 10^{-2}$ | $2.05 \times 10^6$ | $0.66 \times 10^3$ | $1.74 \times 10^4$ | C |
| 9 | $1.14 \times 10^{-4}$ | 0.999 | $1.32 \times 10^{-4}$ | $5.12 \times 10^{-4}$ | $8.27 \times 10^{-2}$ | $6.08 \times 10^{-4}$ | $2.08 \times 10^{-2}$ | $1.28 \times 10^5$ | $0.09 \times 10^3$ | $4.50 \times 10^5$ | C |
| 11 | $1.14 \times 10^{-4}$ | 0.999 | $1.30 \times 10^{-4}$ | $5.11 \times 10^{-4}$ | $3.19 \times 10^{-2}$ | $5.43 \times 10^{-4}$ | $2.34 \times 10^{-2}$ | $2.53 \times 10^7$ | $5.10 \times 10^3$ | $9.14 \times 10^6$ | C |
| 31 | $1.14 \times 10^{-4}$ | 0.999 | $1.36 \times 10^{-4}$ | $5.07 \times 10^{-4}$ | $5.14 \times 10^{-2}$ | $8.73 \times 10^{-4}$ | $3.78 \times 10^{-3}$ | $5.05 \times 10^4$ | $8.04 \times 10^3$ | $3.37 \times 10^3$ | C |
| 43 | $1.14 \times 10^{-4}$ | 0.999 | $9.36 \times 10^{-5}$ | $5.12 \times 10^{-4}$ | $1.94 \times 10^{-1}$ | $1.71 \times 10^{-4}$ | $1.85 \times 10^{-2}$ | $3.27 \times 10^5$ | $3.36 \times 10^3$ | $9.18 \times 10^4$ | C |
| 45 | $1.14 \times 10^{-4}$ | 0.999 | $1.34 \times 10^{-4}$ | $5.11 \times 10^{-4}$ | $8.59 \times 10^{-1}$ | $6.90 \times 10^{-4}$ | $3.72 \times 10^{-2}$ | $9.39 \times 10^4$ | $1.32 \times 10^3$ | $1.95 \times 10^3$ | C |
| 64 | $1.14 \times 10^{-4}$ | 0.999 | $1.34 \times 10^{-4}$ | $5.11 \times 10^{-4}$ | $3.12 \times 10^{-1}$ | $7.06 \times 10^{-4}$ | $3.55 \times 10^{-2}$ | $8.56 \times 10^4$ | $0.94 \times 10^3$ | $4.01 \times 10^3$ | C |
| 65 | $1.14 \times 10^{-4}$ | 0.999 | $1.38 \times 10^{-4}$ | $5.11 \times 10^{-4}$ | $2.85 \times 10^{-1}$ | $1.68 \times 10^{-3}$ | $5.49 \times 10^{-3}$ | $4.74 \times 10^4$ | $1.03 \times 10^3$ | $1.40 \times 10^3$ | C |
| 67 | $1.14 \times 10^{-4}$ | 0.999 | $1.35 \times 10^{-4}$ | $5.04 \times 10^{-4}$ | $2.26 \times 10^{-1}$ | $8.03 \times 10^{-4}$ | $3.18 \times 10^{-3}$ | $2.28 \times 10^5$ | $1.63 \times 10^3$ | $0.51 \times 10^3$ | C |
| 68 | $1.14 \times 10^{-4}$ | 0.999 | $1.15 \times 10^{-4}$ | $5.12 \times 10^{-4}$ | $1.98 \times 10^{-1}$ | $3.01 \times 10^{-4}$ | $1.73 \times 10^{-2}$ | $1.79 \times 10^6$ | $0.05 \times 10^3$ | $3.99 \times 10^3$ | B |
| 76 | $1.14 \times 10^{-4}$ | 0.999 | $1.38 \times 10^{-4}$ | $5.12 \times 10^{-4}$ | $3.46 \times 10^{-1}$ | $2.74 \times 10^{-3}$ | $1.67 \times 10^{-2}$ | $3.04 \times 10^5$ | $1.06 \times 10^3$ | $2.60 \times 10^4$ | C |
| 77 | $1.14 \times 10^{-4}$ | 0.999 | $1.38 \times 10^{-4}$ | $5.12 \times 10^{-4}$ | $2.64 \times 10^{-1}$ | $2.10 \times 10^{-3}$ | $8.80 \times 10^{-3}$ | $1.44 \times 10^6$ | $0.62 \times 10^3$ | $6.63 \times 10^5$ | C |
| 78 | $1.14 \times 10^{-4}$ | 0.999 | $1.38 \times 10^{-4}$ | $5.10 \times 10^{-4}$ | $2.45 \times 10^{-1}$ | $1.74 \times 10^{-3}$ | $4.98 \times 10^{-3}$ | $4.66 \times 10^4$ | $0.64 \times 10^3$ | $4.42 \times 10^4$ | C |
| 79 | $1.14 \times 10^{-4}$ | 0.999 | $1.38 \times 10^{-4}$ | $5.11 \times 10^{-4}$ | $5.23 \times 10^{-2}$ | $1.39 \times 10^{-3}$ | $5.40 \times 10^{-3}$ | $1.56 \times 10^6$ | $0.71 \times 10^3$ | $1.96 \times 10^5$ | C |
| 83 | $1.14 \times 10^{-4}$ | 0.999 | $1.38 \times 10^{-4}$ | $5.13 \times 10^{-4}$ | $1.60 \times 10^{-1}$ | $1.42 \times 10^{-3}$ | $1.02 \times 10^{-2}$ | $5.75 \times 10^4$ | $0.30 \times 10^3$ | $9.08 \times 10^4$ | C |
| 102 | $1.14 \times 10^{-4}$ | 0.999 | $1.29 \times 10^{-4}$ | $5.11 \times 10^{-4}$ | $2.31 \times 10^{-1}$ | $5.04 \times 10^{-4}$ | $5.97 \times 10^{-3}$ | $1.57 \times 10^4$ | $1.87 \times 10^3$ | $0.80 \times 10^3$ | B |
| 105 | $1.14 \times 10^{-4}$ | 0.999 | $1.38 \times 10^{-4}$ | $5.12 \times 10^{-4}$ | $2.80 \times 10^{-1}$ | $1.46 \times 10^{-3}$ | $1.74 \times 10^{-2}$ | $1.07 \times 10^5$ | $1.46 \times 10^3$ | $2.92 \times 10^5$ | C |
| 108 | $1.14 \times 10^{-4}$ | 0.999 | $1.38 \times 10^{-4}$ | $5.11 \times 10^{-4}$ | $3.47 \times 10^{-1}$ | $1.15 \times 10^{-3}$ | $6.24 \times 10^{-3}$ | $1.25 \times 10^5$ | $1.80 \times 10^3$ | $4.28 \times 10^4$ | C |
| 109 | $1.14 \times 10^{-4}$ | 0.999 | $1.23 \times 10^{-4}$ | $5.12 \times 10^{-4}$ | $3.29 \times 10^{-1}$ | $3.92 \times 10^{-4}$ | $1.90 \times 10^{-2}$ | $1.94 \times 10^5$ | $0.22 \times 10^3$ | $2.78 \times 10^4$ | C |
| 114 | $1.14 \times 10^{-4}$ | 0.999 | $1.27 \times 10^{-4}$ | $5.12 \times 10^{-4}$ | $2.57 \times 10^{-1}$ | $4.52 \times 10^{-4}$ | $1.42 \times 10^{-2}$ | $2.97 \times 10^5$ | $0.65 \times 10^3$ | $6.03 \times 10^3$ | C |
| 121 | $1.14 \times 10^{-4}$ | 0.999 | $1.22 \times 10^{-4}$ | $5.12 \times 10^{-4}$ | $3.07 \times 10^{-1}$ | $3.83 \times 10^{-4}$ | $1.90 \times 10^{-2}$ | $1.66 \times 10^5$ | $0.30 \times 10^3$ | $3.43 \times 10^3$ | B |
| J20 | $1.14 \times 10^{-4}$ | 0.999 | $1.37 \times 10^{-4}$ | $5.13 \times 10^{-4}$ | $1.53 \times 10^{-1}$ | $4.88 \times 10^{-3}$ | $1.37 \times 10^{-2}$ | $1.84 \times 10^8$ | $2.38 \times 10^3$ | $3.12 \times 10^7$ | --- |
| J25 | $1.14 \times 10^{-4}$ | 0.999 | $1.19 \times 10^{-4}$ | $4.81 \times 10^{-4}$ | $1.98 \times 10^{-1}$ | $3.39 \times 10^{-4}$ | $1.63 \times 10^{-3}$ | $2.81 \times 10^6$ | $4.72 \times 10^3$ | $2.75 \times 10^5$ | --- |
| J38 | $1.14 \times 10^{-4}$ | 0.999 | $9.31 \times 10^{-5}$ | $4.99 \times 10^{-4}$ | $8.18 \times 10^{-1}$ | $1.69 \times 10^{-4}$ | $2.66 \times 10^{-3}$ | $6.81 \times 10^4$ | $5.45 \times 10^3$ | $1.62 \times 10^5$ | --- |
| J40 | $1.14 \times 10^{-4}$ | 0.999 | $9.03 \times 10^{-5}$ | $4.93 \times 10^{-4}$ | $9.20 \times 10^{-2}$ | $1.58 \times 10^{-4}$ | $2.13 \times 10^{-3}$ | $2.75 \times 10^5$ | $1.52 \times 10^4$ | $1.04 \times 10^4$ | --- |
| J77 | $1.14 \times 10^{-4}$ | 0.999 | $1.32 \times 10^{-4}$ | $5.08 \times 10^{-4}$ | $0.96 \times 10^{-2}$ | $7.02 \times 10^{-4}$ | $1.03 \times 10^{-2}$ | $1.20 \times 10^3$ | $0.03 \times 10^3$ | $3.39 \times 10^3$ | --- |
| J79 | $1.14 \times 10^{-4}$ | 0.999 | $1.31 \times 10^{-4}$ | $5.10 \times 10^{-4}$ | $7.67 \times 10^{-1}$ | $1.83 \times 10^{-3}$ | $1.15 \times 10^{-2}$ | $4.36 \times 10^3$ | $0.07 \times 10^2$ | $4.38 \times 10^3$ | --- |
| J80 | $1.14 \times 10^{-4}$ | 0.999 | $1.38 \times 10^{-4}$ | $5.12 \times 10^{-4}$ | $2.47 \times 10^{-1}$ | $3.29 \times 10^{-4}$ | $8.28 \times 10^{-3}$ | $7.56 \times 10^4$ | $0.07 \times 10^3$ | $3.03 \times 10^3$ | --- |
| **PEG IFN- $\alpha$ and NAS treated patient (HBeAg-negative PVR)** |  |  |  |  |  |  |  |  |  |  |  |
| 1 | $1.14 \times 10^{-4}$ | 0.999 | $1.36 \times 10^{-4}$ | $5.11 \times 10^{-4}$ | $2.30 \times 10^{-1}$ | $8.78 \times 10^{-4}$ | $6.14 \times 10^{-3}$ | $1.24 \times 10^5$ | $0.23 \times 10^3$ | $3.78 \times 10^2$ | B |
| 6 | $1.14 \times 10^{-4}$ | 0.999 | $1.38 \times 10^{-4}$ | $5.12 \times 10^{-4}$ | $1.10 \times 10^{-1}$ | $2.56 \times 10^{-3}$ | $2.14 \times 10^{-2}$ | $2.45 \times 10^6$ | $2.57 \times 10^3$ | $2.83 \times 10^5$ | C |
| 7 | $1.14 \times 10^{-4}$ | 0.999 | $1.31 \times 10^{-4}$ | $5.12 \times 10^{-4}$ | $2.81 \times 10^{-1}$ | $5.55 \times 10^{-4}$ | $1.16 \times 10^{-2}$ | $1.02 \times 10^5$ | $0.32 \times 10^3$ | $8.45 \times 10^4$ | C |
| 25 | $1.14 \times 10^{-4}$ | 0.999 | $1.35 \times 10^{-4}$ | $5.11 \times 10^{-4}$ | $2.86 \times 10^{-1}$ | $8.10 \times 10^{-4}$ | $6.33 \times 10^{-3}$ | $9.22 \times 10^5$ | $2.30 \times 10^3$ | $0.68 \times 10^3$ | C |
| 54 | $1.14 \times 10^{-4}$ | 0.999 | $1.38 \times 10^{-4}$ | $5.12 \times 10^{-4}$ | $3.10 \times 10^{-1}$ | $1.47 \times 10^{-3}$ | $7.63 \times 10^{-3}$ | $5.13 \times 10^4$ | $0.59 \times 10^3$ | $1.14 \times 10^3$ | C |
| 56 | $1.14 \times 10^{-4}$ | 0.999 | $1.37 \times 10^{-4}$ | $512 \times 10^{-4}$ | $2.99 \times 10^{-1}$ | $1.03 \times 10^{-3}$ | $1.80 \times 10^{-2}$ | $6.38 \times 10^4$ | $0.06 \times 10^3$ | $2.64 \times 10^3$ | B |
| 60 | $1.14 \times 10^{-4}$ | 0.999 | $1.38 \times 10^{-4}$ | $512 \times 10^{-4}$ | $2.76 \times 10^{-1}$ | $1.71 \times 10^{-3}$ | $1.24 \times 10^{-2}$ | $9.63 \times 10^5$ | $0.59 \times 10^3$ | $3.99 \times 10^3$ | C |
| 72 | $1.14 \times 10^{-4}$ | 0.999 | $1.38 \times 10^{-4}$ | $4.98 \times 10^{-4}$ | $3.03 \times 10^{-1}$ | $1.80 \times 10^{-3}$ | $2.48 \times 10^{-3}$ | $1.12 \times 10^5$ | $2.59 \times 10^3$ | $0.27 \times 10^3$ | C |
| 75 | $1.14 \times 10^{-4}$ | 0.999 | $1.38 \times 10^{-4}$ | $5.13 \times 10^{-4}$ | $2.09 \times 10^{-1}$ | $1.80 \times 10^{-3}$ | $1.05 \times 10^{-2}$ | $9.26 \times 10^5$ | $0.73 \times 10^3$ | $3.05 \times 10^3$ | C |
| 95 | $1.14 \times 10^{-4}$ | 0.999 | $1.36 \times 10^{-4}$ | $5.11 \times 10^{-4}$ | $1.09 \times 10^{-1}$ | $1.06 \times 10^{-2}$ | $5.69 \times 10^{-3}$ | $1.44 \times 10^6$ | $1.07 \times 10^4$ | $1.61 \times 10^6$ | C |
| 96 | $1.14 \times 10^{-4}$ | 0.999 | $1.38 \times 10^{-4}$ | $5.11 \times 10^{-4}$ | $3.16 \times 10^{-1}$ | $3.45 \times 10^{-3}$ | $5.39 \times 10^{-3}$ | $1.25 \times 10^5$ | $1.67 \times 10^3$ | $0.32 \times 10^3$ | C |
| 104 | $1.14 \times 10^{-4}$ | 0.999 | $1.38 \times 10^{-4}$ | $5.10 \times 10^{-4}$ | $2.60 \times 10^{-1}$ | $2.73 \times 10^{-3}$ | $4.82 \times 10^{-3}$ | $8.46 \times 10^4$ | $2.24 \times 10^3$ | $3.26 \times 10^3$ | C |
| 110 | $1.14 \times 10^{-4}$ | 0.999 | $1.36 \times 10^{-4}$ | $5.08 \times 10^{-4}$ | $3.23 \times 10^{-1}$ | $1.03 \times 10^{-3}$ | $1.95 \times 10^{-2}$ | $6.76 \times 10^4$ | $3.87 \times 10^3$ | $6.78 \times 10^3$ | C |
| 119 | $1.14 \times 10^{-4}$ | 0.999 | $1.38 \times 10^{-4}$ | $5.01 \times 10^{-4}$ | $9.86 \times 10^{-2}$ | $2.15 \times 10^{-3}$ | $2.72 \times 10^{-3}$ | $6.76 \times 10^4$ | $2.41 \times 10^3$ | $2.47 \times 10^5$ | C |
| **PEG IFN-α treated patient (HBeAg-negative non-PVR)** |  |  |  |  |  |  |  |  |  |  |  |
| 2 | $1.14 \times 10^{-4}$ | 0.999 | $1.18 \times 10^{-4}$ | $4.98 \times 10^{-4}$ | $2.43 \times 10^{-1}$ | $3.29 \times 10^{-4}$ | $2.48 \times 10^{-3}$ | $1.63 \times 10^5$ | $6.13 \times 10^3$ | $5.76 \times 10^4$ | C |
| 3 | $1.14 \times 10^{-4}$ | 0.999 | $1.06 \times 10^{-4}$ | $4.99 \times 10^{-5}$ | $6.31 \times 10^{-2}$ | $2.30 \times 10^{-4}$ | $2.64 \times 10^{-3}$ | $2.42 \times 10^5$ | $0.74 \times 10^3$ | $1.81 \times 10^3$ | B |
| 12 | $1.14 \times 10^{-4}$ | 0.999 | $1.08 \times 10^{-4}$ | $4.99 \times 10^{-5}$ | $3.72 \times 10^{-1}$ | $2.45 \times 10^{-4}$ | $2.61 \times 10^{-3}$ | $3.13 \times 10^5$ | $6.40 \times 10^3$ | $1.52 \times 10^4$ | C |
| 14 | $1.14 \times 10^{-4}$ | 0.999 | $9.80 \times 10^{-5}$ | $5.06 \times 10^{-5}$ | $5.67 \times 10^{-2}$ | $1.90 \times 10^{-4}$ | $3.84 \times 10^{-3}$ | $1.45 \times 10^5$ | $0.20 \times 10^3$ | $1.33 \times 10^3$ | B |
| 15 | $1.14 \times 10^{-4}$ | 0.999 | $1.14 \times 10^{-4}$ | $4.97 \times 10^{-5}$ | $3.87 \times 10^{-1}$ | $2.90 \times 10^{-4}$ | $2.40 \times 10^{-3}$ | $1.58 \times 10^6$ | $2.90 \times 10^4$ | $5.30 \times 10^3$ | C |
| 18 | $1.14 \times 10^{-4}$ | 0.999 | $1.31 \times 10^{-4}$ | $5.11 \times 10^{-4}$ | $2.31 \times 10^{-1}$ | $5.58 \times 10^{-4}$ | $1.55 \times 10^{-2}$ | $1.97 \times 10^6$ | $2.97 \times 10^3$ | $5.53 \times 10^4$ | B |
| 20 | $1.14 \times 10^{-4}$ | 0.999 | $1.37 \times 10^{-4}$ | $5.05 \times 10^{-4}$ | $2.69 \times 10^{-1}$ | $4.26 \times 10^{-3}$ | $3.30 \times 10^{-3}$ | $3.41 \times 10^5$ | $3.49 \times 10^4$ | $2.98 \times 10^4$ | C |
| 22 | $1.14 \times 10^{-4}$ | 0.999 | $1.09 \times 10^{-4}$ | $5.00 \times 10^{-4}$ | $639 \times 10^{-2}$ | $2.54 \times 10^{-4}$ | $2.71 \times 10^{-3}$ | $1.04 \times 10^6$ | $4.03 \times 10^3$ | $1.40 \times 10^4$ | B |
| 24 | $1.14 \times 10^{-4}$ | 0.999 | $7.36 \times 10^{-5}$ | $5.12 \times 10^{-4}$ | $2.08 \times 10^{-1}$ | $1.07 \times 10^{-4}$ | $7.07 \times 10^{-3}$ | $8.69 \times 10^5$ | $0.66 \times 10^3$ | $3.04 \times 10^4$ | C |
| 26 | $1.14 \times 10^{-4}$ | 0.999 | $1.30 \times 10^{-4}$ | $5.11 \times 10^{-4}$ | $2.81 \times 10^{-1}$ | $5.43 \times 10^{-4}$ | $6.34 \times 10^{-3}$ | $6.27 \times 10^5$ | $8.81 \times 10^3$ | $1.06 \times 10^3$ | C |
| 29 | $1.14 \times 10^{-4}$ | 0.999 | $1.34 \times 10^{-4}$ | $5.01 \times 10^{-4}$ | $1.56 \times 10^{-2}$ | $7.13 \times 10^{-4}$ | $2.98 \times 10^{-3}$ | $2.73 \times 10^5$ | $2.69 \times 10^3$ | $1.82 \times 10^5$ | C |
| 32 | $1.14 \times 10^{-4}$ | 0.999 | $1.00 \times 10^{-4}$ | $5.04 \times 10^{-4}$ | $2.42 \times 10^{-1}$ | $2.00 \times 10^{-4}$ | $3.16 \times 10^{-3}$ | $9.67 \times 10^5$ | $1.64 \times 10^3$ | $2.76 \times 10^3$ | C |
| 34 | $1.14 \times 10^{-4}$ | 0.999 | $1.37 \times 10^{-4}$ | $5.13 \times 10^{-4}$ | $8.36 \times 10^{-2}$ | $5.39 \times 10^{-3}$ | $1.05 \times 10^{-2}$ | $1.10 \times 10^7$ | $6.57 \times 10^3$ | $9.60 \times 10^4$ | C |
| 37 | $1.14 \times 10^{-4}$ | 0.999 | $1.36 \times 10^{-4}$ | $4.88 \times 10^{-4}$ | $0.34 \times 10^{-2}$ | $8.62 \times 10^{-4}$ | $2.99 \times 10^{-3}$ | $2.31 \times 10^6$ | $3.84 \times 10^3$ | $3.94 \times 10^5$ | C |
| 38 | $1.14 \times 10^{-4}$ | 0.999 | $1.38 \times 10^{-4}$ | $4.96 \times 10^{-4}$ | $8.45 \times 10^{-2}$ | $1.82 \times 10^{-3}$ | $2.32 \times 10^{-3}$ | $4.97 \times 10^4$ | $1.59 \times 10^3$ | $2.57 \times 10^4$ | C |
| 41 | $1.14 \times 10^{-4}$ | 0.999 | $7.66 \times 10^{-5}$ | $4.98 \times 10^{-4}$ | $8.15 \times 10^{-2}$ | $1.15 \times 10^{-4}$ | $2.55 \times 10^{-3}$ | $4.55 \times 10^5$ | $0.77 \times 10^3$ | $1.16 \times 10^3$ | B |
| 48 | $1.14 \times 10^{-4}$ | 0.999 | $9.96 \times 10^{-5}$ | $5.05 \times 10^{-4}$ | $2.15 \times 10^{-1}$ | $1.97 \times 10^{-4}$ | $3.47 \times 10^{-3}$ | $1.64 \times 10^5$ | $1.31 \times 10^3$ | $0.86 \times 10^3$ | B |
| 49 | $1.14 \times 10^{-4}$ | 0.999 | $1.00 \times 10^{-4}$ | $4.93 \times 10^{-4}$ | $3.64 \times 10^{-2}$ | $2.00 \times 10^{-4}$ | $2.17 \times 10^{-3}$ | $6.70 \times 10^5$ | $5.56 \times 10^3$ | $2.05 \times 10^4$ | B |
| 51 | $1.14 \times 10^{-4}$ | 0.999 | $8.10 \times 10^{-5}$ | $5.02 \times 10^{-4}$ | $2.76 \times 10^{-1}$ | $1.27 \times 10^{-4}$ | $2.92 \times 10^{-3}$ | $1.69 \times 10^6$ | $2.95 \times 10^3$ | $1.88 \times 10^5$ | C |
| 52 | $1.14 \times 10^{-4}$ | 0.999 | $1.34 \times 10^{-4}$ | $4.98 \times 10^{-4}$ | $0.55 \times 10^{-2}$ | $7.27 \times 10^{-4}$ | $3.52 \times 10^{-3}$ | $7.60 \times 10^4$ | $7.22 \times 10^3$ | $1.07 \times 10^5$ | B |
| 53 | $1.14 \times 10^{-4}$ | 0.999 | $1.34 \times 10^{-4}$ | $5.00 \times 10^{-4}$ | $0.80 \times 10^{-2}$ | $7.22 \times 10^{-4}$ | $3.34 \times 10^{-3}$ | $8.72 \times 10^6$ | $3.21 \times 10^3$ | $7.09 \times 10^6$ | C |
| 55 | $1.14 \times 10^{-4}$ | 0.999 | $1.38 \times 10^{-4}$ | $4.73 \times 10^{-4}$ | $2.97 \times 10^{-1}$ | $2.21 \times 10^{-3}$ | $1.40 \times 10^{-3}$ | $4.00 \times 10^4$ | $0.92 \times 10^3$ | $1.18 \times 10^4$ | C |
| 58 | $1.14 \times 10^{-4}$ | 0.999 | $1.36 \times 10^{-4}$ | $4.94 \times 10^{-4}$ | $2.13 \times 10^{-1}$ | $8.83 \times 10^{-4}$ | $2.20 \times 10^{-3}$ | $1.78 \times 10^5$ | $1.97 \times 10^3$ | $2.41 \times 10^4$ | C |
| 63 | $1.14 \times 10^{-4}$ | 0.999 | $1.38 \times 10^{-4}$ | $5.06 \times 10^{-4}$ | $1.93 \times 10^{-1}$ | $1.40 \times 10^{-3}$ | $3.50 \times 10^{-3}$ | $1.01 \times 10^5$ | $7.31 \times 10^3$ | $5.75 \times 10^3$ | C |
| 71 | $1.14 \times 10^{-4}$ | 0.999 | $1.37 \times 10^{-4}$ | $4.98 \times 10^{-4}$ | $3.04 \times 10^{-1}$ | $9.34 \times 10^{-4}$ | $2.66 \times 10^{-3}$ | $1.71 \times 10^5$ | $2.37 \times 10^3$ | $1.49 \times 10^3$ | B |
| 74 | $1.14 \times 10^{-4}$ | 0.999 | $1.38 \times 10^{-4}$ | $5.01 \times 10^{-4}$ | $1.75 \times 10^{-1}$ | $2.56 \times 10^{-3}$ | $2.81 \times 10^{-3}$ | $5.25 \times 10^5$ | $3.29 \times 10^4$ | $6.86 \times 10^3$ | C |
| 84 | $1.14 \times 10^{-4}$ | 0.999 | $1.38 \times 10^{-4}$ | $5.12 \times 10^{-4}$ | $1.59 \times 10^{-1}$ | $1.49 \times 10^{-3}$ | $8.14 \times 10^{-3}$ | $5.52 \times 10^4$ | $1.39 \times 10^3$ | $5.96 \times 10^3$ | C |
| 86 | $1.14 \times 10^{-4}$ | 0.999 | $1.33 \times 10^{-4}$ | $5.10 \times 10^{-4}$ | $1.23 \times 10^{-2}$ | $6.47 \times 10^{-4}$ | $1.03 \times 10^{-2}$ | $1.16 \times 10^7$ | $1.92 \times 10^4$ | $2.89 \times 10^7$ | C |
| 88 | $1.14 \times 10^{-4}$ | 0.999 | $1.36 \times 10^{-4}$ | $5.13 \times 10^{-4}$ | $1.12 \times 10^{-1}$ | $1.33 \times 10^{-2}$ | $9.25 \times 10^{-3}$ | $5.20 \times 10^6$ | $3.58 \times 10^3$ | $1.36 \times 10^4$ | C |
| 89 | $1.14 \times 10^{-4}$ | 0.999 | $1.28 \times 10^{-4}$ | $5.08 \times 10^{-4}$ | $4.02 \times 10^{-1}$ | $4.84 \times 10^{-4}$ | $4.27 \times 10^{-3}$ | $1.66 \times 10^5$ | $1.81 \times 10^3$ | $5.01 \times 10^4$ | C |
| 90 | $1.14 \times 10^{-4}$ | 0.999 | $1.38 \times 10^{-4}$ | $4.99 \times 10^{-4}$ | $2.14 \times 10^{-1}$ | $2.03 \times 10^{-3}$ | $2.57 \times 10^{-3}$ | $4.26 \times 10^4$ | $3.07 \times 10^3$ | $0.47 \times 10^3$ | C |
| 92 | $1.14 \times 10^{-4}$ | 0.999 | $1.31 \times 10^{-4}$ | $4.92 \times 10^{-4}$ | $3.27 \times 10^{-1}$ | $5.88 \times 10^{-4}$ | $2.08 \times 10^{-3}$ | $1.17 \times 10^5$ | $1.67 \times 10^3$ | $2.09 \times 10^3$ | B |
| 98 | $1.14 \times 10^{-4}$ | 0.999 | $1.33 \times 10^{-4}$ | $5.05 \times 10^{-4}$ | $1.23 \times 10^{-2}$ | $6.34 \times 10^{-4}$ | $3.93 \times 10^{-3}$ | $3.40 \times 10^4$ | $2.34 \times 10^3$ | $1.69 \times 10^5$ | C |
| 100 | $1.14 \times 10^{-4}$ | 0.999 | $1.26 \times 10^{-4}$ | $5.10 \times 10^{-4}$ | $3.65 \times 10^{-1}$ | $4.38 \times 10^{-4}$ | $4.77 \times 10^{-3}$ | $1.83 \times 10^6$ | $0.70 \times 10^3$ | $9.29 \times 10^3$ | B |
| 101 | $1.14 \times 10^{-4}$ | 0.999 | $1.38 \times 10^{-4}$ | $4.99 \times 10^{-4}$ | $2.26 \times 10^{-1}$ | $2.21 \times 10^{-3}$ | $2.55 \times 10^{-3}$ | $7.36 \times 10^6$ | $3.46 \times 10^3$ | $1.62 \times 10^6$ | B |
| 103 | $1.14 \times 10^{-4}$ | 0.999 | $1.38 \times 10^{-4}$ | $4.89 \times 10^{-4}$ | $2.38 \times 10^{-1}$ | $3.05 \times 10^{-3}$ | $1.92 \times 10^{-3}$ | $5.02 \times 10^5$ | $2.15 \times 10^4$ | $5.26 \times 10^3$ | C |
| 112 | $1.14 \times 10^{-4}$ | 0.999 | $1.36 \times 10^{-4}$ | $4.96 \times 10^{-4}$ | $0.98 \times 10^{-2}$ | $8.30 \times 10^{-4}$ | $2.59 \times 10^{-3}$ | $6.47 \times 10^6$ | $5.09 \times 10^3$ | $2.60 \times 10^6$ | C |
| 117 | $1.14 \times 10^{-4}$ | 0.999 | $1.36 \times 10^{-4}$ | $5.10 \times 10^{-4}$ | $2.87 \times 10^{-1}$ | $8.63 \times 10^{-4}$ | $4.78 \times 10^{-3}$ | $7.37 \times 10^4$ | $4.75 \times 10^3$ | $1.60 \times 10^4$ | C |
| 122 | $1.14 \times 10^{-4}$ | 0.999 | $1.25 \times 10^{-4}$ | $4.84 \times 10^{-4}$ | $3.37 \times 10^{-1}$ | $4.16 \times 10^{-4}$ | $1.72 \times 10^{-3}$ | $3.84 \times 10^5$ | $3.46 \times 10^3$ | $3.48 \times 10^4$ | C |
| 124 | $1.14 \times 10^{-4}$ | 0.999 | $1.38 \times 10^{-4}$ | $4.89 \times 10^{-4}$ | $2.30 \times 10^{-1}$ | $1.13 \times 10^{-3}$ | $1.94 \times 10^{-3}$ | $6.27 \times 10^4$ | $1.25 \times 10^4$ | $2.45 \times 10^3$ | C |
| 125 | $1.14 \times 10^{-4}$ | 0.999 | $1.38 \times 10^{-4}$ | $4.91 \times 10^{-4}$ | $9.75 \times 10^{-2}$ | $3.23 \times 10^{-3}$ | $2.01 \times 10^{-3}$ | $9.74 \times 10^4$ | $6.79 \times 10^3$ | $0.82 \times 10^3$ | C |
| J18 | $1.14 \times 10^{-4}$ | 0.999 | $1.26 \times 10^{-4}$ | $4.67 \times 10^{-4}$ | $9.85 \times 10^{-2}$ | $4.33 \times 10^{-4}$ | $1.28 \times 10^{-3}$ | $7.57 \times 10^7$ | $2.75 \times 10^4$ | $6.24 \times 10^5$ | --- |
| J19 | $1.14 \times 10^{-4}$ | 0.999 | $1.31 \times 10^{-4}$ | $5.11 \times 10^{-4}$ | $2.12 \times 10^{-2}$ | $6.04 \times 10^{-4}$ | $1.12 \times 10^{-2}$ | $3.58 \times 10^7$ | $2.32 \times 10^4$ | $8.92 \times 10^7$ | --- |
| J21 | $1.14 \times 10^{-4}$ | 0.999 | $1.20 \times 10^{-4}$ | $5.03 \times 10^{-4}$ | $2.74 \times 10^{-1}$ | $3.55 \times 10^{-4}$ | $3.01 \times 10^{-3}$ | $6.34 \times 10^5$ | $2.51 \times 10^3$ | $4.80 \times 10^3$ | --- |
| J22 | $1.14 \times 10^{-4}$ | 0.999 | $1.11 \times 10^{-4}$ | $4.81 \times 10^{-4}$ | $3.80 \times 10^{-1}$ | $2.68 \times 10^{-4}$ | $1.61 \times 10^{-3}$ | $6.98 \times 10^5$ | $1.13 \times 10^4$ | $1.13 \times 10^3$ | --- |
| J23 | $1.14 \times 10^{-4}$ | 0.999 | $1.20 \times 10^{-4}$ | $4.95 \times 10^{-4}$ | $2.73 \times 10^{-1}$ | $3.47 \times 10^{-4}$ | $2.24 \times 10^{-3}$ | $7.77 \times 10^5$ | $4.75 \times 10^3$ | $2.20 \times 10^5$ | --- |
| J26 | $1.14 \times 10^{-4}$ | 0.999 | $1.34 \times 10^{-4}$ | $5.00 \times 10^{-4}$ | $0.81 \times 10^{-2}$ | $6.91 \times 10^{-4}$ | $3.25 \times 10^{-3}$ | $3.33 \times 10^6$ | $0.19 \times 10^3$ | $7.80 \times 10^6$ | --- |
| J27 | $1.14 \times 10^{-4}$ | 0.999 | $1.08 \times 10^{-4}$ | $4.86 \times 10^{-4}$ | $8.73 \times 10^{-2}$ | $2.45 \times 10^{-4}$ | $1.80 \times 10^{-3}$ | $2.16 \times 10^8$ | $9.31 \times 10^3$ | $9.84 \times 10^6$ | --- |
| J28 | $1.14 \times 10^{-4}$ | 0.999 | $8.58 \times 10^{-5}$ | $5.11 \times 10^{-4}$ | $2.31 \times 10^{-1}$ | $1.42 \times 10^{-4}$ | $6.17 \times 10^{-3}$ | $1.46 \times 10^6$ | $7.02 \times 10^3$ | $1.32 \times 10^4$ | --- |
| J32 | $1.14 \times 10^{-4}$ | 0.999 | $1.36 \times 10^{-4}$ | $5.12 \times 10^{-4}$ | $3.28 \times 10^{-1}$ | $7.90 \times 10^{-4}$ | $7.38 \times 10^{-3}$ | $2.08 \times 10^8$ | $1.92 \times 10^3$ | $4.59 \times 10^6$ | --- |
| J33 | $1.14 \times 10^{-4}$ | 0.999 | $1.38 \times 10^{-4}$ | $4.83 \times 10^{-4}$ | $0.67 \times 10^{-2}$ | $1.31 \times 10^{-3}$ | $1.88 \times 10^{-3}$ | $4.31 \times 10^8$ | $2.48 \times 10^4$ | $1.51 \times 10^8$ | --- |
| J34 | $1.14 \times 10^{-4}$ | 0.999 | $1.10 \times 10^{-4}$ | $4.87 \times 10^{-4}$ | $4.32 \times 10^{-1}$ | $2.56 \times 10^{-4}$ | $1.83 \times 10^{-3}$ | $1.49 \times 10^6$ | $1.92 \times 10^4$ | $5.31 \times 10^3$ | --- |
| J35 | $1.14 \times 10^{-4}$ | 0.999 | $1.34 \times 10^{-4}$ | $4.95 \times 10^{-4}$ | $9.01 \times 10^{-2}$ | $7.25 \times 10^{-4}$ | $2.58 \times 10^{-3}$ | $1.42 \times 10^8$ | $1.82 \times 10^4$ | $5.19 \times 10^7$ | --- |
| J37 | $1.14 \times 10^{-4}$ | 0.999 | $1.34 \times 10^{-4}$ | $4.94 \times 10^{-4}$ | $0.57 \times 10^{-2}$ | $7.42 \times 10^{-4}$ | $2.83 \times 10^{-3}$ | $1.29 \times 10^8$ | $2.45 \times 10^4$ | $2.35 \times 10^8$ | --- |
| J39 | $1.14 \times 10^{-4}$ | 0.999 | $1.38 \times 10^{-4}$ | $4.82 \times 10^{-4}$ | $0.53 \times 10^{-2}$ | $1.51 \times 10^{-3}$ | $1.95 \times 10^{-3}$ | $2.30 \times 10^8$ | $4.63 \times 10^4$ | $6.00 \times 10^8$ | --- |
| J53 | $1.14 \times 10^{-4}$ | 0.999 | $1.13 \times 10^{-4}$ | $5.11 \times 10^{-4}$ | $3.90 \times 10^{-1}$ | $2.81 \times 10^{-4}$ | $6.25 \times 10^{-3}$ | $1.35 \times 10^6$ | $5.69 \times 10^2$ | $0.66 \times 10^3$ | --- |
| J54 | $1.14 \times 10^{-4}$ | 0.999 | $1.38 \times 10^{-4}$ | $4.87 \times 10^{-4}$ | $0.92 \times 10^{-2}$ | $1.45 \times 10^{-3}$ | $2.00 \times 10^{-3}$ | $8.39 \times 10^8$ | $2.44 \times 10^4$ | $1.45 \times 10^8$ | --- |
| J55 | $1.14 \times 10^{-4}$ | 0.999 | $1.28 \times 10^{-4}$ | $5.04 \times 10^{-4}$ | $2.63 \times 10^{-1}$ | $4.78 \times 10^{-4}$ | $3.09 \times 10^{-3}$ | $4.18 \times 10^6$ | $3.46 \times 10^3$ | $7.09 \times 10^4$ | --- |
| J56 | $1.14 \times 10^{-4}$ | 0.999 | $1.38 \times 10^{-4}$ | $4.75 \times 10^{-4}$ | $0.25 \times 10^{-2}$ | $1.80 \times 10^{-3}$ | $2.28 \times 10^{-3}$ | $1.48 \times 10^6$ | $3.33 \times 10^3$ | $4.97 \times 10^5$ | --- |
| J57 | $1.14 \times 10^{-4}$ | 0.999 | $9.94 \times 10^{-5}$ | $5.08 \times 10^{-4}$ | $2.64 \times 10^{-1}$ | $1.96 \times 10^{-4}$ | $4.29 \times 10^{-3}$ | $9.57 \times 10^4$ | $2.40 \times 10^3$ | $0.61 \times 10^3$ | --- |
| J73 | $1.14 \times 10^{-4}$ | 0.999 | $1.33 \times 10^{-4}$ | $4.79 \times 10^{-4}$ | $2.79 \times 10^{-1}$ | $6.55 \times 10^{-4}$ | $1.56 \times 10^{-3}$ | $3.81 \times 10^5$ | $2.48 \times 10^3$ | $1.12 \times 10^3$ | --- |
| J74 | $1.14 \times 10^{-4}$ | 0.999 | $1.35 \times 10^{-4}$ | $5.07 \times 10^{-4}$ | $4.22 \times 10^{-2}$ | $7.44 \times 10^{-4}$ | $3.91 \times 10^{-3}$ | $4.91 \times 10^7$ | $1.08 \times 10^4$ | $3.48 \times 10^6$ | --- |
| J75 | $1.14 \times 10^{-4}$ | 0.999 | $1.37 \times 10^{-4}$ | $4.83 \times 10^{-4}$ | $0.33 \times 10^{-2}$ | $1.11 \times 10^{-3}$ | $2.41 \times 10^{-3}$ | $1.22 \times 10^9$ | $3.55 \times 10^4$ | $1.23 \times 10^7$ | --- |
| J76 | $1.14 \times 10^{-4}$ | 0.999 | $1.36 \times 10^{-4}$ | $5.06 \times 10^{-4}$ | $1.29 \times 10^{-2}$ | $8.72 \times 10^{-4}$ | $4.30 \times 10^{-3}$ | $1.12 \times 10^9$ | $4.21 \times 10^4$ | $1.23 \times 10^7$ | --- |
| J82 | $1.14 \times 10^{-4}$ | 0.999 | $1.19 \times 10^{-4}$ | $5.00 \times 10^{-4}$ | $3.02 \times 10^{-1}$ | $3.43 \times 10^{-4}$ | $2.70 \times 10^{-3}$ | $1.33 \times 10^6$ | $1.36 \times 10^3$ | $1.78 \times 10^3$ | --- |
| J83 | $1.14 \times 10^{-4}$ | 0.999 | $1.09 \times 10^{-4}$ | $4.83 \times 10^{-4}$ | $3.62 \times 10^{-1}$ | $2.53 \times 10^{-4}$ | $1.69 \times 10^{-3}$ | $4.33 \times 10^5$ | $5.58 \times 10^3$ | $1.28 \times 10^3$ | --- |
| **PEG IFN-α and NAs treated patient (HBeAg-negative non-PVR)** |  |  |  |  |  |  |  |  |  |  |  |
| 4 | $1.14 \times 10^{-4}$ | 0.999 | $1.28 \times 10^{-4}$ | $4.82 \times 10^{-4}$ | $2.10 \times 10^{-1}$ | $4.73 \times 10^{-4}$ | $1.65 \times 10^{-3}$ | $2.41 \times 10^5$ | $0.55 \times 10^3$ | $2.21 \times 10^3$ | B |
| 8 | $1.14 \times 10^{-4}$ | 0.999 | $1.32 \times 10^{-4}$ | $5.12 \times 10^{-4}$ | $3.30 \times 10^{-1}$ | $6.22 \times 10^{-4}$ | $1.26 \times 10^{-2}$ | $1.33 \times 10^5$ | $0.31 \times 10^3$ | $4.01 \times 10^4$ | C |
| 10 | $1.14 \times 10^{-4}$ | 0.999 | $1.37 \times 10^{-4}$ | $5.11 \times 10^{-4}$ | $1.75 \times 10^{-1}$ | $1.12 \times 10^{-3}$ | $5.80 \times 10^{-3}$ | $6.44 \times 10^5$ | $6.43 \times 10^3$ | $1.96 \times 10^6$ | C |
| 13 | $1.14 \times 10^{-4}$ | 0.999 | $1.37 \times 10^{-4}$ | $5.02 \times 10^{-4}$ | $3.00 \times 10^{-1}$ | $1.10 \times 10^{-3}$ | $2.89 \times 10^{-3}$ | $6.30 \times 10^4$ | $5.94 \times 10^3$ | $1.08 \times 10^4$ | C |
| 16 | $1.14 \times 10^{-4}$ | 0.999 | $1.38 \times 10^{-4}$ | $5.00 \times 10^{-4}$ | $5.15 \times 10^{-1}$ | $1.13 \times 10^{-3}$ | $2.64 \times 10^{-3}$ | $6.20 \times 10^4$ | $2.66 \times 10^3$ | $4.18 \times 10^4$ | C |
| 17 | $1.14 \times 10^{-4}$ | 0.999 | $1.38 \times 10^{-4}$ | $5.12 \times 10^{-4}$ | $2.34 \times 10^{-1}$ | $2.30 \times 10^{-3}$ | $1.15 \times 10^{-2}$ | $6.55 \times 10^4$ | $0.77 \times 10^3$ | $1.34 \times 10^5$ | B |
| 19 | $1.14 \times 10^{-4}$ | 0.999 | $1.37 \times 10^{-4}$ | $5.10 \times 10^{-4}$ | $2.69 \times 10^{-1}$ | $1.10 \times 10^{-3}$ | $5.04 \times 10^{-3}$ | $1.29 \times 10^5$ | $2.58 \times 10^3$ | $2.89 \times 10^4$ | C |
| 21 | $1.14 \times 10^{-4}$ | 0.999 | $1.37 \times 10^{-4}$ | $5.06 \times 10^{-4}$ | $7.45 \times 10^{-2}$ | $1.11 \times 10^{-3}$ | $3.52 \times 10^{-3}$ | $2.03 \times 10^5$ | $6.47 \times 10^3$ | $1.94 \times 10^4$ | C |
| 27 | $1.14 \times 10^{-4}$ | 0.999 | $1.38 \times 10^{-4}$ | $5.11 \times 10^{-4}$ | $9.19 \times 10^{-2}$ | $1.38 \times 10^{-3}$ | $5.59 \times 10^{-3}$ | $6.43 \times 10^4$ | $2.06 \times 10^3$ | $3.72 \times 10^3$ | B |
| 28 | $1.14 \times 10^{-4}$ | 0.999 | $1.20 \times 10^{-4}$ | $5.11 \times 10^{-4}$ | $2.51 \times 10^{-1}$ | $3.49 \times 10^{-4}$ | $6.32 \times 10^{-3}$ | $3.14 \times 10^5$ | $2.59 \times 10^3$ | $1.23 \times 10^5$ | C |
| 33 | $1.14 \times 10^{-4}$ | 0.999 | $1.38 \times 10^{-4}$ | $5.10 \times 10^{-4}$ | $2.39 \times 10^{-1}$ | $1.27 \times 10^{-3}$ | $4.77 \times 10^{-3}$ | $4.95 \times 10^5$ | $3.54 \times 10^3$ | $1.26 \times 10^4$ | C |
| 35 | $1.14 \times 10^{-4}$ | 0.999 | $1.37 \times 10^{-4}$ | $5.04 \times 10^{-4}$ | $2.86 \times 10^{-1}$ | $4.19 \times 10^{-3}$ | $3.09 \times 10^{-3}$ | $2.04 \times 10^6$ | $3.47 \times 10^3$ | $9.16 \times 10^5$ | C |
| 36 | $1.14 \times 10^{-4}$ | 0.999 | $1.36 \times 10^{-4}$ | $5.10 \times 10^{-4}$ | $2.17 \times 10^{-1}$ | $1.19 \times 10^{-3}$ | $5.20 \times 10^{-3}$ | $1.46 \times 10^5$ | $1.52 \times 10^3$ | $2.38 \times 10^4$ | C |
| 39 | $1.14 \times 10^{-4}$ | 0.999 | $1.38 \times 10^{-4}$ | $5.11 \times 10^{-4}$ | $3.19 \times 10^{-1}$ | $1.41 \times 10^{-3}$ | $5.72 \times 10^{-3}$ | $1.56 \times 10^5$ | $8.92 \times 10^3$ | $3.11 \times 10^3$ | C |
| 40 | $1.14 \times 10^{-4}$ | 0.999 | $1.38 \times 10^{-4}$ | $5.03 \times 10^{-4}$ | $2.55 \times 10^{-1}$ | $2.21 \times 10^{-3}$ | $3.07 \times 10^{-3}$ | $1.19 \times 10^5$ | $2.01 \times 10^4$ | $5.23 \times 10^3$ | C |
| 42 | $1.14 \times 10^{-4}$ | 0.999 | $1.38 \times 10^{-4}$ | $5.04 \times 10^{-4}$ | $3.11 \times 10^{-1}$ | $2.36 \times 10^{-3}$ | $3.13 \times 10^{-3}$ | $5.55 \times 10^4$ | $8.11 \times 10^3$ | $2.08 \times 10^4$ | C |
| 44 | $1.14 \times 10^{-4}$ | 0.999 | $1.26 \times 10^{-4}$ | $5.12 \times 10^{-4}$ | $3.48 \times 10^{-1}$ | $4.39 \times 10^{-4}$ | $1.03 \times 10^{-2}$ | $1.23 \times 10^5$ | $0.13 \times 10^3$ | $1.17 \times 10^5$ | C |
| 46 | $1.14 \times 10^{-4}$ | 0.999 | $1.34 \times 10^{-4}$ | $5.10 \times 10^{-4}$ | $3.03 \times 10^{-1}$ | $7.37 \times 10^{-4}$ | $5.08 \times 10^{-3}$ | $2.04 \times 10^5$ | $1.13 \times 10^3$ | $6.59 \times 10^3$ | B |
| 47 | $1.14 \times 10^{-4}$ | 0.999 | $1.36 \times 10^{-4}$ | $5.06 \times 10^{-4}$ | $1.68 \times 10^{-1}$ | $9.69 \times 10^{-3}$ | $3.59 \times 10^{-3}$ | $1.16 \times 10^6$ | $4.80 \times 10^3$ | $2.93 \times 10^4$ | C |
| 50 | $1.14 \times 10^{-4}$ | 0.999 | $1.38 \times 10^{-4}$ | $5.10 \times 10^{-4}$ | $2.10 \times 10^{-1}$ | $3.06 \times 10^{-3}$ | $4.68 \times 10^{-3}$ | $9.60 \times 10^4$ | $7.71 \times 10^3$ | $1.48 \times 10^3$ | C |
| 57 | $1.14 \times 10^{-4}$ | 0.999 | $1.38 \times 10^{-4}$ | $5.03 \times 10^{-4}$ | $3.11 \times 10^{-1}$ | $3.40 \times 10^{-3}$ | $2.98 \times 10^{-3}$ | $8.98 \times 10^4$ | $4.87 \times 10^3$ | $1.15 \times 10^4$ | C |
| 59 | $1.14 \times 10^{-4}$ | 0.999 | $1.38 \times 10^{-4}$ | $5.05 \times 10^{-4}$ | $2.13 \times 10^{-1}$ | $3.69 \times 10^{-3}$ | $3.28 \times 10^{-3}$ | $1.03 \times 10^5$ | $5.80 \times 10^3$ | $2.20 \times 10^5$ | C |
| 61 | $1.14 \times 10^{-4}$ | 0.999 | $1.25 \times 10^{-4}$ | $5.00 \times 10^{-4}$ | $2.21 \times 10^{-1}$ | $4.25 \times 10^{-4}$ | $2.64 \times 10^{-3}$ | $3.92 \times 10^6$ | $1.41 \times 10^3$ | $2.67 \times 10^6$ | C |
| 62 | $1.14 \times 10^{-4}$ | 0.999 | $1.34 \times 10^{-4}$ | $5.08 \times 10^{-4}$ | $2.49 \times 10^{-1}$ | $7.30 \times 10^{-4}$ | $3.95 \times 10^{-3}$ | $2.45 \times 10^5$ | $3.93 \times 10^3$ | $2.70 \times 10^4$ | C |
| 66 | $1.14 \times 10^{-4}$ | 0.999 | $1.38 \times 10^{-4}$ | $5.11 \times 10^{-4}$ | $5.56 \times 10^{-1}$ | $2.95 \times 10^{-3}$ | $5.40 \times 10^{-3}$ | $1.98 \times 10^5$ | $7.88 \times 10^3$ | $1.81 \times 10^4$ | C |
| 69 | $1.14 \times 10^{-4}$ | 0.999 | $1.37 \times 10^{-4}$ | $5.06 \times 10^{-4}$ | $2.72 \times 10^{-1}$ | $9.95 \times 10^{-4}$ | $3.48 \times 10^{-3}$ | $6.76 \times 10^4$ | $2.28 \times 10^4$ | $3.28 \times 10^5$ | C |
| 70 | $1.14 \times 10^{-4}$ | 0.999 | $1.39 \times 10^{-4}$ | $5.04 \times 10^{-4}$ | $3.01 \times 10^{-1}$ | $3.40 \times 10^{-3}$ | $3.17 \times 10^{-3}$ | $9.09 \times 10^4$ | $4.96 \times 10^3$ | $8.42 \times 10^3$ | C |
| 73 | $1.14 \times 10^{-4}$ | 0.999 | $1.37 \times 10^{-4}$ | $5.00 \times 10^{-4}$ | $1.02 \times 10^{-1}$ | $3.27 \times 10^{-3}$ | $2.60 \times 10^{-3}$ | $1.17 \times 10^5$ | $4.12 \times 10^3$ | $1.76 \times 10^5$ | C |
| 80 | $1.14 \times 10^{-4}$ | 0.999 | $1.38 \times 10^{-4}$ | $5.08 \times 10^{-4}$ | $2.93 \times 10^{-1}$ | $2.19 \times 10^{-3}$ | $4.12 \times 10^{-3}$ | $5.73 \times 10^4$ | $1.18 \times 10^3$ | $2.81 \times 10^3$ | B |
| 81 | $1.14 \times 10^{-4}$ | 0.999 | $1.30 \times 10^{-4}$ | $5.12 \times 10^{-4}$ | $3.16 \times 10^{-1}$ | $5.44 \times 10^{-4}$ | $6.81 \times 10^{-3}$ | $6.22 \times 10^5$ | $0.65 \times 10^3$ | $4.29 \times 10^3$ | B |
| 82 | $1.14 \times 10^{-4}$ | 0.999 | $1.19 \times 10^{-4}$ | $5.08 \times 10^{-4}$ | $3.36 \times 10^{-1}$ | $3.36 \times 10^{-4}$ | $4.18 \times 10^{-3}$ | $2.27 \times 10^5$ | $1.19 \times 10^3$ | $1.66 \times 10^3$ | C |
| 85 | $1.14 \times 10^{-4}$ | 0.999 | $1.38 \times 10^{-4}$ | $5.10 \times 10^{-4}$ | $9.43 \times 10^{-2}$ | $1.25 \times 10^{-3}$ | $4.91 \times 10^{-3}$ | $5.23 \times 10^4$ | $0.28 \times 10^3$ | $4.53 \times 10^5$ | C |
| 91 | $1.14 \times 10^{-4}$ | 0.999 | $1.38 \times 10^{-4}$ | $4.99 \times 10^{-4}$ | $2.90 \times 10^{-1}$ | $2.03 \times 10^{-3}$ | $2.55 \times 10^{-3}$ | $4.25 \times 10^4$ | $4.10 \times 10^3$ | $0.25 \times 10^3$ | C |
| 93 | $1.14 \times 10^{-4}$ | 0.999 | $1.37 \times 10^{-4}$ | $5.05 \times 10^{-4}$ | $3.16 \times 10^{-1}$ | $9.58 \times 10^{-4}$ | $3.40 \times 10^{-3}$ | $6.88 \times 10^4$ | $1.14 \times 10^4$ | $2.55 \times 10^3$ | C |
| 94 | $1.14 \times 10^{-4}$ | 0.999 | $1.38 \times 10^{-4}$ | $4.96 \times 10^{-4}$ | $3.22 \times 10^{-1}$ | $1.31 \times 10^{-3}$ | $2.31 \times 10^{-3}$ | $8.06 \times 10^4$ | $3.38 \times 10^4$ | $1.63 \times 10^4$ | C |
| 97 | $1.14 \times 10^{-4}$ | 0.999 | $1.38 \times 10^{-4}$ | $5.06 \times 10^{-4}$ | $3.04 \times 10^{-1}$ | $4.10 \times 10^{-3}$ | $3.61 \times 10^{-3}$ | $3.87 \times 10^5$ | $5.13 \times 10^3$ | $7.34 \times 10^3$ | C |
| 106 | $1.14 \times 10^{-4}$ | 0.999 | $1.24 \times 10^{-4}$ | $5.12 \times 10^{-4}$ | $3.36 \times 10^{-1}$ | $4.14 \times 10^{-4}$ | $1.14 \times 10^{-2}$ | $1.30 \times 10^5$ | $3.07 \times 10^3$ | $1.05 \times 10^3$ | C |
| 107 | $1.14 \times 10^{-4}$ | 0.999 | $1.36 \times 10^{-4}$ | $4.94 \times 10^{-4}$ | $3.70 \times 10^{-2}$ | $8.48 \times 10^{-4}$ | $2.18 \times 10^{-3}$ | $2.14 \times 10^4$ | $1.05 \times 10^4$ | $1.20 \times 10^3$ | C |
| 111 | $1.14 \times 10^{-4}$ | 0.999 | $1.37 \times 10^{-4}$ | $5.05 \times 10^{-4}$ | $3.28 \times 10^{-1}$ | $1.12 \times 10^{-3}$ | $3.34 \times 10^{-3}$ | $8.83 \times 10^4$ | $1.92 \times 10^3$ | $0.26 \times 10^3$ | B |
| 113 | $1.14 \times 10^{-4}$ | 0.999 | $1.36 \times 10^{-4}$ | $4.93 \times 10^{-4}$ | $2.32 \times 10^{-1}$ | $7.63 \times 10^{-3}$ | $2.14 \times 10^{-3}$ | $8.62 \times 10^5$ | $1.18 \times 10^4$ | $1.98 \times 10^4$ | C |
| 115 | $1.14 \times 10^{-4}$ | 0.999 | $1.37 \times 10^{-4}$ | $4.96 \times 10^{-4}$ | $9.53 \times 10^{-2}$ | $4.69 \times 10^{-3}$ | $2.29 \times 10^{-3}$ | $1.69 \times 10^5$ | $1.62 \times 10^3$ | $4.35 \times 10^5$ | C |
| 116 | $1.14 \times 10^{-4}$ | 0.999 | $1.38 \times 10^{-4}$ | $5.00 \times 10^{-4}$ | $1.79 \times 10^{-1}$ | $2.05 \times 10^{-3}$ | $2.61 \times 10^{-3}$ | $4.32 \times 10^4$ | $8.78 \times 10^3$ | $3.03 \times 10^3$ | C |
| 118 | $1.14 \times 10^{-4}$ | 0.999 | $1.33 \times 10^{-4}$ | $4.94 \times 10^{-4}$ | $2.09 \times 10^{-1}$ | $6.31 \times 10^{-4}$ | $2.16 \times 10^{-3}$ | $9.38 \times 10^4$ | $9.50 \times 10^3$ | $9.83 \times 10^3$ | C |
| 120 | $1.14 \times 10^{-4}$ | 0.999 | $1.38 \times 10^{-4}$ | $5.05 \times 10^{-4}$ | $2.74 \times 10^{-1}$ | $1.40 \times 10^{-3}$ | $3.38 \times 10^{-3}$ | $9.20 \times 10^4$ | $1.37 \times 10^3$ | $1.76 \times 10^5$ | C |
| 123 | $1.14 \times 10^{-4}$ | 0.999 | $1.68 \times 10^{-4}$ | $5.13 \times 10^{-4}$ | $2.57 \times 10^{-1}$ | $4.16 \times 10^{-3}$ | $1.02 \times 10^{-2}$ | $4.92 \times 10^6$ | $1.07 \times 10^4$ | $2.05 \times 10^6$ | C |
| NAs (ETV or LAM) |  |  |  |  |  |  |  |  |  |  |  |
| E01 | $1.14 \times 10^{-4}$ | 0.999 | $1.37 \times 10^{-4}$ | - | $3.22 \times 10^{-2}$ | $1.21 \times 10^{-3}$ | - | $4.84 \times 10^3$ | $1.57 \times 10^1$ | $4.88 \times 10^5$ | --- |
| E02 | $1.14 \times 10^{-4}$ | 0.999 | $1.36 \times 10^{-4}$ | - | $1.29 \times 10^{-2}$ | $4.62 \times 10^{-3}$ | - | $2.62 \times 10^5$ | $6.42 \times 10^4$ | $2.76 \times 10^8$ | --- |
| E03 | $1.14 \times 10^{-4}$ | 0.999 | $1.38 \times 10^{-4}$ | - | $8.95 \times 10^{-2}$ | $1.51 \times 10^{-3}$ | - | $2.05 \times 10^6$ | $2.68 \times 10^3$ | $2.32 \times 10^5$ | --- |
| E04 | $1.14 \times 10^{-4}$ | 0.999 | $1.30 \times 10^{-4}$ | - | $3.06 \times 10^{-1}$ | $5.35 \times 10^{-4}$ | - | $3.45 \times 10^4$ | $0.95 \times 10^3$ | $0.89 \times 10^3$ | --- |
| E05 | $1.14 \times 10^{-4}$ | 0.999 | $1.38 \times 10^{-4}$ | - | $2.77 \times 10^{-2}$ | $1.33 \times 10^{-3}$ | - | $6.55 \times 10^4$ | $0.28 \times 10^3$ | $3.49 \times 10^4$ | --- |
| E06 | $1.14 \times 10^{-4}$ | 0.999 | $1.36 \times 10^{-4}$ | - | $2.13 \times 10^{-1}$ | $4.00 \times 10^{-3}$ | - | $2.44 \times 10^7$ | $0.32 \times 10^3$ | $2.80 \times 10^4$ | --- |
| E07 | $1.14 \times 10^{-4}$ | 0.999 | $1.38 \times 10^{-4}$ | - | $1.73 \times 10^{-1}$ | $3.65 \times 10^{-3}$ | - | $1.23 \times 10^7$ | $2.35 \times 10^3$ | $1.31 \times 10^4$ | --- |
| E08 | $1.14 \times 10^{-4}$ | 0.999 | $1.36 \times 10^{-4}$ | - | $1.60 \times 10^{-1}$ | $8.76 \times 10^{-4}$ | - | $1.79 \times 10^6$ | $0.35 \times 10^3$ | $1.32 \times 10^4$ | --- |
| E09 | $1.14 \times 10^{-4}$ | 0.999 | $1.09 \times 10^{-4}$ | - | $1.36 \times 10^{-1}$ | $2.49 \times 10^{-4}$ | - | $5.89 \times 10^6$ | $2.70 \times 10^3$ | $1.23 \times 10^3$ | --- |
| L01 | $1.14 \times 10^{-4}$ | 0.999 | $1.26 \times 10^{-4}$ | - | $8.75 \times 10^{-2}$ | $4.51 \times 10^{-4}$ | - | $5.37 \times 10^5$ | $0.53 \times 10^3$ | $6.96 \times 10^5$ | --- |
| L02 | $1.14 \times 10^{-4}$ | 0.999 | $1.18 \times 10^{-4}$ | - | $2.57 \times 10^{-2}$ | $3.26 \times 10^{-4}$ | - | $1.10 \times 10^6$ | $2.99 \times 10^3$ | $1.39 \times 10^3$ | --- |
| L03 | $1.14 \times 10^{-4}$ | 0.999 | $1.38 \times 10^{-4}$ | - | $2.65 \times 10^{-1}$ | $2.23 \times 10^{-3}$ | - | $1.02 \times 10^5$ | $4.45 \times 10^1$ | $0.76 \times 10^3$ | --- |
| L04 | $1.14 \times 10^{-4}$ | 0.999 | $1.16 \times 10^{-4}$ | - | $4.68 \times 10^{-1}$ | $1.91 \times 10^{-3}$ | - | $2.82 \times 10^6$ | $0.20 \times 10^3$ | $2.25 \times 10^3$ | --- |
| L05 | $1.14 \times 10^{-4}$ | 0.999 | $1.08 \times 10^{-4}$ | - | $1.20 \times 10^{-1}$ | $3.08 \times 10^{-4}$ | - | $1.91 \times 10^6$ | $1.44 \times 10^3$ | $2.78 \times 10^6$ | --- |
| L06 | $1.14 \times 10^{-4}$ | 0.999 | $1.31 \times 10^{-4}$ | - | $3.63 \times 10^{-2}$ | $2.44 \times 10^{-4}$ | - | $1.97 \times 10^5$ | $5.10 \times 10^3$ | $2.48 \times 10^5$ | --- |
| L07 | $1.14 \times 10^{-4}$ | 0.999 | $1.27 \times 10^{-4}$ | - | $1.39 \times 10^{-1}$ | $5.71 \times 10^{-4}$ | - | $7.40 \times 10^5$ | $3.74 \times 10^3$ | $7.31 \times 10^4$ | --- |
| L08 | $1.14 \times 10^{-4}$ | 0.999 | $1.36 \times 10^{-4}$ | - | $1.53 \times 10^{-1}$ | $4.61 \times 10^{-4}$ | - | $6.01 \times 10^5$ | $6.04 \times 10^3$ | $1.24 \times 10^5$ | --- |
| L09-1 | $1.14 \times 10^{-4}$ | 0.999 | $1.24 \times 10^{-4}$ | - | $9.05 \times 10^{-2}$ | $2.75 \times 10^{-3}$ | - | $1.25 \times 10^7$ | $1.42 \times 10^3$ | $1.58 \times 10^5$ | --- |
| L09-2 | $1.14 \times 10^{-4}$ | 0.999 | $1.38 \times 10^{-4}$ | - | $1.99 \times 10^{-2}$ | $4.14 \times 10^{-4}$ | - | $1.37 \times 10^4$ | $0.44 \times 10^3$ | $3.25 \times 10^4$ | --- |
| L10-1 | $1.14 \times 10^{-4}$ | 0.999 | $1.38 \times 10^{-4}$ | - | $7.04 \times 10^{-2}$ | $8.73 \times 10^{-4}$ | - | $2.43 \times 10^7$ | $1.92 \times 10^3$ | $2.37 \times 10^3$ | --- |
| L10-2 | $1.14 \times 10^{-4}$ | 0.999 | $1.38 \times 10^{-4}$ | - | $6.24 \times 10^{-2}$ | $1.93 \times 10^{-3}$ | - | $2.48 \times 10^4$ | $0.55 \times 10^3$ | $2.01 \times 10^3$ | --- |
| L11 | $1.14 \times 10^{-4}$ | 0.999 | $1.38 \times 10^{-4}$ | - | $6.81 \times 10^{-2}$ | $1.58 \times 10^{-3}$ | - | $4.52 \times 10^6$ | $1.66 \times 10^3$ | $1.90 \times 10^5$ | --- |
| L12 | $1.14 \times 10^{-4}$ | 0.999 | $1.26 \times 10^{-4}$ | - | $6.79 \times 10^{-1}$ | $4.37 \times 10^{-4}$ | - | $8.33 \times 10^5$ | $0.27 \times 10^3$ | $2.11 \times 10^3$ | --- |
| L13 | $1.14 \times 10^{-4}$ | 0.999 | $1.36 \times 10^{-4}$ | - | $6.93 \times 10^{-1}$ | $1.10 \times 10^{-2}$ | - | $2.69 \times 10^8$ | $0.14 \times 10^3$ | $4.93 \times 10^5$ | --- |
| L14 | $1.14 \times 10^{-4}$ | 0.999 | $1.08 \times 10^{-4}$ | - | $1.00 \times 10^{-1}$ | $2.47 \times 10^{-4}$ | - | $3.28 \times 10^5$ | $8.79 \times 10^3$ | $3.90 \times 10^4$ | --- |
| L15 | $1.14 \times 10^{-4}$ | 0.999 | $1.38 \times 10^{-4}$ | - | $6.79 \times 10^{-2}$ | $2.23 \times 10^{-3}$ | - | $3.17 \times 10^5$ | $3.24 \times 10^3$ | $1.93 \times 10^6$ | --- |
| L16 | $1.14 \times 10^{-4}$ | 0.999 | $1.37 \times 10^{-4}$ | - | $2.01 \times 10^{-2}$ | $1.41 \times 10^{-3}$ | - | $7.45 \times 10^4$ | $1.12 \times 10^4$ | $1.55 \times 10^7$ | --- |

<sup>†</sup> Production rate of HBV DNA from cccDNA × Fraction of HBV DNA recycling for cccDNA.

### Supplementary Note 1: Modeling intracellular HBV replication in primary human hepatocytes

To describe the intracellular virus life cycle in HBV-infected primary human hepatocytes, we developed the following mathematical model:

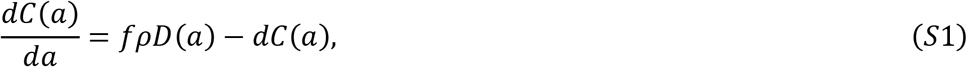

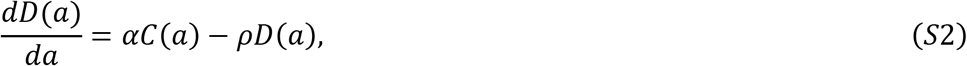

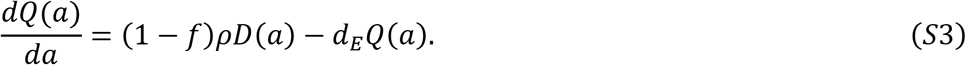

The variables *C*(*a*), *D*(*a*) and *Q*(*a*) represent the amount of intracellular cccDNA and intracellular and extracellular HBV DNA in cultures that have been infected for time *a* (i.e., *a* is considered as an infection age), respectively. The intracellular HBV DNA is produced from cccDNA at rate *α* and is lost at rate *ρ* of which a fraction 1 – *f* of HBV DNA is assembled with viral proteins as virus particles that are exported out of infected cells, and the other fraction *f* is reused for further cccDNA formation. The viral particles have a degradation rate *d_E_* and cccDNA has a degradation rate of *d*. We have ignored the degradation of intracellular DNA since it is small compared with the consumption rate of HBV DNA due to virion production2,3 (see **Table S1**). This intracellular HBV replication model can be modified to include the antiviral effects of different classes of drugs. For example, under treatment with entecavir (ETV), which is a reverse transcriptase inhibitor, the antiviral effect of ETV is assumed to be in blocking HBV DNA production with an effectiveness, *ε*, 0 < *ε* ≤ 1, and is modelled by assuming

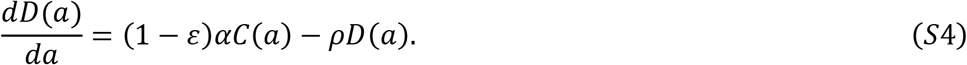

In addition, to predict unknown but possible mechanisms of action of cytokines and estimate their antiviral effect in promoting cccDNA degradation, *ε_d_*, inhibiting HBV DNA production, *ε_α_*, or inhibiting viral release, *ε_f_*, we further expand the mathematical model assuming these hypothetical mechanisms of action:

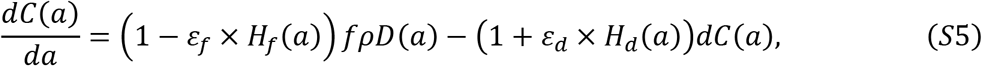

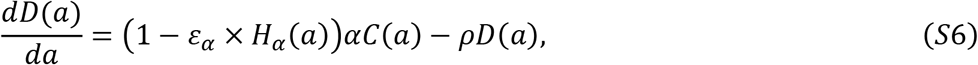

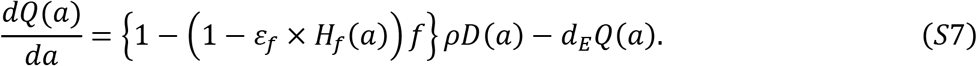

Here *H*(*a*) is a Heaviside step function defined as *H*(*t*) = 0 if *a* > *τ_d_, τ_α_, τ_f_*: otherwise *H*(*a*) = 1, where *τ_d_, τ_α_, τ_f_* are the times the cytokine effects end for promoting cccDNA degradation, inhibiting HBV DNA production, and inhibiting viral releasing, respectively. Note that, in our data fitting, to predict the “major” mechanism of action of each cytokine, we separately assumed each of the three antiviral effects and estimated its corresponding ε.

### Supplementary Note 2: Transformation to a system of ODEs from a PDE multiscale model

We here introduce a multiscale model using partial differential equations (PDEs) that couple intra-, inter- and extra-cellular virus dynamics for analyzing multiscale experimental data of HBV infection (c.f.^4^) (**Fig. 3A**):

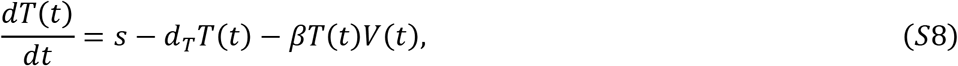

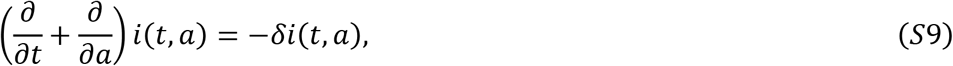

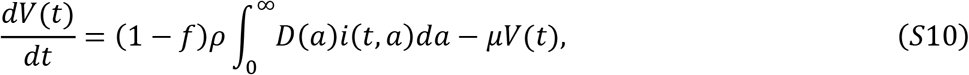

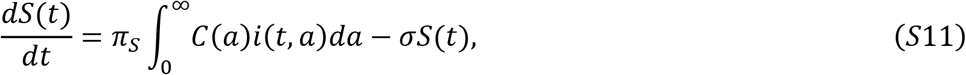

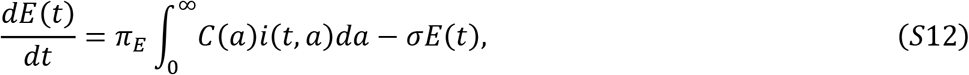

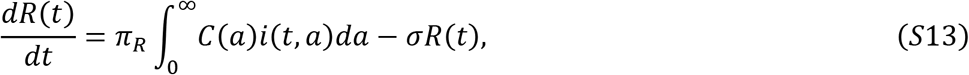

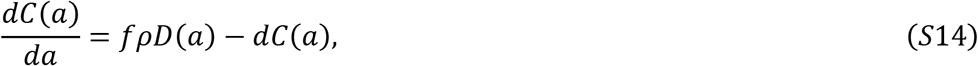

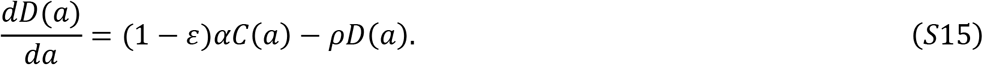

with the boundary condition *i*(*t*, 0) = *βT*(*t*)*V*(*t*) and initial condition *i*(0, *a*) = *i*_0_(*a*). The intercellular variables *T*(*t*) and *V*(*t*) are the number of uninfected cells and the (extracellular) HBV DNA load, respectively. We defined the density of infected cells with infection age *a* as *i*(*t, a*), and therefore the total number of infected cells is 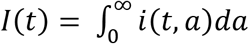. The intracellular variables *C*(*a*) and *D*(*a*), which evolve depending on the age *a*, represent the amount of intracellular cccDNA and HBV DNA, respectively. We also defined extracellular variables used as “surrogate biomarkers” to predict the dynamics of cccDNA in hepatocytes, that is, the amount of HBsAg, HBeAg and HBcrAg antigens as *S*(*t*), *E*(*t*) and *R*(*t*), respectively. The definition of an age-structured population model is found in^5^.

In addition to the intracellular HBV replication dynamics (see **Supplementary Note 1**), we assumed target cells, *T*, are supplied at rate *s*, die at per capita rate *d_T_*, are infected by viruses at rate *β*, and the infected cells die at per capita rate *δ*. We also assumed that HBsAg, HBeAg and HBcrAg antigens are produced from cccDNA in infected cells at rates *π_S_, π_E_* and *π_R_*, and are cleared at rate *σ*, respectively. The exported viral particles, i.e., extracellular HBV DNA load, is assumed to be cleared at rate *μ* per virion.

Since Eqs. (*S*14-*S*15) are a set of linear ordinary differential equations (ODEs), we directly solved them and obtained the following analytical solutions:

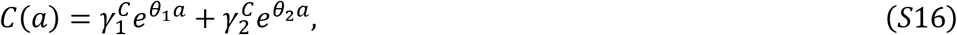

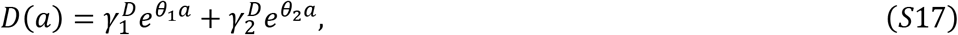

where

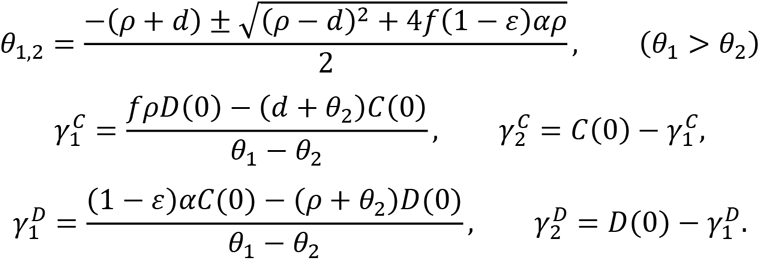

As we recently reported, 6,7 the multiscale PDE model, Eqs. (*S*8-*S*15), can be transformed into a mathematically identical set of ordinary differential equations as follows. Using the method of characteristics with initial and boundary conditions of *i*(*t, a*), we transform Eq. (*S*9) into

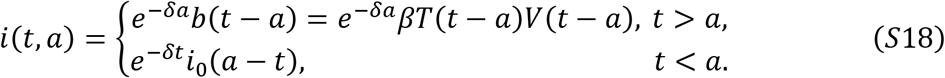

Then, *I*(*t*) is evaluated as follows:

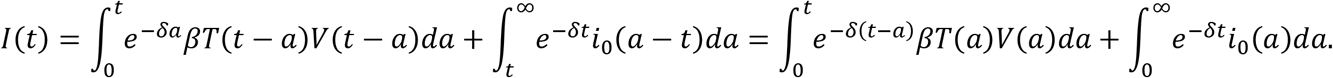

Since 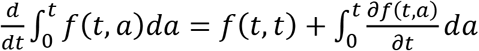, differentiating *I*(*t*) with respect to time *t*, we obtain the following ODE:

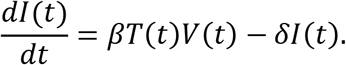

In addition, inserting Eq. (*S*17-18) into Eq. (*S*10), we have

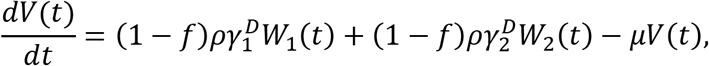

where the variables *W*_1_(*t*) and *W*_2_(*t*) are defined as

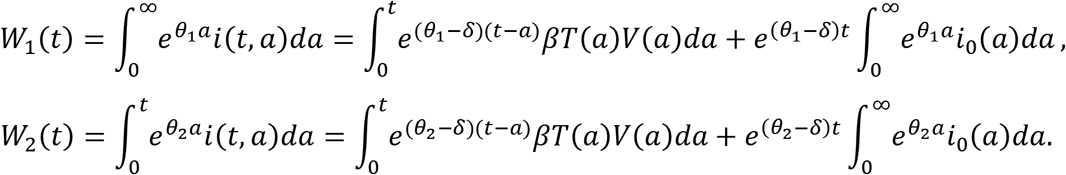

We obtain the following ODEs for *W*_1_(*t*) and *W*_2_(*t*):

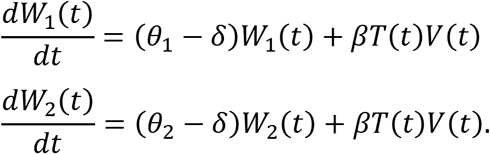

In similar manner, inserting Eq. (*S*16-*S*18) into Eqs. (*S*11-*S*13), we have the corresponding ODEs. Therefore, the multiscale PDE model is described as the following equivalent system of ODEs:

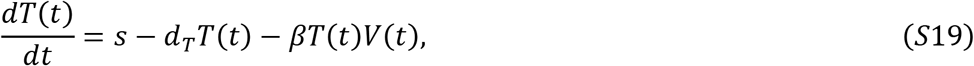

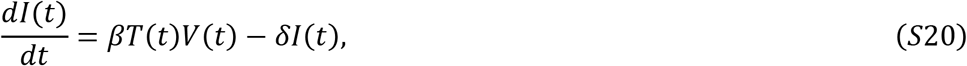

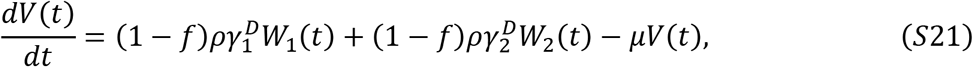

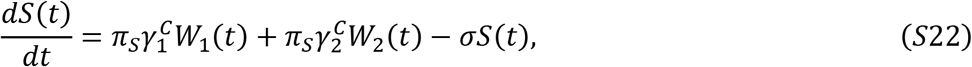

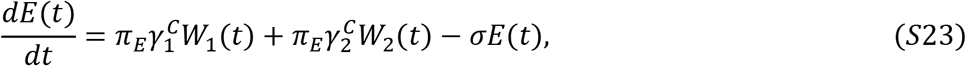

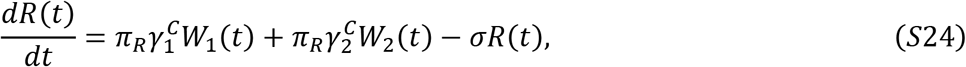

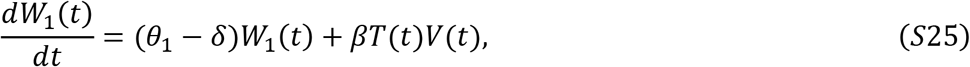

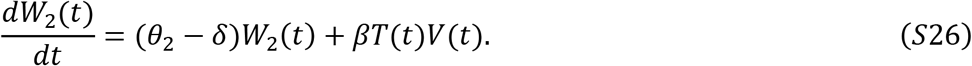

Note that Eqs. (*S*19-*S*26) will be further simplified for the purpose of data analysis depending on the antiviral treatment assumed (see later).

### Supplementary Note 3: Linearized equations under potent NAs treatment *in vivo*

We assumed that NAs treatment is potent enough that intracellular HBV replications and *de novo* infections are negligible after treatment initiation^8–11^, i.e., the antiviral effectiveness of NAs on intracellular HBV replications is assumed to be 0 < *ε* ≤ 1 and

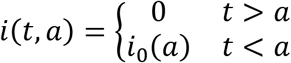

Then Eqs. (*S*19-*S*26) can be simplified as follows:

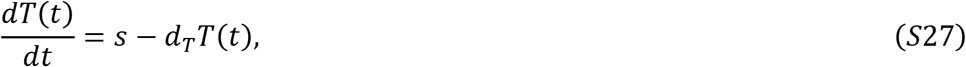

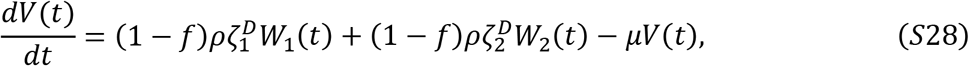

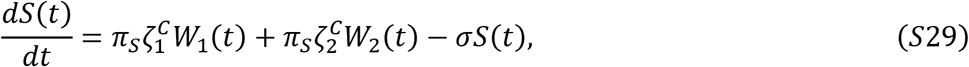

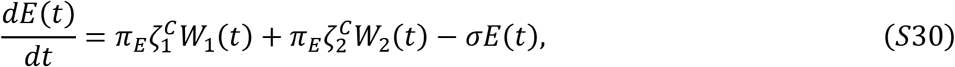

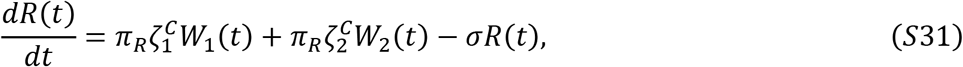

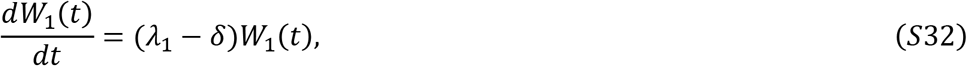

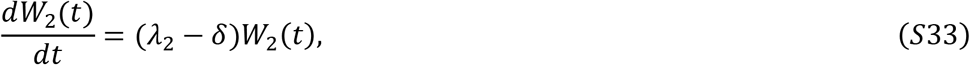

where, 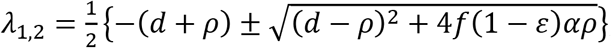, 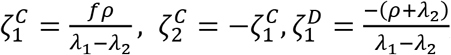 and 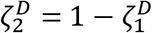. We also consider all variables in Eqs. (*S*19-*S*26) are in steady state before treatment initiation^12^, and particularly that the infected cells obtain a stable age distribution, i.e., *i*_0_(*a*) = *βT*(0)*V*(0)*e*^−*δa*^.

Since Eqs. (*S*27-*S*33) are a set of linear ODEs, we directly solve them, and find the following analytical solutions:

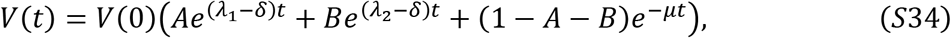

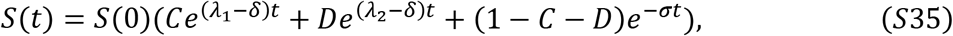

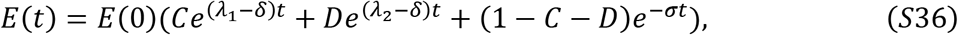

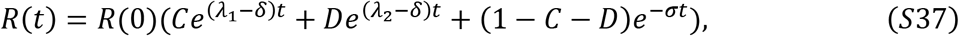

where 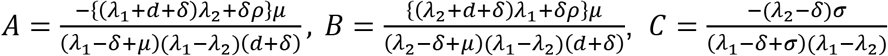 and 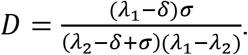.

### Supplementary Note 4: Linearized equations under potent PEG IFN-α treatment *in vivo*

We also assumed that PEG IFN-α treatment is potent enough that intracellular HBV replication and *de novo* infections are negligible after treatment initiation^2,9,10,13,14^ (**Fig. 1C**), i.e., the antiviral effect of PEG IFN-α on intracellular HBV replications is assumed to be 0 < *ε* ≤ 1 and

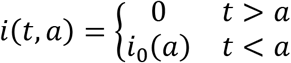

Then Eqs. (*S*19-*S*26) can be simplified to

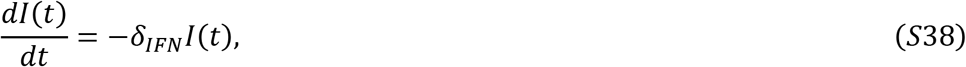

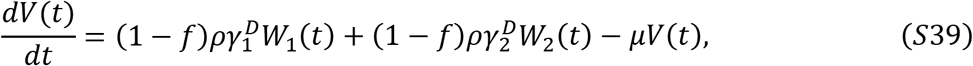

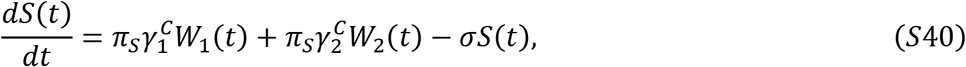

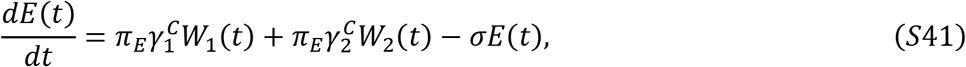

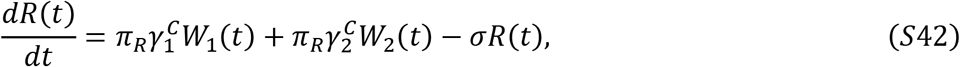

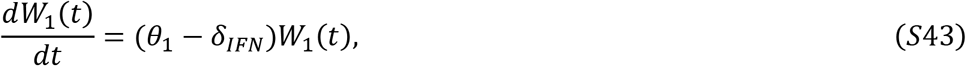

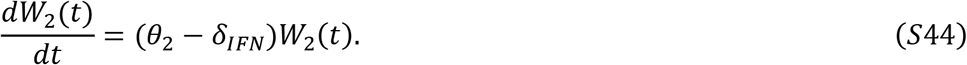

In addition, it has been reported that PEG IFN-α induces interferon-stimulated genes (ISGs) and ISGs potentially degrade intracellular cccDNA. Therefore, we assumed PEG IFN-α increases the cccDNA degradation rate^15^, i.e., *d_IFN_* (> *d*). Similarly, we assume all variables in Eqs. (*S*19-*S*26) are in steady state before treatment initiation, and that the infected cells have obtained a stable age distribution, i.e., *i*_0_(*α*) = *βT*(0)*V*(0)*e*^−*δa*^. As shown in **Fig. *S*3**, because PEG IFN-α enhances the decay rate of infected cells in HBV infection in humanized mouse due to cytotoxic effects (but relatively mild), we assumed *δ_IFN_* (> *δ*) in the data fitting (**Fig. 2BC**). Solving Eqs. (*S*38-*S*44) we find

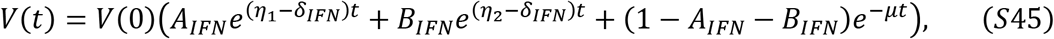

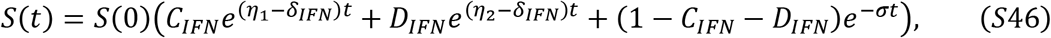

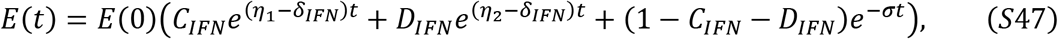

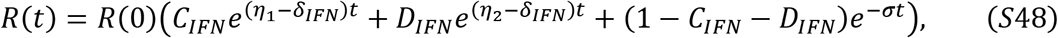

moreover, the total amount of cccDNA and the amount of cccDNA per infected cell are derived from 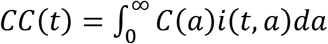 and *C*(*t*) = *CC*(*t*)/*I*(*t*) as follows

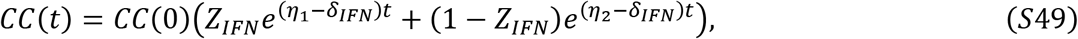

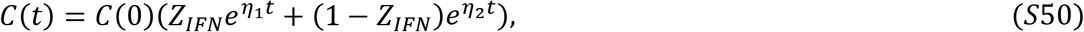

where 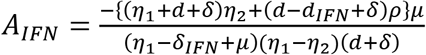, 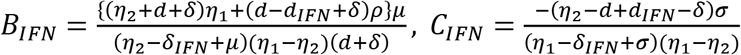, 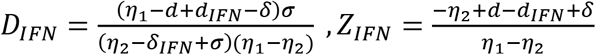 and 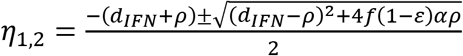.

### Supplementary Note 5: Data fitting and parameter estimation

#### (1) Data analysis for HBV infection on PHH

We categorized datasets as follows: [condition 1 = No ETV treatment], [condition 2 = ETV treatment from day 1] and [condition 3 = ETV treatment from day 10] (**Fig.S1A**). To assess the variability of kinetic parameters and model predictions, we performed Bayesian inference for the dataset of condition 1, 2 and 3 using Markov chain Monte Carlo (MCMC) sampling^16^. A statistical model adopted from Bayesian inference assumed that measurement error followed a normal distribution with mean zero and constant variance (error variance). Simultaneously, we fitted Eqs. (*S*1–*S*3) and Eqs. (*S*1–*S*2)(*S*4) to the experimental data of intracellular HBV DNA and cccDNA, and extracellular HBV DNA in condition 1 and conditions 2, 3, respectively (**Fig.1B**). Note that we estimated model parameters (i.e., *α, f, d, α, d_E_, ε*) for all conditions as common values because the HBV used in this assay is identical. On the other hand, susceptibility and permissiveness of PHH to HBV are known as heterogeneity; thus, we used different initial values (i.e., *C*(0), *D*(0), *Q*(0)) for each condition (**Table S1**). Distributions of model parameters and initial values were inferred directly by MCMC computations^16^.

We also categorized datasets as follows; [condition 4 = ETV+IFN-α treatment from day 1 and 10] and [condition 5 = IFN-α treatment from day 1 and 10] (**Fig.S1A**). To evaluate the mechanism of action of ETV, we first estimated *α, f, d, ρ, d_E_, τ_i_, ε_i_* and *C_i_*(0), *D_i_*(0), *Q_i_*(0) (*i* = *d, α, f*) by fitting Eqs. (S5–S7) to the experimental data in conditions 1, 2 and 3 simultaneously using nonlinear least squares regression (**Fig.S2** and **Table S2**), and confirmed that calculating the sum of squared residuals (SSR) and selecting the mathematical model with the smallest SSR was able to successfully predict the known mechanism of action of ETV (**Fig.1C**). Then, fixing estimated parameter values for *α, f, d, ρ* and *d_E_*, we further estimated *τ_i_*, *ε_i_* and *C_i_*(0), *d_i_*(0), *Q_i_*(0) (*i* = *d, α, f*) for ETV+IFN-α and IFN-α treatment by fitting Eqs. (*S*5–*S*7) to the experimental data in conditions 4 and 5, respectively (**Fig.S2** and **Table S2**). The SSR for data fitting by mathematical models assuming hypothetical mechanisms of action of cytokines are summarized in **Fig.1C**.

#### (2) Data analysis for HBV infection on humanized mouse

To quantify HBV infection and the antiviral effect of ETV or IFN-α in humanized mice, we also performed Bayesian inference using MCMC sampling because the inter-individual variations are almost negligible. We here used a previously estimated half-life of extracellular HBV DNA in peripheral blood (PB), that is, 62 minutes (*μ* = 16.1 d^-1^)^17^, and that of extracellular HBsAg in PB, 0.69 day (*σ* = 1 d^-1^)^18^. Simultaneously, we fitted Eqs. (*S*34-*S*37) and Eqs. (*S*45-*S*48) to the experimentally measured extracellular HBV DNA, HBcrAg, HBeAg and HBsAg obtained from HBV-infected humanized mice treated with ETV and PEG IFN-α, respectively (**Fig. 2BC**), and estimated *d, d_IFN_* and *ρ* (**Table S3**).

Note that we fixed all initial values as initial points of our dataset (**Table S4**), and the decay rates of infected cells were separately estimated from h-Alb in PB of the humanized mice (**Fig.S3** and **Table S3**).

#### (3) Data analysis for PEG IFN-α or ETV/LAM treated HBV patients

MONOLIX 2019R2 (www.lixoft.com), a program for maximum likelihood estimation for a nonlinear mixed-effects model, was employed to fit the model, Eqs. (*S*45-*S*46)(*S*48), to extracellular HBV DNA, HBcrAg and HBsAg in patients PB receiving PEG IFN-α monotherapy or PEG IFN-α combination with ETV/LAM (**Fig. S4**). In addition, we fit the model, Eqs. (*S*34-*S*35)(*S*37), to extracellular HBV DNA, HBcrAg and HBsAg in patients PB receiving NAs (**Fig. S4**). We assumed that the clearance rates of extracellular HBV DNA and antigens were *μ* = 0.57819 d^-1 20^ and *σ* = 0.13919 d^-1 10^ as previously estimated, respectively. Nonlinear mixed-effects modelling approaches incorporate a fixed effect as well as a random effect describing the inter-patient variability in parameters. Including a random effect amounts to a partial pooling of the data between individuals to improve estimates of the parameters applicable across the population of cases. By using this approach, the differences between the above 3 different biomarkers in PB in different individuals were not estimated explicitly, nor did we fully pool the data which would bias estimates towards highly sampled cases. In this method of estimation, each parameter estimate (*ϑ_i_* = *ϑ* × *e*^*π_i_*^) depends on the individual where *ϑ* is fixed effect, and *π_i_* is random effect with an assumed Gaussian distribution with mean 0 and standard deviation *Ω*. Population parameters and individual parameters were estimated using the stochastic approximation expectation-approximation algorithm^21^ and empirical Bayes’ method^22^, respectively. We divided our datasets into five groups; [PEG IFN-α treated HBeAg positive patients achieving VR], [PEG IFN-α treated HBeAg positive patients showing non-VR], [PEG IFN-α treated HBeAg negative patients achieving PVR], [PEG IFN-α HBeAg negative treated patients showing non-PVR] and [ETV/LAM treated patients]. Estimated population parameters, initial values, and their interpatient variability are listed in **Table S6**. Using estimated parameters, goodness of fit was also assessed based on individual predictions and the measured HBV DNA, HBcrAg and HBsAg for all patients (see **Fig. S5**).

### Supplementary Note 6: Detection limit for HBV DNA, HBsAg and cccDNA

In the Asian-Pacific clinical practice guidelines on the management of hepatitis B and previous papers, the detection limits of HBV markers were described as HBV DNA <12 (IU/ml)^23–25^ and HBsAg <0.05 (IU/ml)^26–28^. On the other hand, Caviglia et al. constructed a highly sensitive method using droplet digital PCR with a lower limit of detection of 0.8 × 10^-5^ copies/cell for quantitation of cccDNA in the liver of HBV-infected patients^29^. According to these reports, we evaluated predicted PEG IFN-α treatment periods for achieving these detection limits. Note that we here cannot directly evaluate “HBV cure” as recently defined in^30^, because our clinical datasets do not include integrated HBV DNA and hepatitis B surface antibody (anti-HBs).

